# i^6^A-seq maps *N*^6^-isopentenyladenosine and uncovers its role as a regulator of mRNA stability through recruitment of DIS3L2

**DOI:** 10.1101/2025.11.26.690722

**Authors:** C. Avrahami-Rozenblum, L. Cohen Ben-Aderet, R. Ashwal-Fluss, R. Masoud, O. Nayshool, T. Zinger, M. Sevilla-Sharon, E. Glick-Saar, A. Yahalom, L. Coll-SanMartin, M. Esteller, G. Rechavi, D. Dominissini, S. Moshitch-Moshkovitz

## Abstract

Dynamic post-transcriptional RNA modifications are crucial regulators of RNA metabolism and cell fate. Recent advances in sequencing, combined with antibody, enzymatic, or chemical approaches, have enabled transcriptome-wide mapping of these modifications. While prevalent in tRNAs and rRNAs, a growing number of modifications also adorn lower-abundance mRNA transcripts. Global mapping efforts, particularly for *N*^6^-methyladenosine (m^6^A), uncovered the affected mechanisms governing RNA metabolism, shedding light on novel, modification-dependent, modes of gene regulation.

*N*^6^-isopentenyladenosine (i^6^A) is a conserved tRNA modification involved in translation fidelity and efficiency. i^6^A depletion is linked to mitochondrial defects and human diseases. Current i^6^A detection methods are low-throughput, and its lack of base-pairing effects makes sequencing-based identification challenging.

Here we developed a novel and robust mapping technique, i^6^A-seq, a method that utilizes antibody-mediated enrichment and iodine chemical labeling to generate a reverse transcription signature, enabling transcriptome-wide i^6^A detection. Our global mapping revealed hundreds of i^6^A sites in human and mouse mRNA, indicating its presence in this RNA species, with conserved isopentenylome features. These sites exhibit a typical consensus sequence, primarily in coding transcripts (CDS), preferentially within lysine codons. Consistent with its role in tRNA, tRNA-isopentenyltransferase (TRIT1) also appears to install i^6^A in mRNA. Manipulation of TRIT1 revealed that i^6^A regulates the expression of a subset of genes through mRNA decay, specifically those isopentenylated at the CDS and translated on ER-bound ribosomes. Importantly, DIS3 like 3’-5’ exoribonuclease 2 (DIS3L2), an RNA exoribonuclease known to regulate ER-translated mRNA, was identified as the first i^6^A reader protein.

Our findings introduce i^6^A as a new modified nucleotide that decorates mRNA, allowing us to decipher its regulatory roles in gene expression. This study not only establishes the presence and location of i^6^A in mRNA, but also uncovers its first functional mechanism. Similar to the knowledge accumulated on m^6^A, this work paves the way for further discoveries with potential relevance for understanding gene regulation, disease diagnosis, and therapy.

## Introduction

Fine regulation of gene expression requires coordinated multi-layered control to achieve precise controlled spatiotemporal output. Beyond well studied epigenetic mechanisms that involve chemical modifications of DNA and histones, mRNA modifications, collectively known as the epitranscriptome, have emerged as a critical regulatory layer. The epitranscriptome repertoire is continually expanding, comprising at least 11 distinct mRNA modifications to date^1^, among which *N*^6^-methyladenosine (m^6^A) is the most prevalent and extensively characterized^2^. m^6^A is a dynamic regulator of gene expression, influencing central mechanisms of mRNA metabolism including splicing, stability, translation, and nuclear export^3^. m^6^A metabolism is governed by writers (methyltransferases), erasers (demethylases) and readers (m^6^A-dependent RNA-binding proteins). m^6^A readers are at the heart of gene regulation, recruiting downstream cellular machineries to exert context-dependent biological functions^4^. Other mRNA modifications also regulate gene expression, but their research, compared to m^6^A is still lagging.

RNA modifications of tRNAs are equally essential in regulating translation efficiency and accuracy^5^. The *N*^6^-isopentenyladenosine (i^6^A) modification at position 37 (i^6^A37) exemplifies this regulatory layer by reinforcing the anti-codon loop in a U-turn configuration, and stabilizing the codon-anticodon pairing fidelity^6–8^. tRNA i^6^A is particularly important in U/A rich codons that are inherently weaker and more error prone^9^. It is installed by the enzyme tRNA-isopentenyltransferase (TRIT1), which targets specific cytosolic tRNAs such as tRNA^Ser^ (with anticodons UCU, UCC, UCG, UCA, IGA) and the selenocysteine specific tRNA^Sec^ (anticodon UCA)^6,10^. In mitochondrial tRNAs i^6^A37 can be further modified to ms^2^i^6^A by the Cdk5 regulatory subunit associated protein 1 (CDK5RAP1) enzyme^11^. Notably, i^6^A37 in tRNA^[Ser]Sec^ is indispensable for the translation of selenoproteins, enabling the recoding of the UGA canonical stop codon into selenocysteine^12^.

Pathogenic mutations in TRIT1 in humans are associated with neurodevelopmental delay, myoclonic epilepsy, progressive spasticity, and impaired mitochondrial function^13^. still, the comprehensive understanding of the role TRIT1 in diseases remains unclear. In cancer TRIT1 displays dysregulation in both directions: while it is downregulated across a broad range of tumors, marking it as a potential tumor suppressor^14^, some malignancies express very high levels of TRIT1^15^.

Despite the significance of i^6^A, its global mapping was hindered by technical limitations. Earlier detection methods of i^6^A relied on a combination of chemical, biochemical and analytical approaches^16,17^. A new approach harnessed the unique reactivity of the isopentenyl (prenyl) group of i^6^A with iodine at mild buffer conditions, to develop an unbiased method for global i^6^A mapping. The new method, Iodine Mediated cyclization and Reverse Transcription (IMCRT) relies on the bulkiness of the iodination product to interfere with reverse transcription (RT), resulting in a typical RT signature of misincorporations at the i^6^A site^18^. IMCRT successfully identified all 9 i^6^A37-containging tRNAs of *Saccharomyces cerevisiae*^18^. Although this tool opens new options in epitranscriptomics, its use for transcriptome-wide mapping in complex samples, such as mRNA or total RNA, is still limited and requires further development.

The rapid development of the field of epitranscriptomics has sparked several controversies over the biological relevance of low-abundance and rare mRNA modifications, arguing that these reflect a non-specific activity of tRNA or RNA writer enzymes. However, functional studies uncovered that these modifications play important roles in regulation of gene expression. Among these are *N*^1^-methyadenosine (m^1^A)^19–21^, dihydrouridine^22^, 1-methyguanosine^23^ and more^1^.

In this work, we uncovered the presence of i^6^A in human and mouse mRNA, and developed a new tool, i^6^A-seq, for its global mapping in highly diverse samples. Similar to IMCRT^18^, our new method takes advantage of the unique chemical properties of i^6^A and its reactivity with iodine. To identify i^6^A sites in mRNA, we combined iodine-labeling reaction with antibody-based enrichment and dedicated bioinformatic analysis, resulting in a defined i^6^A-specific RT signature that identifies this modification at a single-base resolution.

In search of a functional role for i^6^A in mRNA, we discovered an inverse correlation between TRIT1 expression and transcript stability of several i^6^A-modified transcripts, suggesting a role in regulation of transcript degradation. By harnessing RNA affinity chromatography, we discovered that DIS3 like 3’-5’ exoribonuclease 2 (DIS3L2) binds and degrades i^6^A-modified targets, revealing a new regulatory mechanism. Our findings expand the landscape of mRNA modifications and reveal i⁶A as a new regulator of transcript stability.

## Materials and methods

### Cells and media

Human Hela, HepG2 and HEK293 cell lines were maintained in DMEM (Gibco), and human DMS273 cell line was maintained in Waymouth’s medium (Gibco). Culture medium was supplemented with 10% FBS and penicillin/streptomycin. Mouse tissues were obtained from wild-type C57BL/6 mice. mESCs were expanded in IMDM (Biological Industries) supplemented with 20% FBS, 1% L-glutamin, non-essential amino acids, and penicillin/streptomycin, 0.7µM β-mercaptoethanol, and 10µg recombinant human LIF (PeproTech), and maintained in 20% O_2_ conditions on irradiation inactivated mouse embryonic fibroblast feeder cells.

### Plasmids and transfections

Plasmids coding WT or mutant FLAG-tagged human TRIT1 (Origene, RC210476) were transfected into HEK293 and DMS273-shTRIT1 cells using TransIT-X2 reagent (Mirus Bio), according to manufacturer instructions, and harvested after 48 hours. **siRNAs** including scramble control or siRNA directed against DIS3L2 (Origene, SR315069) were transfected twice into DMS273-Scramble cells using RNAiMAX reagent (Invitrogen), according to manufacturer instructions, every 48 hours. Transfected cells were lastly harvested after 96 hours.

### RNA purification

Total RNA of human and mouse cells and tissues were purified using Direct-zol™ RNA Miniprep Plus (Zymo). To avoid DNA contamination, all samples were treated with DNase. Separating total RNA into small (17-200 nucleotides) and large (>200 nucleotides) transcript populations were performed using RNA Clean & Concentrator-25 (Zymo). Enrichment of polyadenylated RNA (poly(A)+ RNA) from total RNA was performed using GenElute mRNA miniprep kit (Sigma-Aldrich). RNA samples were chemically fragmented into ~60-nucleotide-long fragments using RNA Fragmentation Reagents (Invitrogen) for 15 min at 70 °C. RNA integrity and size validation was assessed using RNA High Sensitivity ScreenTape Assay (Agilent).

### i^6^A enrichment

IP of i^6^A in small RNA or mRNA is based on previously described m^6^A- and m^1^A-seq protocols^3,24^, with the following modifications: Anti-i^6^A antibody (Agrisera) was pre-coupled to Protein G Dynabeads (Invitrogen) and used to immunoprecipitate RNA for 3 hours at 4°C in IPP buffer (150 mM NaCl, 0.1% NP-40, 10 mM Tris-HCl, pH 7.4). After extensive washing, bound RNA was eluted from the beads by digestion with Proteinase K (NEB) for 1.5 hours at 37 °C, followed by TRIzol (Invitrogen) extraction of the supernatant and RNA cleanup using RNA Clean & Concentrator-5 (Zymo).

### Iodine labeling

Denatured RNA was labeled with 0.2M iodine (Sigma-Aldrich) for 1 hour at 37°C. RNA was then cleaned using RNA Clean & Concentrator-5 (Zymo) and used for library preparation as described below.

### Massive parallel sequencing

RNA libraries were prepared using either NEXTflex Small RNA-Seq Kit, NEBNext Ultra II Directional RNA Library Prep Kit or as previously described for the library construction protocol for m^1^A-MAP^25^. Sequencing was carried out using the Illumina NovaSeq 6000 platform and Illumina kits (NovaSeq 6000 SP Reagent Kit v1.5, 20040719). Samples were sequenced in a 300 pair-end manner.

**Direct RNA sequencing** of DMS273 cells was carried out on GridIon using Direct RNA Sequencing kits (SQK-RNA004).

### Bioinformatic analysis

i⁶A discovery was performed by integrating mutation-based and truncation-based signals from paired I₂-treated and untreated RNA-seq BAM files. The pipeline identifies A→T-associated mutation hotspots and 5′-read truncations on sense strands, merges and filters these events by coverage and non-reference frequency, and outputs annotated consensus sites representing candidate i⁶A modification positions.

### i^6^A detection and quantitation

Purified RNA was subjected to LC-MS/MS for detection and accurate quantitation of i^6^A. 300ng of purified RNA, or i^6^A-enriched RNA, was digested by P1 nuclease as described^26^. Detection and analysis of the digested sample was separated by ultra-performance liquid chromatography (Vanquish UHPLC Thermo Fisher scientific) on Hyprsil GOLD aQ column (Thermo Scientific 25303-152130), and then detected by Q Exactive™ HF hybrid quadrupole-Orbitrap mass spectrometer (Thermo Scientific) in the Parallel Reaction Monitoring Mode (PRM).

#### Site directed mutagenesis

TRIT1 mutations p.R323Q^27^ and p.E327K^28^ were introduced, separately, into a purchased human TRIT1 plasmid (Origene, RC210476) using the Q5 Site-Directed Mutagenesis Kit (NEB). Primers used are detailed in Table 1.

**Table 1.**
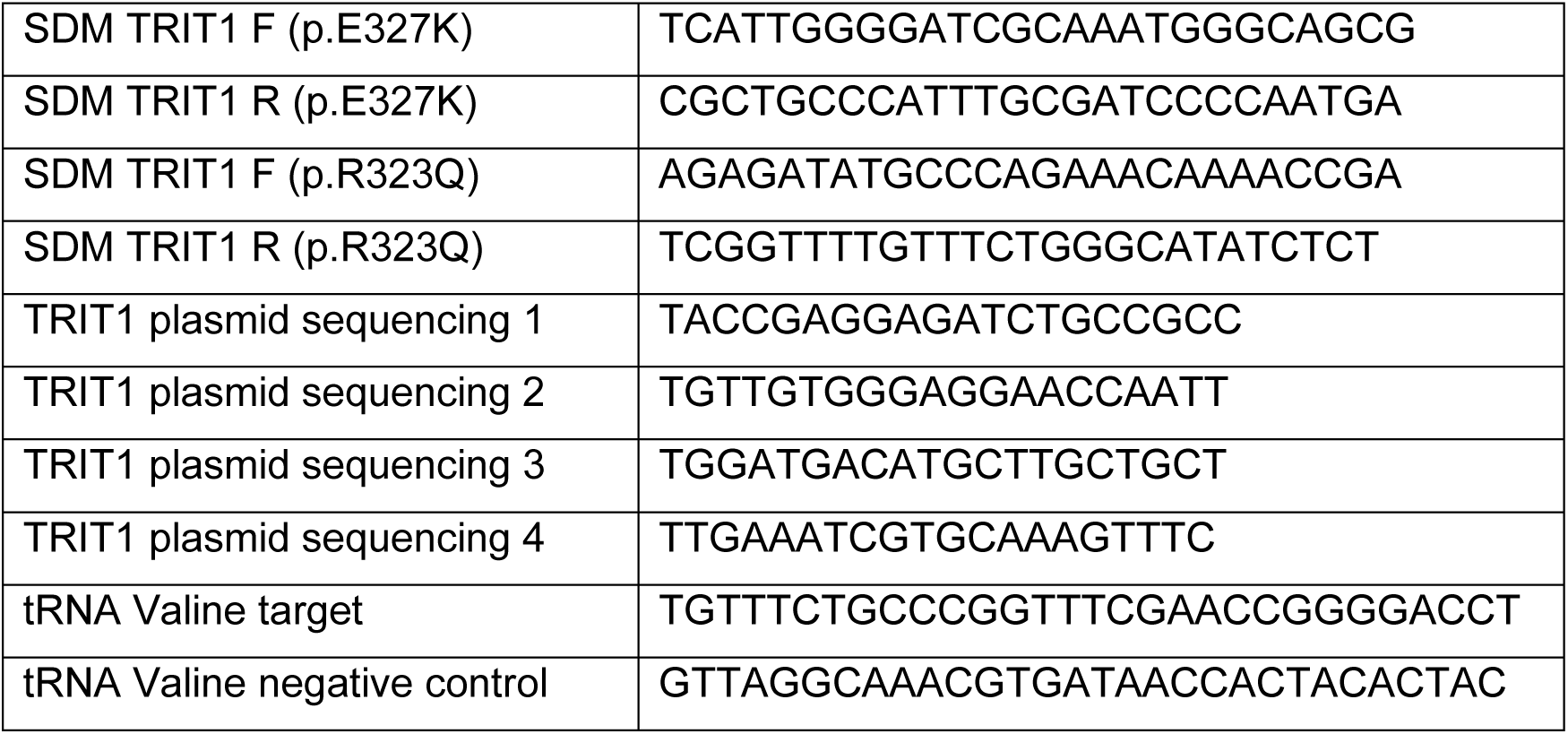
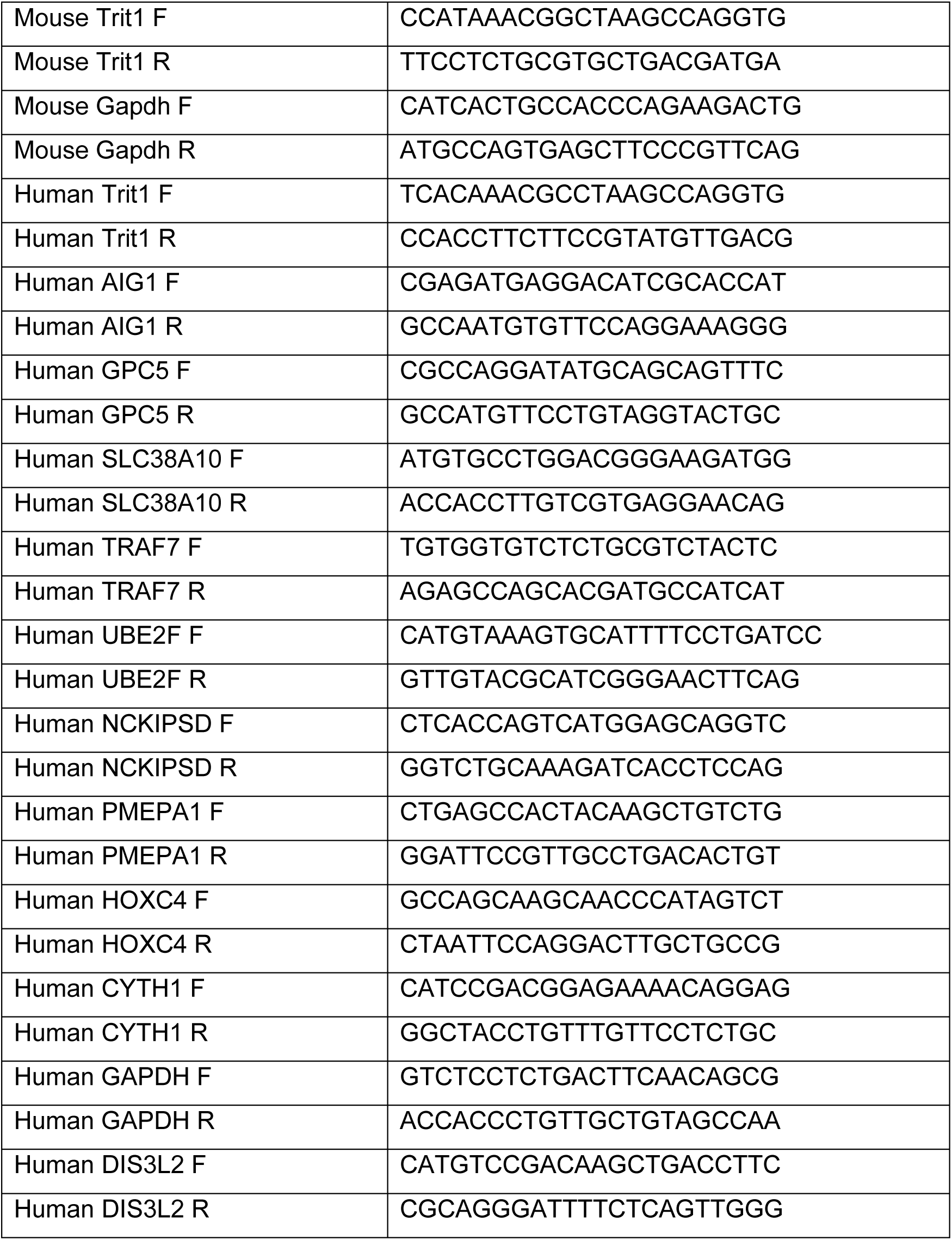
DNA Primers used in this study.

##### Immunostaining

Cells were fixated using 4% formaldehyde and permeabilized in 0.1% Triton. Blocking was done in 1% BSA. Cells were then incubated with the appropriate primary antibody (anti-i^6^A antibody Agrisera 1:250, anti-SLC38A10 antibody Novus 1:50, anti-AIG1 antibody Abcam 1:50) for 2 hours at room temperature, followed by incubation with a secondary antibody (Invitrogen) for 1 hour. Lastly, cells were incubated for 5 minutes with DAPI (Sigma-Aldrich 1:10,000). Analysis was carried out by a Leica confocal microscope.

#### Real-time qPCR

Isolated RNA was reverse transcribed into cDNA using High-Capacity cDNA Reverse Transcription Kit (Invitrogen) and analyzed in triplicates for each sample. Samples were normalized to the housekeeping gene GAPDH. Primers used are detailed in Table 1.

##### Western blot

Samples were separated on 4-12% Bis-Tris Plus gels (Invitrogen) and transferred onto nitrocellulose membrane using iBlot gel transfer system set to P0 for 8 min with iBlot gel transfer stacks (Invitrogen). Membranes were blocked in 5% BSA, 0.05% Tween-20 in PBS for 1 hour. The appropriate primary antibody was then incubated overnight at 4°C. Secondary antibodies were HRP-linked appropriate secondary antibodies (Cell Signaling). Blots were developed using the SuperSignal West Pico Luminol/Enhancer solution. The following primary antibodies were used: TRIT1 (Proteintech TA345223), GAPDH (BioRad 12004168), SLC38A10 (Novus NBP1-81193), AIG1 (Abcam ab140186), DIS3L2 (Novus NBP2-38264).

#### *in vitro* isopentenylation

RNA oligonucleotides were incubated with recombinant TRIT1 and DMAPP (Sigma-Aldrich) for 4 hours at 37°C. The enzymatic reaction was terminated by TRIzol (Invitrogen), followed by RNA isolation and RNA desalting by gel filtration. On-bead *in vitro* isopentenylation was similarly performed, with WT or mutant TRIT1 proteins isolated following transfection in HEK293 cells and immunoprecipitated using anti-Flag beads (Merck). Products were analyzed by HPLC and sequencing. RNA oligonucleotides used are detailed in Table 2. **Nuclease protection assay:** RNA isolated from HEK293 cells was incubated with DNA oligonucleotide for 5 min at 90°C and cooled slowly to 45°C. The resultant hybrids were digested using Mung bean nuclease (NEB) and RNase A (Sigma-Aldrich). The fragments were separated on denaturing Urea 15% PAGE gel and then extracted. The purified hybrids were digested using P1 nuclease (Sigma-Aldrich) and subjected to MS. Primers used are detailed in Table 1.

**Table 2.**
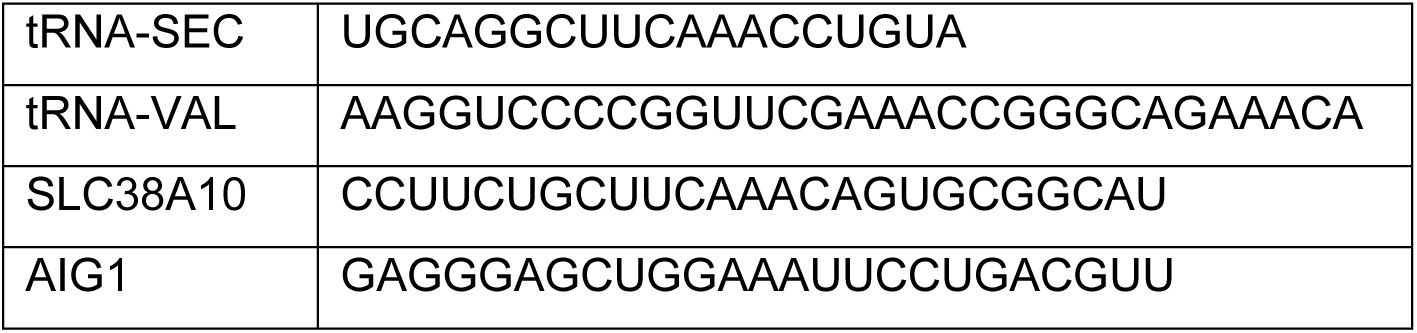
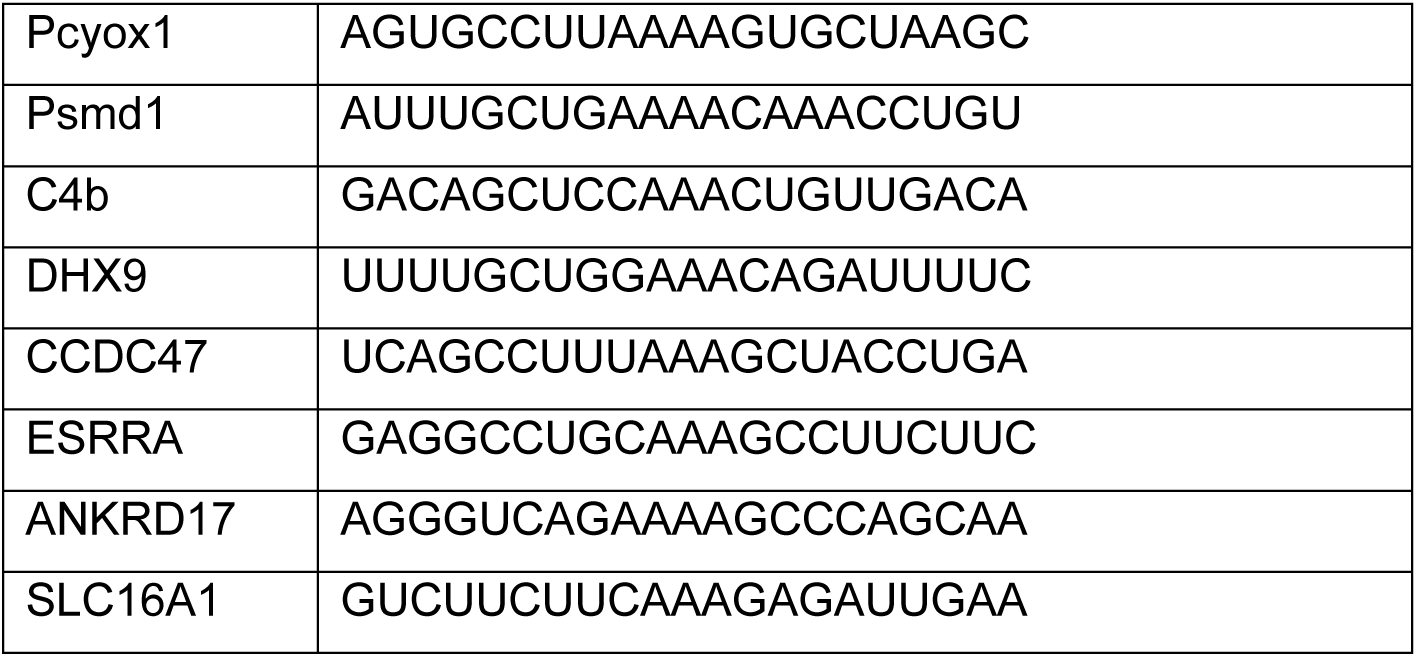
RNA oligonucleotides used for validation and RAC.

##### RNA stability

Each cell type was plated and incubated for 24 hours. Medium was replaced to fresh medium containing 5μM actinomycin-D (Sigma-Aldrich) for inhibition of mRNA transcription. Cells were harvested at the indicated time points and total RNA was extracted and analyzed by real-time qPCR. mRNA half-life was calculated as described in Chen et al. Methods Enzymol, 2008^29^.

#### RNA affinity chromatography

Performed as described in Edupuganti et al. Nat Struct Mol Biol, 2017^30^. For each gene, biotin-labelled RNA oligonucleotide bait (detailed in Table 2) spanning 25 nucleotides centered on a characterized i^6^A site was incubated for in vitro isopentenylation. Bound proteins were identified by either WB, or LC-MS/MS.

#### RIP

Flag-tagged control and DIS3L2 plasmid were transfected into HEK293 cells and conjugated to flag-tag beads after lysis (0.5% Triton (Aigma-Aldrich) in UPW). Protein-bound beads were then washed with PBS and incubated with 80μg of isolated mRNA from control DMS273 cells for 4 hours at 4°C. Bound RNA was eluted from the beads by digestion with Proteinase K (NEB) for 1.5 hours at 37 °C, followed by TRIzol (Invitrogen) extraction of the supernatant and RNA cleanup using RNA Clean & Concentrator-5 (Zymo). Analysis was performed by real-time qPCR.

## Results

### 1. i^6^A is present in mRNA

i^6^A is well documented, at position 37 in a set of tRNAs, where it maintains decoding fidelity and reading-frame preservation^7–9^. Whether i^6^A is also present in mRNA has not been established. To address this gap in our knowledge, we first tested the affinity of an anti-i^6^A antibody for i^6^A enrichment. Dot blot of synthetic RNA oligonucleotide with i^6^A, showed a gradual increase in signal at increasing amounts, and did not detect any signal in the unmodified or methylated RNA oligonucleotides (**Fig. 1a**). We also performed liquid chromatography coupled to tandem mass spectrometry (LC-MS/MS) analyses of yeast tRNA samples before and after immunoprecipitation (IP) with anti-i^6^A antibodies and observed a 9-fold enrichment of i^6^A, but not of other modifications (**Fig. S1a**). i^6^A was detectable in human mRNA only after antibody enrichment (**Fig. S1b**), providing the first evidence for its presence in this RNA species in human. Although measuring i^6^A in mRNA after antibody-based enrichment carries risks as bias due to cross-reactivity or mRNA impurity, i^6^A was the sole modification enriched, supporting its specific detection. These findings suggest that i^6^A may be a new epitranscriptomic mark, raising the need for a tool to globally map its sites in mRNA.

**Figure 1.**
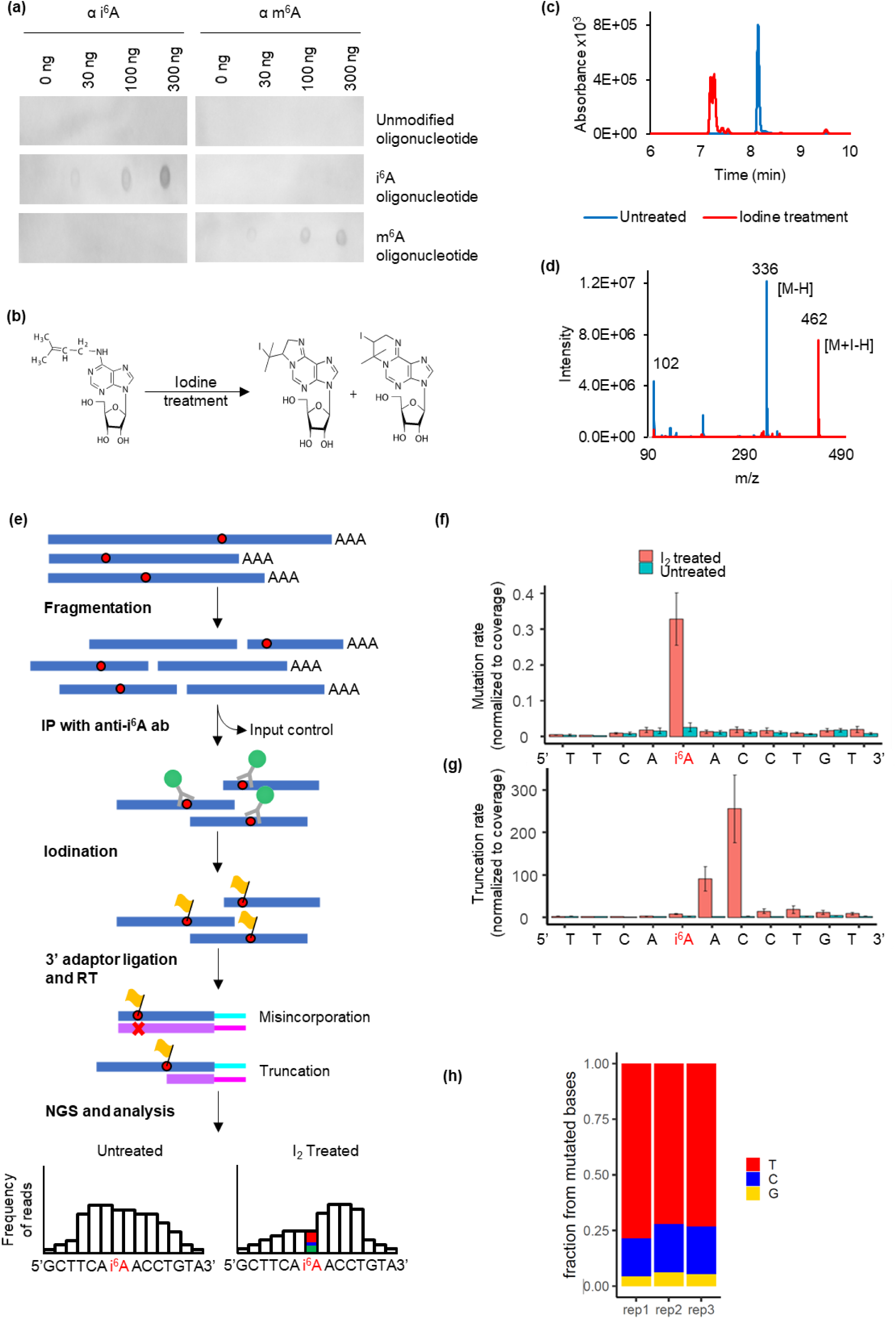
Development of i^6^A-seq. **(a)** Dot blots demonstrating high anti-i^6^A antibody specificity. Increasing amounts of synthetic RNA oligonucleotides containing i^6^A or m^6^A residues were spotted onto a membrane, and immunodetected with the indicated antibody. i^6^A, but not m^6^A, was detected only by anti-i^6^A antibody, reflecting minimal cross-reactivity. **(b)** Enzymatic labeling of i^6^A with iodine creates two hydrophilic products. **(c)** HPLC retention times of i^6^A before (untreated) and after (treated) iodine labeling. **(d)** LC-MS/MS of treated and untreated i^6^A nucleoside. Iodine addition to i^6^A created a molecule of greater molecular weight by 126 m/z. **(e)** Schematic diagram of the i^6^A-seq protocol. ab, antibody; IP, immunoprecipitation; RT, reverse transcription; NGS, next generation sequencing. **(f)** Elevated nucleotide misincorporation rate at the i^6^A position in an RNA oligonucleotide treated with iodine (red) compared to control untreated (blue), *n*=3 library replicates. **(g)** Truncation rate at the i^6^A site of the same RNA oligonucleotide shown in (f), with and without iodine treatment, *n*=3 library replicates. **(h)** Misincorporation profile at the i^6^A position after iodine treatment, revealing a non-random pattern.

### 2. i^6^A-seq is a new method to map i^6^A in mRNA

Similar to its impact on translation when present at position 37 of tRNA^6,7,9^, the presence of i^6^A in mRNA has the potential to influence biological processes such as RNA translation and structure. To gain insight into possible biological functions of i^6^A in mRNA, it is imperative to first identify its positions along the transcripts. We reason that a dedicated tool should overcome two main challenges: the low abundance in mRNA and the fact that in spite of its bulkiness and hydrophobic nature it does not leave an RT signature^4^. To address these challenges, we developed a new approach, i^6^A-seq that combines immuno-enrichment (IP) by antibodies with targeted chemical labeling of i^6^A (using a framework similar to IMCRT-tRNA-seq^18^), which impairs RT proofreading and processivity. The chemical reaction was validated by treating i^6^A nucleoside with iodine, creating two possible products (**Fig. 1b**). The iodine reacts as a nucleophile, attacking the double bond on the i^6^A, thereby closing either a five- or a six-member ring to make two possible hydrophilic products. Validated by reverse phase LC-MS, the two products had a shorter retention time due to their hydrophilic nature (**Fig. 1c**) and a higher molecular weight by MS (**Fig. 1d**). The i^6^A-seq protocol compares four sequencing libraries per mRNA sample: (1) untreated input; (2) iodine treated input; (3) untreated IP; (4) iodine treated IP. This design identifies i^6^A-specific RT signatures by enrichment (IP vs. input) and confirming iodine-dependency by comparison to untreated controls (**Fig. 1e**). To enhance the RT signatures, we leveraged the sensitivity of TGIRT^31^ at sub-optimal conditions to yield higher mutation and premature termination (PTM) levels in the cDNA.

### 3. Validation of i^6^A-seq

To establish i^6^A-seq as a mapping tool for mRNA application, we performed rigorous validations. First, we ensured the chemical specificity of the iodination treatment by subjecting 35 different RNA modifications to iodine labeling. The reaction products were analyzed by HPLC, confirming exclusive reactivity of i^6^A and its downstream modification, ms^2^i^6^A, under the reaction conditions (**Supplementary data file 1**). The next imperative step was to test the positional precision. We designed a synthetic RNA oligonucleotide derived from the tRNA^Sec^ sequence, which was i^6^A-modified at the known position, and subjected it to i^6^A-seq. Sequencing analysis revealed a high rate of misincorporation exclusively at the i^6^A position, only in iodine-treated oligonucleotide (**Fig. 1f**). Analysis revealed a non-random misincorporation profile (**Fig. 1h**). We also assessed the occurrence of premature terminations (PMTs) with respect to the modified position. PMTs in the cDNA, occurred 2-3 nucleotides upstream to the point where the RTase tackled the iodinated i^6^A position (**Fig 1g**). These PMTs occur only in an iodinated i^6^A-dependent manner. The final step involved defining the RT signature through integration of the i^6^A-dependent mutation profile and PMT landscape.

### 4. Testing i^6^A-seq on small RNA

To validate i^6^A-seq we applied it to small RNAs (<200 nts) from HEK293 and HAP1 human cell lines, focusing on tRNA with established i^6^A sites. In mice, i^6^A-seq was carried out on small RNA from brain and liver tissues. i^6^A-seq identified all known human and mouse i^6^A tRNA sites (**Fig. 2a**).

**Figure 2.**
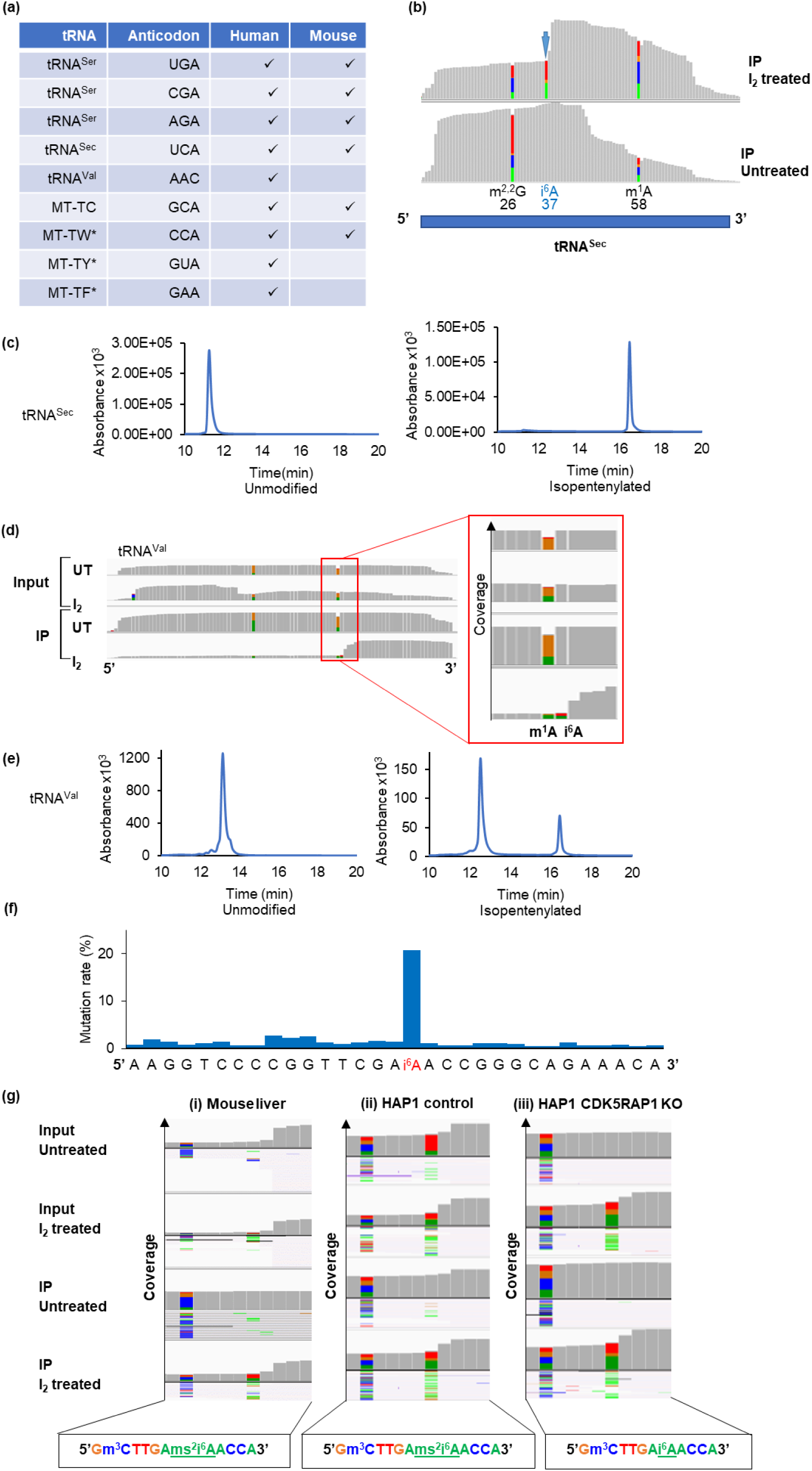
i^6^A in tRNA can be identified using i^6^A-seq. **(a)** Table of all i^6^A sites in tRNAs that were identified by i^6^A-seq in human and mouse. **(b)** IGV plots of tRNA^Sec^ IP samples in iodine-treated (top) and untreated (bottom) samples, indicating the misincorporation profile and truncation pattern at the known i^6^A position. Presence of two additional modifications is indicated, along with their known locations. These additional modifications were not affected by iodine treatment. **(c)** HPLC graphs of an RNA oligonucleotide derived from the tRNA^Sec^ before and after *in vitro* isopentenylation by TRIT1 enzyme. **(d)** IGV plots of tRNA^Val^ before and after iodine labeling, in input and IP samples, indicating the mutation profile and truncation pattern at the identified i^6^A position and the location of the adjacent m^1^A modification. **(e)** HPLC graphs of an RNA oligonucleotide derived from the tRNA^Val^ before and after *in vitro* isopentenylation by TRIT1 enzyme. **(f)** Sequencing of synthetic tRNA^Val^ oligonucleotide with i^6^A after iodine treatment, showing elevated mutation rates at the modification position. **(g)** IGV plots of mitochondrial tRNA^Ser^ in mouse liver tissue (right), HAP1 control cells (middle), and in HAP1 CDK5RAP1 KO cells (left), showing that the mutation profile of ms^2^i^6^A is independent of iodine treatment.

Analysis of tRNA^Sec^ sequencing, which contains additional RNA modifiactions, showed that other modifications such as m^1^A and m^2,2^G exhibit enhanced mutation rates regardless of iodine treatment (**Fig. 2b**), further validating i^6^A-seq as a reliable tool for global mapping of i^6^A in mRNA at base precision. The drop in coverage and mismatch accumulation on i^6^A position only in response to iodine treatment are depicted in tRNA^Sec^ sequencing in an IGV format (**Fig. 2b**). The tRNA^Sec^ i^6^A site was validated by *in vitro* isopentenylation^17^ and analyzed by HPLC (**Fig. 2c**). Results showed a prolong retention time for the modification after isopentenylation, installed by TRIT1 enzyme, making the molecule more hydrophobic. In addition to the known i^6^A tRNA sites, we identified a novel i^6^A site at position 59 in human tRNA^Val^, adjacent to a known m^1^A site at position 58 (**Fig. 2d**). This position was detected only after IP, implying lower stoichiometry.

To test the accuracy of i^6^A-seq we selected this position for validation using several strategies. *In vitro* isopentenylation of a synthetic oligonucleotide derived from the tRNA^Val^ sequence (nts 49-69) followed by LC-MS/MS confirmed that tRNA^Val^ is a substrate of TRIT1 (**Fig. 2e**). Sequence analysis of the *in vitro* isopentenylated product showed a high mutation rate at the expected position (**Fig. 2f**) with a typical i^6^A mutation profile. Furthermore, the tRNA^Val^ site was verified by nuclease protection assay and LC-MS/MS, demonstrating presence of i^6^A is in endogenous transcripts (**Fig. S1c**). Since tRNA^Val^ is co-modified at position 58 with m^1^A, LC-MS/MS analysis revealed the enrichment of both modifications (**Fig. S1c**), which was also evident as mutations in the sequence analysis, regardless of iodine treatment or i^6^A enrichment (**Fig. 2d**). However, i^6^A enrichment was lower than that of m^1^A (molar ratios of 1.44 i^6^A/A vs. 3.04 m^1^A/A) suggesting a lower stoichiometry of i^6^A in tRNA^Val^, yet the structural inaccessibility of this position in tRNA may point to an actual higher stoichiometry. This result may explain why i^6^A was detectable only after IP. The ability of i^6^A-seq to identify new positions, even those present at low stoichiometry, underscores the suitability of this method for global mapping of i^6^A in mRNA.

Based on the sequence output of i^6^A-seq in small RNA we established a dedicated pipeline for identifying sites with high confidence. i^6^A sites were considered only when all the following criteria were met: a minimal coverage of five reads in both treated and untreated samples; a mutation rate of at least 10%; mutation rate ≥ 10 fold higher than matched untreated samples; detection of minimum two mutated reads together with a truncation pattern or at least four mutated reads without PMTs; and reproducibility, requiring that sites are detected in at least two out of three biological replicates per cell line or tissue (**Fig. S2**).

### 5. Assessing the detection of ms^2^i^6^A by i^6^A-seq

Among the 35 RNA modification tested, the only modifications that reacted with iodine under the tested conditions were i^6^A and ms^2^i^6^A (**Supplementary data file 1**). To assess whether i^6^A-seq can distinguish i^6^A from ms^2^i^6^A we compared the signals obtained from control and CDK5RAP1 knockout (KO) HAP1 cells. Since CDK5RAP1 catalyzes the conversion of i^6^A into ms^2^i^6^A, differences in i^6^A-seq signatures at known modified sites allowed us to discriminate between the two modifications.

We evaluated the frequency of mutations and PTMs at the known ms^2^i^6^A sites in human mitochondrial tRNAs^32^. For example, ms^2^i^6^A is located at position 37 in MT-TS1, the mitochondrially encoded tRNA^Ser^ 1. In iodine treated HAP1 control samples a characteristic RT signature was observed, with high rates of mutations and PMTs (**Fig. 2g (ii)**). However, unlike the i^6^A signature, high rates of mutations and truncations were evident in untreated RNA samples (**Fig. 2g (ii)**), indicating that ms^2^i^6^A can impair RTase proofreading and processivity independent of iodine labeling. In contrast, in HAP1 CDK5RAP1 KO RNA, where i^6^A is not converted to ms^2^i^6^A, the baseline levels of mutations and PMTs were low, resembling the i^6^A typical signature, and were elevated only after labeling with iodine (**Fig. 2g (iii)**). Similar results were obtained in the signature of ms^2^i^6^A in position 37 of mt-Ts1 from mouse liver RNA (**Fig. 2g (i)**). To date there are no indications for ms^2^i^6^A in nuclear encoded RNA species^33^. These results indicate that ms^2^i^6^A leaves a distinct RT signature, enabling clear discrimination between i^6^A and ms^2^i^6^A, further validating the specificity of the i^6^A-seq method.

### 6. Characterizing the human and mouse isopentenylome

Recent years have established mRNA modifications as a new frontier in gene expression regulation. Having confirmed the presence of i^6^A in mRNA by LC-MS/MS and validating the i^6^A-seq protocol, our next step was to map i^6^A at a transcriptome-wide manner. We reasoned that if i^6^A plays a biological role, at least a fraction of its positions will be reproducibly detected across different cell lines. To this end we applied i^6^A-seq to mRNA from five human cell lines: HEK293, HAP1, HeLa, HepG2 and DMS273 and identified 815, 1,657, 1,375, 1,485 and 8,233 i^6^A sites, respectively. Mapping was carried out in at least three biological replicates per cell line, considering only sites that were identified in two or more replicates. Of note, DMS273 cells exhibited a substantially greater number of sites, likely reflecting their markedly elevated TRIT1 expression levels, compared to the other tested cell lines^15^. Importantly, a large number of sites reappeared in more than one cell line (**Fig. 3a**), with over 500 sites common to all tested cell lines, alluding to a biological function. A representative i^6^A site output in IGV format reveals the enrichment, mutations and drop in coverage due to PMTs (**Fig. 3b**). Meta-site analysis of site coverage in IP samples showed that this pattern is common (**Fig. 3c**).

**Figure 3.**
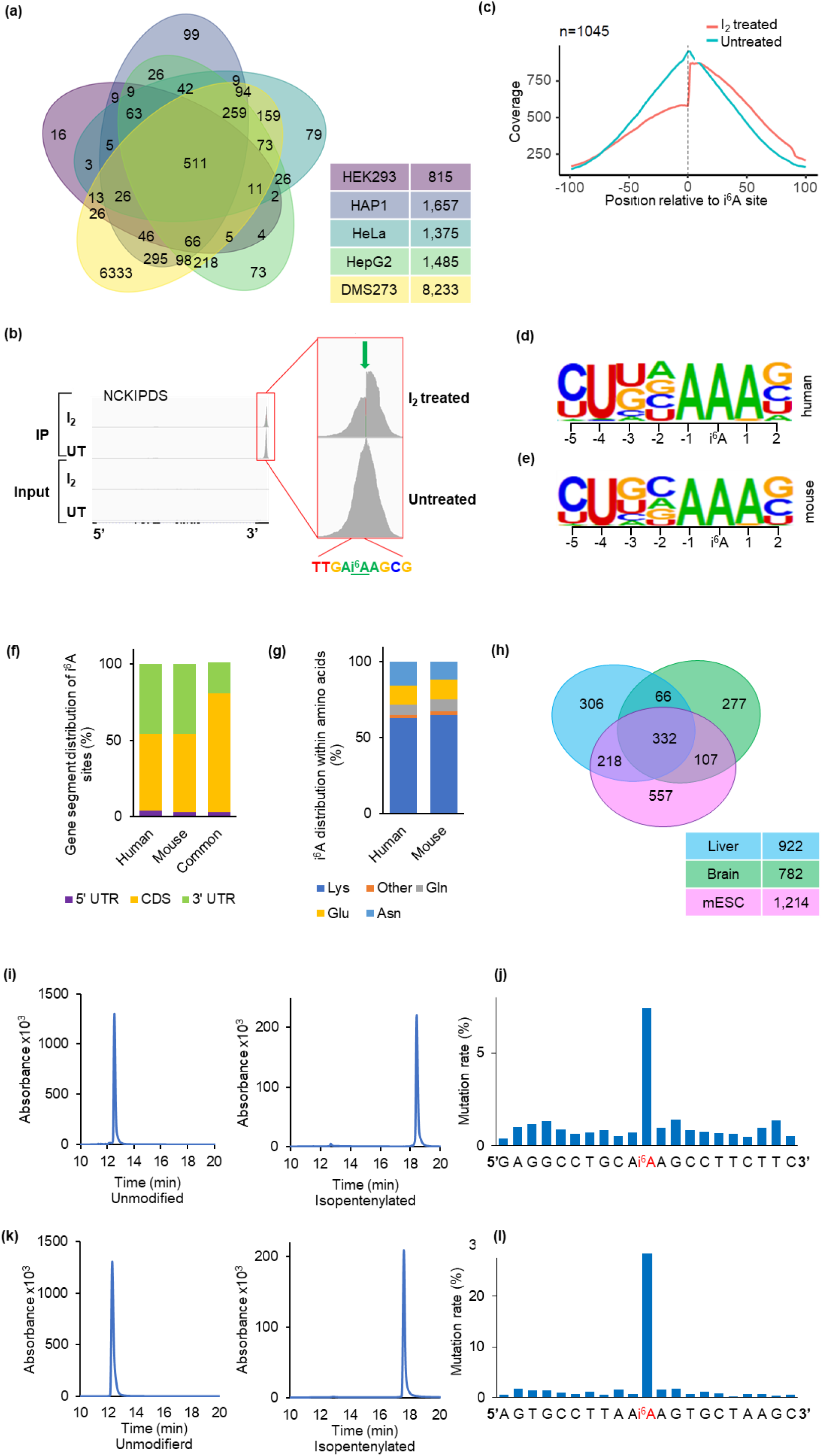
i^6^A-seq uncovers the human and mouse isopentenylome. **(a)** Venn diagram showing the number of i^6^A sites in mRNA identified in the indicated human cell lines. **(b)** IGV plots of the NCKIPSD gene, illustrating the presence of an i^6^A site. **(c)** Metagene profiles before and after iodine treatment, depicting a drop in sequence coverage just before the i^6^A position. **(d-e)** Sequence logos representing the deduced consensus of i^6^A sites in human (d) and mouse (e). **(f)** Bar plot representing distribution of i^6^A sites within gene segments in human, mouse and common sites between the two species. **(g)** Bar plot representing amino acid distribution of i^6^A sites in human and mouse. **(h)** Venn diagram showing the number of i^6^A sites in mRNA identified in the two indicated mouse tissues and mouse ESCs. **(i)** and **(k)** HPLC graphs of RNA oligonucleotides derived from representative i^6^A sites in human ESRRA (i) and mouse Pcyox1 (k), before (left) and after (right) *in vitro* isopentenylation by TRIT1. **(j)** and **(l)** Sequencing of synthetic ESRRA (j) and Pcyox1 (k) RNA oligonucleotides with i^6^A after iodine treatment showing elevated mutation rates at the modification position.

Based on identified i^6^A sites, we characterized the isopentenylome in human mRNA. An unbiased search for sequence motifs enriched around i^6^A sites revealed a consensus motif of YWNNA<u>A</u>RS (Y=C or U, W=A or U, N= any, <u>A</u>=i^6^A, R=A or G, S=G or C, *p*=1×10^−1221^, **Fig. 3d**) perfectly recapitulating the established TRIT1 consensus motif in tRNAs^34^, further supporting our conclusion that TRIT1 is the writer of i^6^A in both mRNA and tRNA. Analysis of the i^6^A distribution across transcript segments (5’UTR, CDS and 3’UTR) revealed that, on average, the vast majority of the sites were distributed nearly evenly between the CDS (50.1 %) and the 3’UTR (45.6 %), while the 5’UTR was minimally modified (4.3 %) (**Fig. 3f**). Within the CDS, i^6^A was predominantly found in lysine codons (AAA and AAG, **Fig. 3g**), which is expected from the sequence of conserved motif, suggesting a possible effect on translation.

Global mapping of i^6^A in mouse mRNA obtained from liver, brain and mouse embryonic stem cells (mESCs) identified 922, 782 and 1,214 i^6^A sites, respectively. i^6^A topology in mouse recapitulated the main features of the human isopentenylome, including sequence motif (**Fig. 3e**), site distribution along transcript segments (**Fig. 3f**), and amino acid codon preferences (**Fig. 3g**). As in human, a large group of sites was shared among all three mouse samples (**Fig. 3h**). One hundred and ninety three sites were common to human and mouse, primarily located in the CDS (78 %, **Fig. 3f**). The conservation between human and mouse underscores a biological function for i^6^A in mRNA. Selected mRNA sites in both human and mouse were validated by *in vitro* isopentenylation, followed by iodine labeling and sequencing (**Fig. 3i-l, Supplementary data file 2**).

The next essential step was to test whether TRIT1 is also an i^6^A writer in mRNA. We generated Trit1 KO in mESCs by harnessing CRISPR-Cas9 technology, targeting exons 2 and 11 to introduce a large deletion of 409 out of 467 amino acids in the Trit1 protein. Successful clones were confirmed by PCR, DNA sequencing and real-time qPCR (**Fig. 4a**). i^6^A-seq of mRNA from control and Trit1 KO mESCs revealed a drastic decrease of >93% in site numbers in Trit1 KO (**Fig. 4b)**, which was evident already in the quality control of the libraries by tape station analysis (**Fig. S3a**), attesting to Trit1 being the writer of mRNA i^6^A. Based on the expression of Thy1, which is a marker for mouse embryonic fibroblasts, not expressed in mESCs, we estimate that the 67 identified sites in Trit1 KO mESCs represent residual contamination of mRNA from irradiated feeder cells.

**Figure 4.**
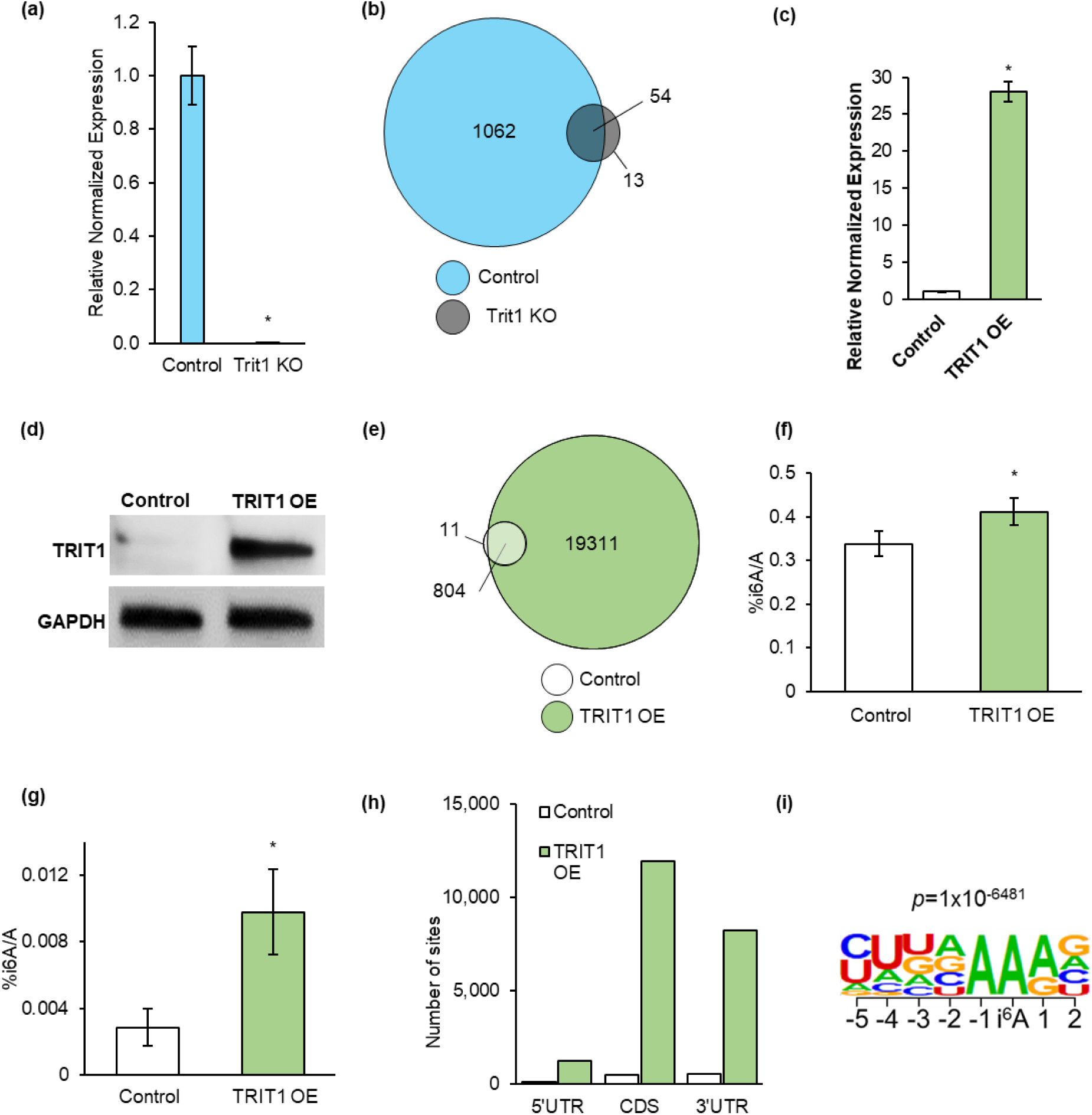
Isopentenylation is affected by TRIT1 protein levels. **(a)** Relative normalized expression of Trit1 analysis using real-time qPCR for Trit1 KO validation, *n*=3, unpaired *t*-test \**p* < 0.05. **(b)** Venn diagram showing the decreased number of i^6^A sites found in mESCs Trit1 KO compared to control. **(c)** and **(d)** Analysis of TRIT1 OE after transfection with TRIT1 plasmid in HEK293 cells, using real-time qPCR (c), *n*=3, unpaired *t*-test, \**p*<0.05 and WB (d). **(e)** Venn diagram showing the overlap of i^6^A sites found in HEK293 TRIT1 OE and control in mRNA samples. **(f-g)** LC-MS/MS quantification of i^6^A/A ratio in HEK293 control and TRIT1 OE in small RNA (f) and mRNA (g), *n*=3, unpaired *t*-test *p < 0.05. **(h)** i^6^A sites in TRIT1 OE compared to control cells, represented according to their segment location in the transcript. **(i)** Sequence logo representing the deduced consensus of i^6^A sites in TRIT1 OE cells.

To further establish the function of TRIT1 as mRNA i^6^A writer, we overexpressed (OE) it in HEK293 cells. Elevated TRIT1 levels were verified by qPCR and WB (**Fig. 4c, d**). i^6^A-seq analysis identified 20,115 sites in OE compare to 815 sites in control cells, with >98% of control sites also present in TRIT1 OE (**Fig. 4e**). These results align with an increase in i^6^A nucleoside detection in both small RNA and mRNA samples (**Fig. 4f, g**). Site distribution patterns shifted slightly, showing preference for the CDS over the 3’UTR (**Fig. 4h**). Unbiased motif search revealed a consensus sequence (**Fig. 4i**) similar to that identified in control cells (**Fig. 3d**).

Small cell lung carcinoma cell line, DMS273, express high TRIT1 levels^15^. We used a system generated by the Esteller lab^15^ that includes DMS273 cells stably expressing short hairpin RNAs to knock down (KD) TRIT1 (shTRIT1) and negative control DMS273 cells (scramble). i^6^A sites from scramble vs. TRIT1 KD mirrored the correlation of increased i^6^A site numbers with elevation of TRIT1 levels (**Fig. S3b**).

To evaluate the stoichiometry of isopentenylated transcripts, we generated a standard calibration curve by sequencing varying known ratios of iodine-treated and untreated synthetic RNA spike-in oligonucleotides, derived from the tRNA^Sec^ sequence, with several changes to allow their discrimination from any residual endogenous tRNA^Sec^ contamination in the mRNA sample. The calibration curve demonstrated a strong correlation between the fraction of affected reads and the proportion of iodine-labeled i^6^A content (**Fig. S4a**). We then performed ultra-deep sequencing of i^6^A in mRNA of mouse brain and mouse liver tissues, omitting the enrichment step and including i^6^A spike-ins in the i^6^A-seq protocol. Applying the standard curve to our data, enabled the estimation of the percentage of i^6^A modification for each site, in both human and mouse. In mouse, about 10% of the sites exhibited relatively high misincorporations and truncation rates (**Fig. S4b**). Interestingly, the same analysis, carried out on DMS273 cells identified over 600 highly isopentenylated sites, presumably a consequence of high TRIT1 levels.

Direct RNA sequencing (DRS) is a relatively new method developed by Oxford Nanopore Technologies. The method sequences intact transcripts directly and identifies changes in the electric signal that flows through the pores when the transcripts move through it. Electric output data is analyzed by typical changes of the nucleotides, producing long-read sequences^35,36^. DRS distinguishes modified nucleotides by observing disruptions in ionic current as RNA strands pass through nanopores. These disruptions create specific signal patterns that computational tools can interpret to identify RNA modifications such as m^6^A, Ψ (pseudouridine), 5-methylcytosine (m^5^C), and inosine^37,38^. We subjected mRNA of DMS273 scramble and shTRIT1 cells to DRS and analyzed it using the Dorado tool^39^. The signal output of the sequences surroundings i^6^A sites (that were identified by i^6^A-seq) showed a pronounced signature of mutations and deletions that were less prominent and even absent, in shTRIT1 compered to scramble DMS273 cells (**Fig. S5a**). Out of 210 sites examined individually, we found 172 sites (over 80%) with misincorporation or deletion patterns in scramble cells (**Fig. S5b**). Developing a robust pipeline in the future for the analysis of i^6^A, using the DRS technology, would further validate i^6^A-seq, also allowing an antibody-free approach.

Together, the human and mouse isopentenylomes not only reveal the conserved landscape of i^6^A in mammalian mRNA, but also pave the way for functional studies of this modification in regulation of gene expression.

### 7. i^6^A regulates mRNA stability in genes isopentenylated at the CDS

mRNA modifications are important regulators of gene expression, acting through diverse mechanisms including regulation of transcript stability. For instance, m^6^A promotes mRNA decay through its reader protein YTHDF2, while IGF2BPs stabilize m^6^A-modified transcripts by protecting them from degradation^4^. Other modifications such as pseudouridine (Ψ) and *N*^4^-acetylcitidine have also been implicated in mRNA stability^40,41^.

In search for biological roles of i^6^A, we selected several mRNAs that are isopentenylated at high stoichiometry, half of which are modified at the CDS, while the rest at the 3’UTR. We monitored changes in their mRNA levels upon TRIT1 OE in HEK293 cells, using real-time qPCR (**Fig. 5a**). Collectively, genes that are isopentenylated at the 3’UTR did not show significant trend in their expression after TRIT1 OE (**Fig. 5a, dark blue bars**). In contrast, genes isopentenylated at the CDS exhibited reduced mRNA levels upon TRIT1 OE, suggesting that i^6^A deposition in the CDS may promote transcript decay (**Fig. 5a, light blue bars**). The most pronounced effects were evident for SLC38A10 (coding for Solute Carrier Family 38 Member 10) and AIG1 (coding for Androgen Induced 1), exhibiting a ~75% and ~33% downregulation, respectively.

**Figure 5.**
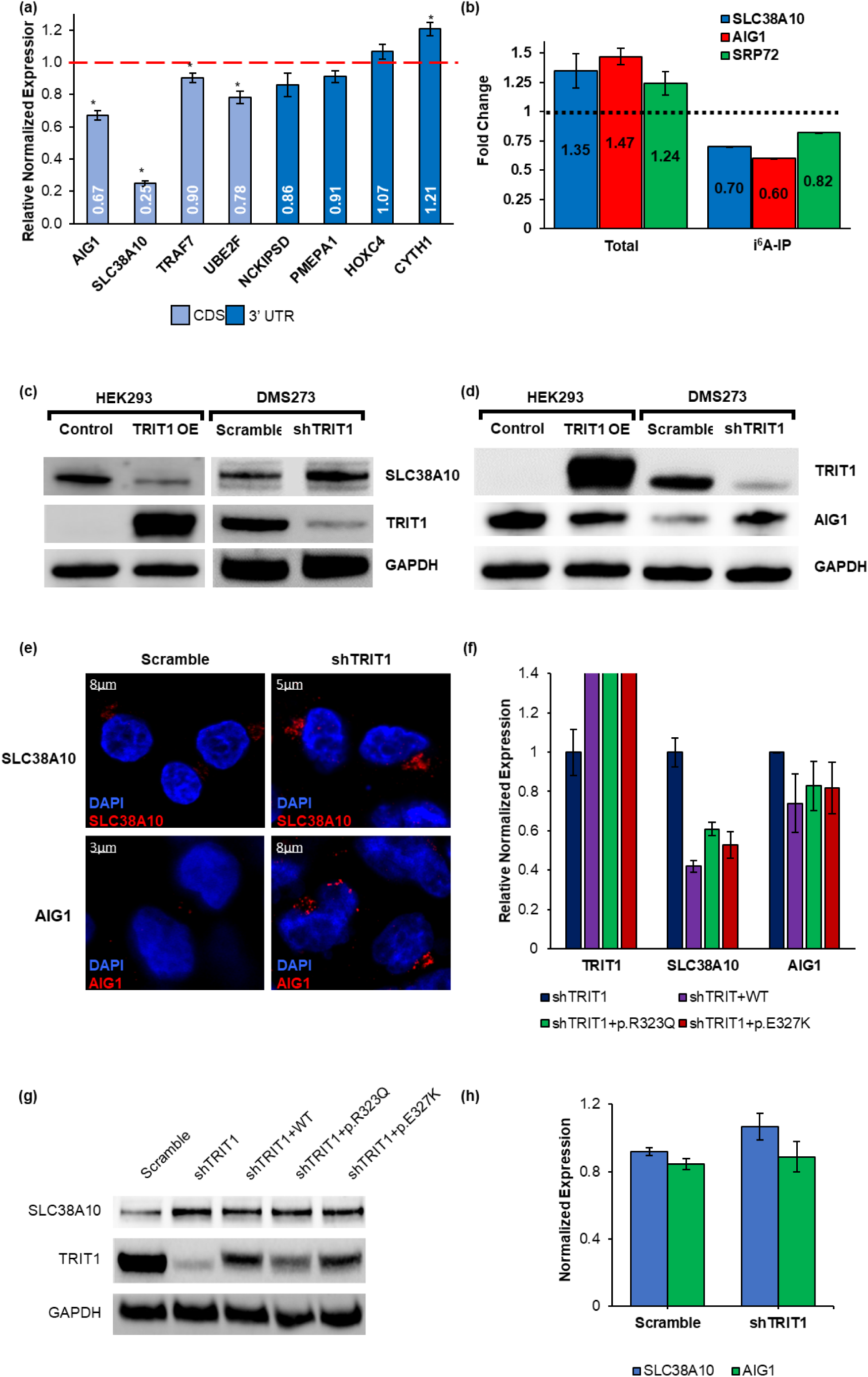
The effect of i^6^A on gene expression. **(a)** Real-time qPCR showing TRIT1 OE effect on gene expression in highly isopentenylated genes in HEK293 cells, *n*=3, unpaired *t*-test, \**p*<0.05. **(b)** Real-time qPCR of the indicated genes in shTRIT1 cells compared to their control, before and after i^6^A-IP, *n*=3. **(c-d)** WB of the indicated cell lines, after TRIT1 OE/KD, displaying an inverse change in SLC38A10 (c) and AIG1 (d) in response to shifts in TRIT1 protein expression. **(e)** Immunofluorescence staining images of SLC38A10 and AIG1 in scramble and shTRIT1 DMS273 cells, revealing their increased expression under TRIT1 KD. **(f)** Analysis of TRIT1, SLC38A10 and AIG1 after TRIT1 WT or mutant rescue in shTRIT1 cells by real-time qPCR, *n*=3, unpaired *t*-test, \**p*<0.05. **(g)** WB of the indicated cell lines, representing similar trend for TRIT1 and SLC38A10 as in (f). **(h)** Real-time qPCR of SLC38A10 and AIG1 mRNA levels, in DMS273 scramble versus shTRIT1, monitored 6 hours after Actinomycin-D addition, compared to 0 hours, *n*=3.

Conversely, reduced TRIT1 levels in DMS273 shTRIT1 cells resulted in a marked increase in the mRNA levels of both SLC38A10 and AIG1 genes (**Fig. 5b, Total**). To directly assess changes in the isopentenylation fraction of these transcripts, we performed i^6^A-IP on mRNA from DMS273 scramble and shTRIT1 cells and evaluated the levels of the i^6^A-modified fractions of SLC38A10 and AIG1 by real-time qPCR. As expected, reduced TRIT1 levels significantly decreased the fraction of i^6^A-modified SLC38A10 and AIG1 mRNAs (by 22% and 29%, respectively, **Fig. 5b, i^6^A-IP**), implying that the elevation in total mRNA levels of these genes is attributed to reduced i^6^A isopentenylation. We also assessed the changes in SRP72 that is highly isopentenylated in DMS273 but not in HEK293. SRP72 showed the same trend as SLC38A10 and AIG1, but to a lesser extent (increasing by 22% in the total samples and decreasing by 12% in the IP, **Fig. 5b**). These results suggest that while the total levels of these transcripts increase under TRIT1 KD, the fractions of the isopentenylated transcripts decrease, drawing a link between isopentenylation and transcript decay.

The effects of i^6^A on the expression of SLC38A10 and AIG1 were also assessed at the protein level, revealing an inverse correlation between TRIT1 expression and the abundance of these proteins (**Fig. 5c, d**). Immunofluorescence staining of SLC38A10 and AIG1 in the DMS273 cell model further corroborated our findings (**Fig. 5e**) and providing additional support for a biological role of i^6^A in regulating gene expression.

To determine whether this regulation depends on TRIT1 enzymatic activity, we generated two mutated TRIT1 plasmids, based on previously characterized human TRIT1 mutations^27,28,42^, and assessed their impact on the expression of SLC38A10 and AIG1. The enzymatic activity of the two TRIT1 mutants was evaluated by *in vitro* isopentenylation of a tRNA^Sec^-derived oligonucleotide. As expected, both mutants exhibited only partial isopentenylation activity, compared to wild type (WT) TRIT1, reflecting their impaired function (**Fig. S6a-e**). All TRIT1 constructs increased TRIT1 expression, at both the RNA and at the protein levels (**Fig. 5f, g**). OE of WT TRIT1 in DMS273 shTRIT1 cells resulted in a robust decrease in SLC38A10 (**Fig. 5f, g**) and AIG1 (**Fig. 5fa**) expression levels, whereas OE of TRIT1 mutants produced a milder reduction in RNA expression (**Fig. 5f, g**). Still, the expression levels of SLC38A10 and AIG1 were higher when TRIT1 mutants were OE, compared to their levels after WT TRIT1 OE. Together, these results support a role for i^6^A in regulation of transcripts isopentenylated at the CDS, with TRIT1 enzymatic activity being critical for this effect.

To validate the effect of i^6^A on mRNA stability of SLC38A10 and AIG1, we monitored their half-life values by real-time qPCR following transcription inhibition with actinomycin-D in the DMS273 system (**Fig. 5h**). In DMS273 shTRIT1 cells, where i^6^A levels are lower, the half-life of SLC38A10 mRNA is higher (T_1/2_ = 8.6 and 10.4 hours in control (scramble) and shTRIT1 cells, respectively) (**Fig. 5h**). A similar effect was observed for AIG1 (T_1/2_ = 4.5 and 6.2 hours in control (scramble) and shTRIT1 DMS273 cells, respectively) (**Fig. 5h**). These findings demonstrate that i^6^A deposition promotes the degradation of SLC38A10 and AIG1 mRNA in the cell.

### 8. DIS3L2 is a novel reader protein of i^6^A

mRNA modifications exert their regulatory roles by recruiting specific reader proteins. To identify novel readers that are involved in i^6^A-dependent mRNA degradation, we harnessed the RNA affinity chromatography (RAC) approach. Specifically, two pairs of RNA oligonucleotide baits were designed based on the SLC38A10 and AIG1 sequences surrounding their i^6^A positions. Each pair comprising one isopentenylated (i^6^A-modified) and one unmodified (control) oligonucleotides. Baits were incubated with HEK293 cell lysates and analyzed by LC-MS/MS. Proteomic analysis revealed that the mRNA exoribonuclease DIS3L2 selectively bound the isopentenylated SLC38A10 bait but not its unmodified counterpart (**Fig. 6a**), identifying DIS3L2 as the first i^6^A reader protein. DIS3L2 is an RNA-binding protein with 3′-to-5′ exoribonuclease activity that mediates degradation of several RNA species and is known to target transcripts that are translated on ER-bound ribosomes^43,44^. Both SLC38A10 and AIG1 proteins are localized at the Golgi/ER organelles, thus translated on ER-bound ribosomes^45–47^.

**Figure 6.**
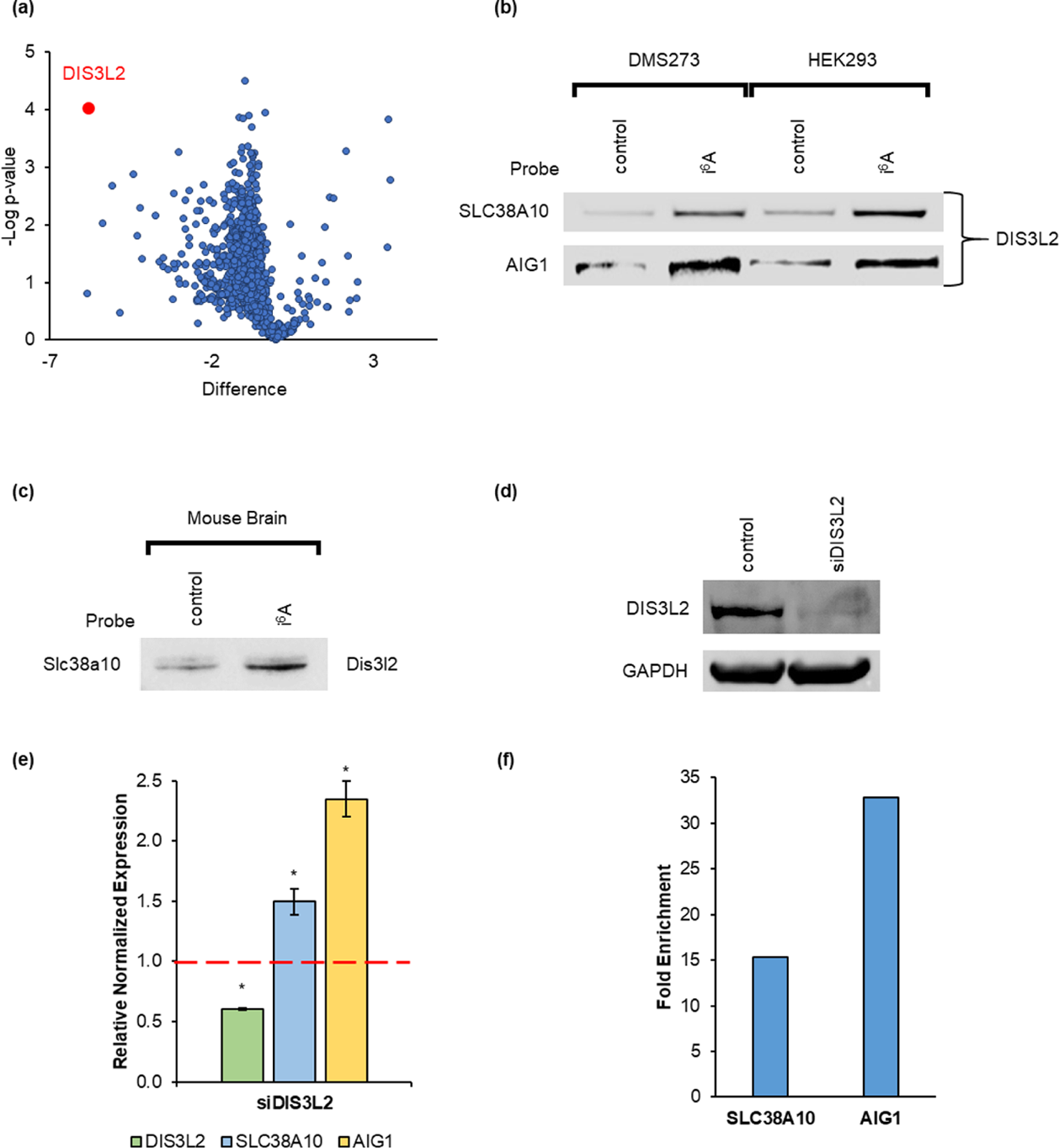
RNA affinity chromatography identifies DIS3L2 as a novel reader protein of i^6^A. **(a)** Volcano plot of proteins bound to isopentenylated versus control RNA baits with a fragment sequence of SLC38A10. Assigned in red is DIS3L2, an enriched protein with the largest difference between baits and most significant *p*-value. **(b-c)** WB validation of RAC approach showing DIS3L2 as a significant i^6^A binding protein in the indicated cell lines and RNA targets (b) and in mouse brain tissue for SLC38A10 (c). **(d)** WB validation of DIS3L2 KD after 4 days in DMS273 cells. **(e)** Validation of DIS3L2 KD in DMS273 cells by real-time qPCR, revalidating its effect on SLC38A10 and AIG1, *n*=3, unpaired *t*-test, \**p*<0.05. **(f)** Real-time qPCR analysis of RIP assay for SLC38A10 and AIG1 interaction with DIS3L2 in DMS273 cells.

The i^6^A-dependent binding of DIS3L2 to isopentenylated sequences derived from SLC38A10 and AIG1 was further validated by Western blot analysis in HEK293 and DM273 scramble cell lines (**Fig. 6b**). Slc38a10 is also orthologically isopentenylated in mouse. Thus, we performed the RAC assay using mouse brain lysate, revealing that Dis3l2 is an i^6^A reader in mouse as well (**Fig. 6c**). These results provide further reinforcement for the functionality of i^6^A as a regulator of gene expression. To confirm the functional relevance of DIS3L2 as an i^6^A reader that exerts it regulatory role by enhancing mRNA degradation, we performed siRNA-mediated KD of DIS3L2 in DMS273 scramble cells (**Fig. 6d, e**). DIS3L2 KD resulted in increased expression of both SLC38A10 and AIG1 (**Fig. 6e**), indicating that it is responsible for their degradation. Further validation of DIS3L2 binding to isopentenylated transcripts was carried out by ribonucleoprotein immunoprecipitation (RIP) assay. RNA isolated from scramble DMS273 cells was incubated with Flag-tag DIS3L2. Quantification of the RNAs was carried out using real-time qPCR. A significant fold enrichment increase for both SLC38A10 and AIG1, in comparison to Flag-tagged control lysates was observed (**Fig. 6f**).

Together, our findings establish i^6^A as a new epitranscriptomic mark in mRNA, with a biological function, and identified DIS3L2 as its reader. Our findings provide mechanistic insight into i^6^A-mediated regulation of gene expression, paving the way for the discovery of additional roles for this modification.

## Discussion

In the last decade, a new layer of control was rapidly exposed, centered on the ability of mRNA modifications to recruit cellular machineries through binding of reader proteins^2,4,48^. Most of the information emerged from m^6^A studies^2^, but with time, data regarding other, less abundant mRNA modifications, was uncovered as well, extending the regulatory layer of mRNA modifications^49^. Our work reveals i^6^A as a novel epitranscriptomic mark in mammalian mRNA. By development and validation of i^6^A-seq, we exposed the first transcriptome-wide maps of i^6^A in human and mouse, and established TRIT1 as its mRNA writer, beyond tRNA^6^. Although the abundance and stoichiometry of i^6^A in mRNA appears to be low, a subset of sites was conserved among different human cell lines and even between mouse and human, alluding to a biological function. Indeed, using RAC, we discovered the first reader protein of i^6^A, DIS3L2, an RNA exoribonuclease^43^ that mediates i^6^A-dependent mRNA decay.

i^6^A was so far known as a tRNA modification, localized to the A37 position adjacent to the anticodon^8^. Earlier studies for the role of i^6^A revealed that it stabilizes the interaction of the codon with the anticodon, enhancing accurate decoding and preventing frameshifting^6,7^. This prevents errors in decoding of similar codons and affects translation efficiency^9^.

Mutations in DIS3L2 are associated with human diseases, such as the Perlman syndrome, a rare overgrowth condition that is characterized by fetal macrosomia and renal abnormalities and predisposition to Wilms’ tumor^44,50^. Although its engagement in regulation of cell cycle progression, cell proliferation and neural crest development in zebrafish^51–53^, was demonstrated, its associated regulatory networks are still unclear. Interestingly, TRIT1 mutations that reduce its isopentenylation activity affect neurodevelopment, resulting in phenotypes such as intellectual disability, developmental delay and microcephaly^54^. Yet, the cellular machineries that are impaired in patients carrying TRIT1 mutations are still uncertain. TRIT1 is also known to play a role in cancer progression, specifically lung, as its amplification is correlated with drug resistance^15^. Data-mining of TRIT1 RNA expression levels in various type of tumors reveals significantly higher TRIT1 mRNA levels, compared to normal tissues^55,56^ (**Fig. S7**). This implies a potential, yet uncharacterized, role for TRIT1 in oncogenesis. Notably, indications of DIS3L2 involvement in tumor growth in various tissues, including lung, start to accumulate^52,57^. Thus, bridging the knowledge gaps may shed light on the TRIT1-mRNA i^6^A-DIS3L2 axis and its engagement in regulation of gene expression.

The biological importance of low-abundance mRNA modifications, such as m^1^A, provoked a controversy over its transcriptome-wide mapping, which stirred the epitranscriptomic field in 2016^24^. Some scientists suggested that the sites of m^1^A reflect a “leakage” of its tRNA writer, and that the sites have no biological importance^58,59^. This notion proved wrong over the years with a growing number of mRNA m^1^A sites with a regulatory function, uncovering the roles of this mRNA modification in neuronal gene expression and oxygen glucose deprivation/reoxygenation induction^60^, as well as in regulation of glycolysis in cancer cells, through modulating ATP5D^21^.

Still, the low abundance of i^6^A in mRNA raises the doubt on the significance of this regulatory mechanism. At this time, we focused on two gene transcripts subjected to i^6^A-dependent mRNA degradation by DIS3L2, both isopentenylated at the CDS and translated on the ER. As DIS3L2 is known to regulate the degradation of transcripts translated on ER-bound ribosomes^44^, other genes known to be similarly translated, yet isopentenylated at a lower stoichiometry, should be studied. Nevertheless, regulation of gene expression through i^6^A-dependent degradation of mRNA may be indispensable under certain stress conditions or during specific developmental stages. More experiments are needed to find the developmental time window or the stress conditions at which mRNA isopentenylation is essential.

mRNA modifications form an intricate and highly sensitive network that can regulate every step in mRNA metabolism. As mentioned above most of the data was obtained from m^6^A research. However, this network expanded continuously over the last decade, with the identification of less abundant mRNA modifications and their associated proteins (writers, erasers and readers)^48,49,61^. In addition, evidence into crosstalk between mRNA modification, for example m^6^A and m^1^A, starts to accumulate further illustrating the sensitivity of mRNA modification-dependent regulation of gene expression^62^. The question that needs to be asked is what is the advantage of having multiple modifications, readers and checkpoints for gene expression regulation? The answer lays in the nature of RNA. Unlike DNA, each gene transcript is produced in multiple copies, which are exported from the nucleus to the cytoplasm for translation and eventually degradation. The use of various modifications at different sites and in varying stoichiometry provides not only a combinatorial tool for extra-fine tuning, using different regulation mechanisms but also produce a slip-proof network that can act at different checkpoints to guarantee the needed control tailored for each mRNA.

Our research pioneers in the study of i^6^A in mRNA. Adding i^6^A to the growing assembly of the epitranscriptome, identifying its role in gene expression and uncovering the first i^6^A reader protein, can highlight new targets for therapy, paving the way for new drug development.

## Acknowledgments

The authors thank the Kahn Family Foundation; the Flight Attendant Medical Research Institute (FAMRI); the Varda and Boaz Dotan Research Center in Hemato-Oncology, Tel Aviv University; the Ernest and Bonnie Beutler Research Program; and the Israel Innovation Authority, for their continuous support of our research.

