## Supplementary Material for "i^6^A-seq maps *N*^6^-isopentenyladenosine and uncovers its role as a regulator of mRNA stability through recruitment of DIS3L2"

Supplementary Figures 1-7

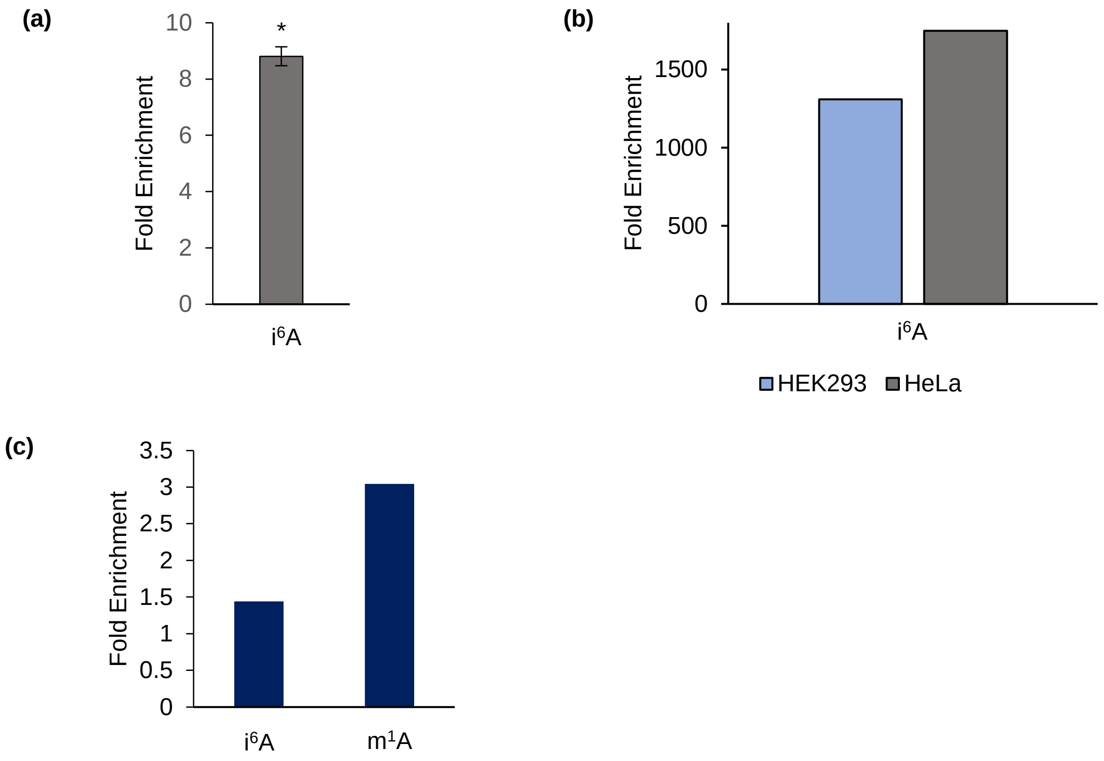


**Figure S1. Quantitative analyses of $i^6A$/A ratio by LC-MS/MS. (a-b)** $i^6A$/A ratio following enrichment with anti-$i^6A$ antibody in yeast tRNA, mean values ± s.e.m. are shown, *n*=3 (a), and in human HEK293 and HeLa mRNA samples (b). **(c)** Validation of increased $i^6A$/A ratio in $tRNA^{Val}$ by nuclease protection assay. $m^1A$/A ratio is increased as well, reflecting its proximity to the $i^6A$ site.

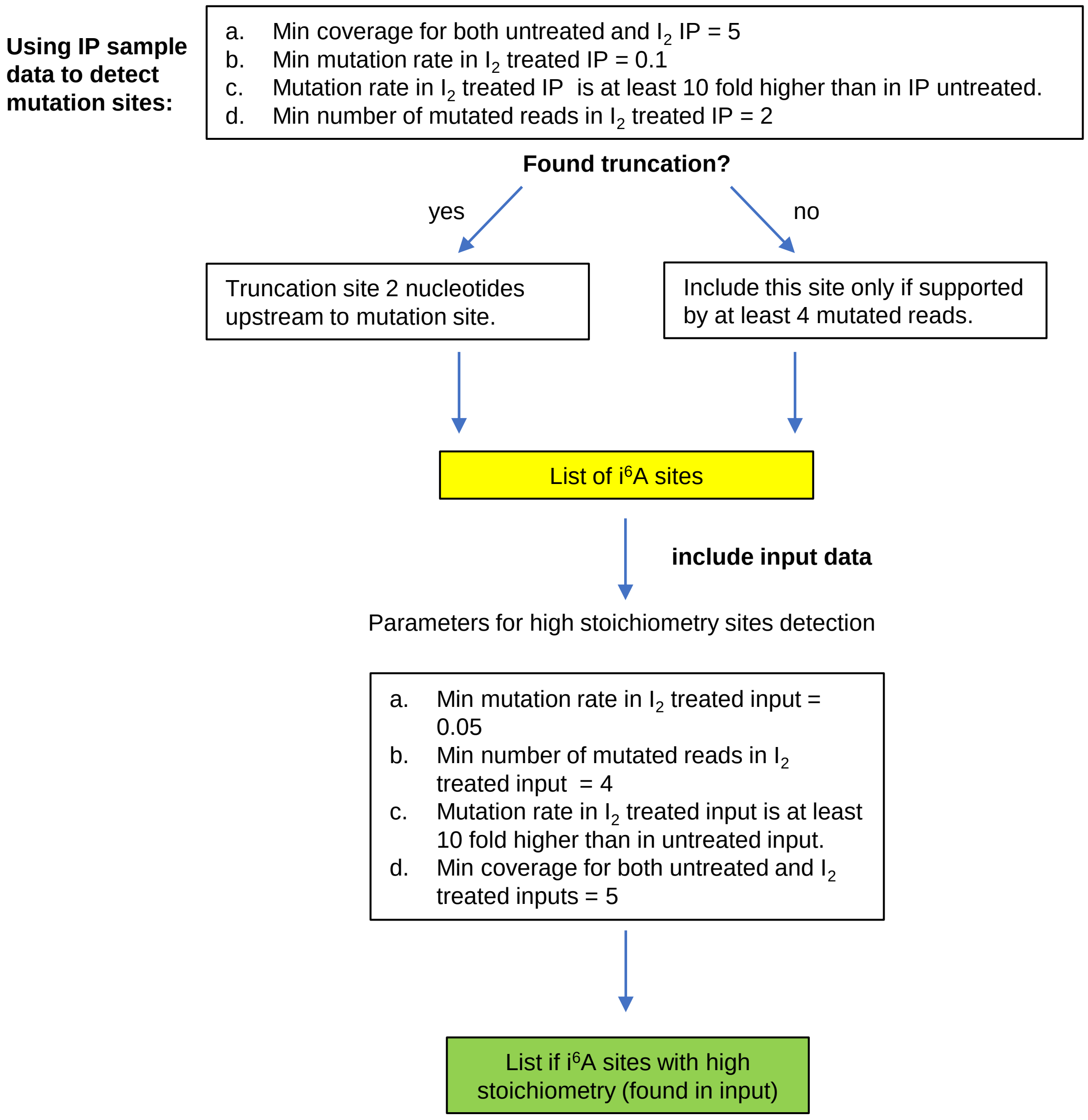


**Figure S2. Parameters for $i^6A$ site detection pipeline.**

**(a)**

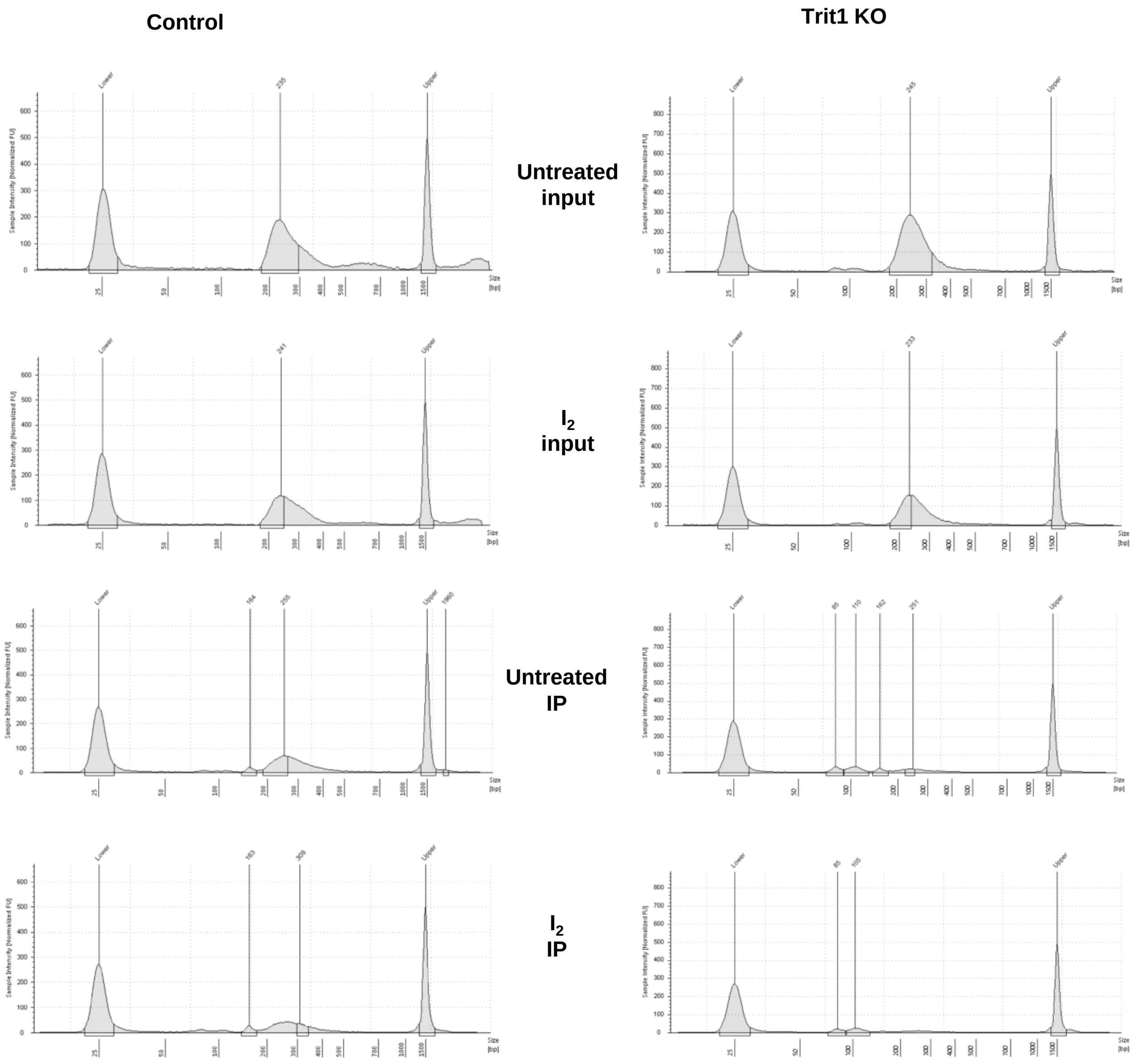


**(b)**

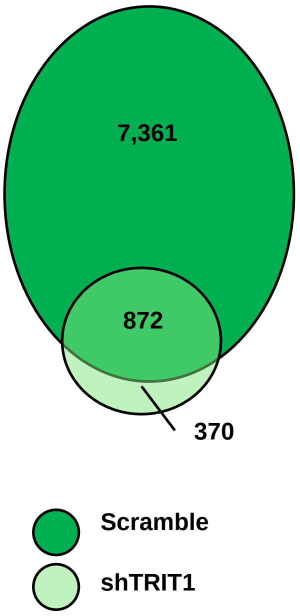


**Figure S3. Reduced TRIT1 levels further validates i$^6$A sites prevalence (a)** Example of TapeStation panels for i$^6$A-seq libraries obtained from control and Trit1 KO samples, depicting the lack of i$^6$A in IP samples when Trit1 is knocked out in mESCs. **(b)** Venn diagram showing the reduced number of i$^6$A sites in mRNA identified in DMS273 shTRIT1 compared to control (scramble) cells.

(a)

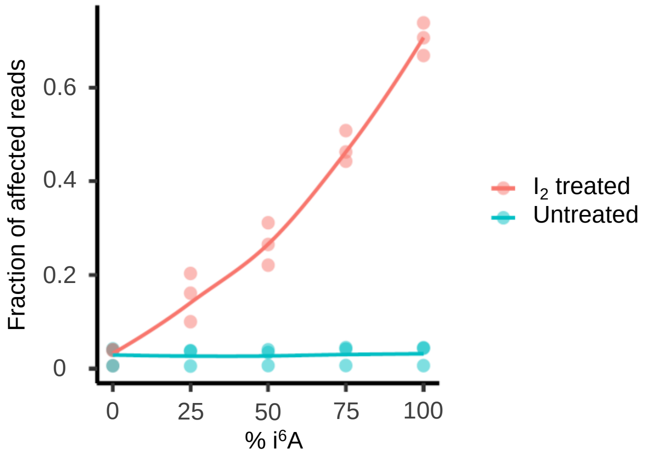


(b)

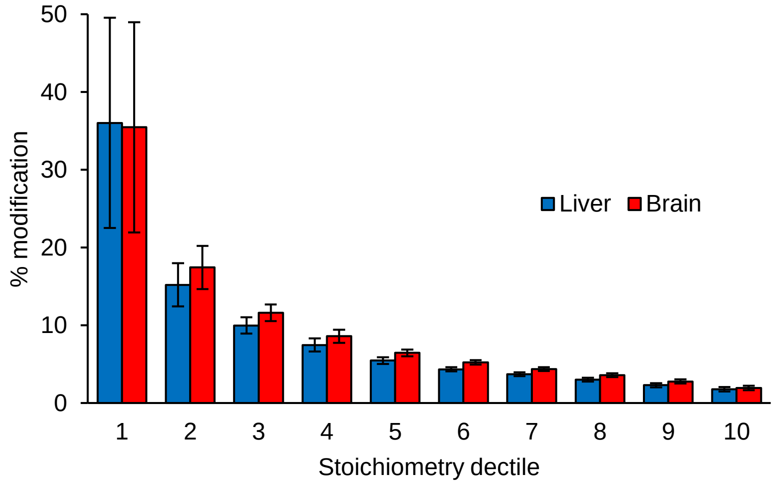


**Figure S4. i$^6$A stoichiometry. (a)** Calibration curve showing the fraction of affected reads at different percentages of i$^6$A. **(b)** Stoichiometry distribution of i$^6$A sites identified in ultra-deep sequencing of unenriched mRNA samples from mouse liver and brain.

(a)

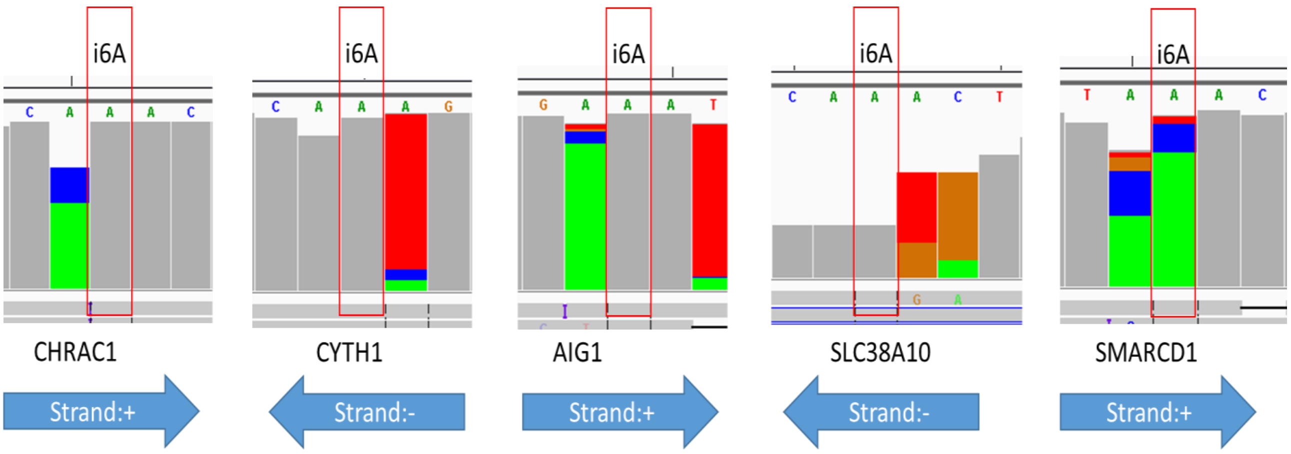


(b)

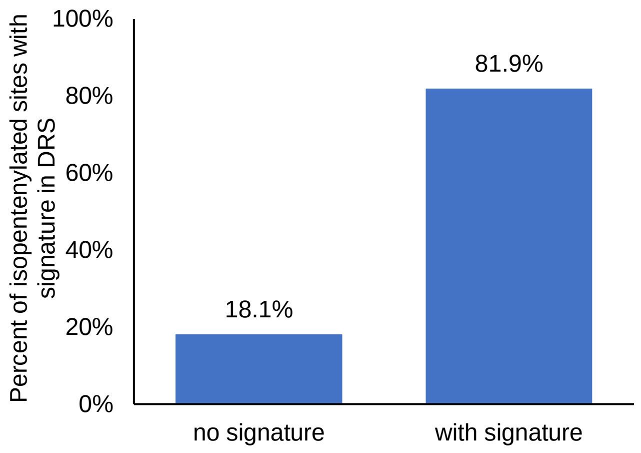


**Figure S5. Analysis of i⁶A mRNA sites using the DRS technology exposing a unique signature. (a)** IGV plots of selected i⁶A sites, illustrating a signature. **(b)** A bar plot representing the percentage of i⁶A sites exhibiting a DRS signature, out of 210 i⁶A sites that were identified by i⁶A-seq.

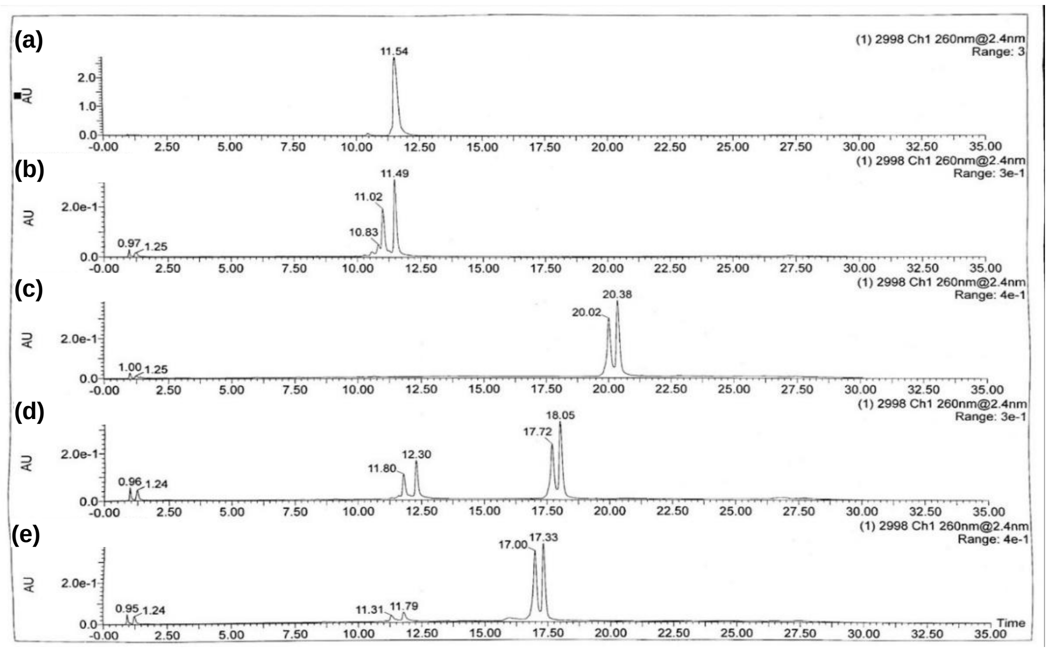


**Figure S6. Validation of TRIT1 WT and mutant protein activity by *in vitro* isopentenylation.** HEK293 cells were transfected with WT or mutant TRIT1-Flag and incubated with tRNA$^{Sec}$-derived oligonucleotide. HPLC retention times of the oligonucleotides are shown. **(a)** untreated; **(b)** untransfected; **(c)** transfected with WT TRIT1-Flag; **(d-e)** transfected with mutant TRIT1-Flag.

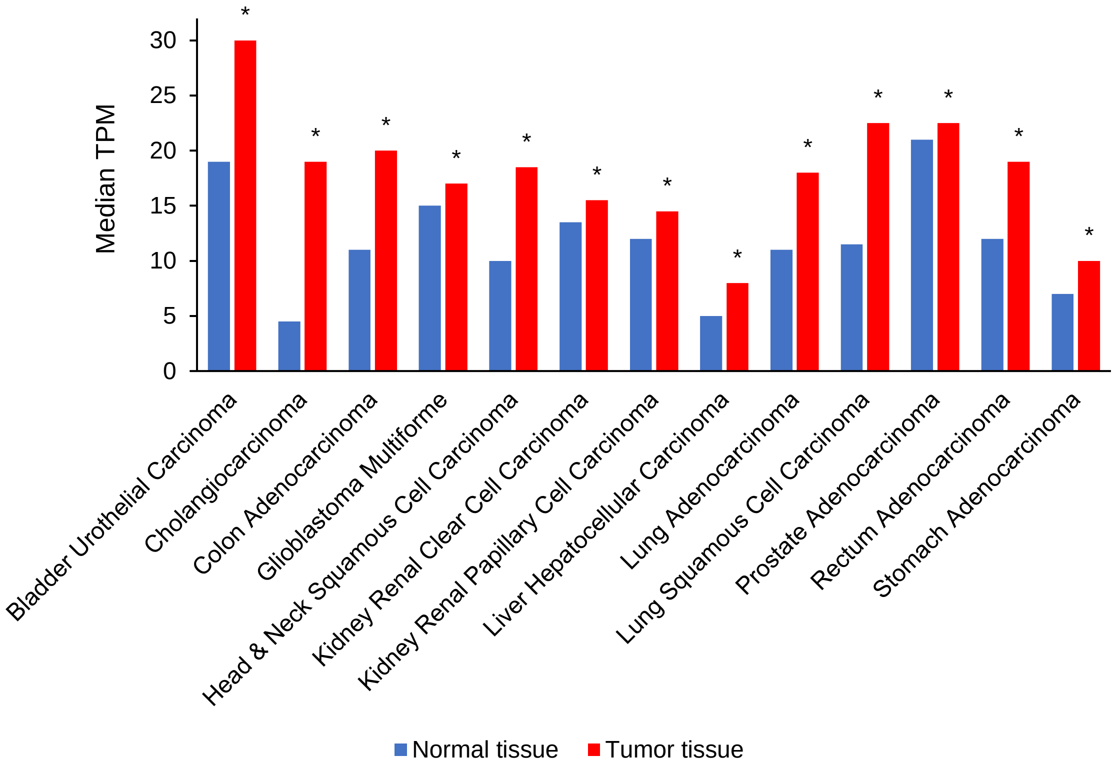


**Figure S7. TRIT1 mRNA expression is upregulated in various malignancies.** RNA expression of TRIT1 in 13 analyzed types of cancers, exhibiting significantly higher levels in tumors, *$p$<0.05.

Supplementary Data 1

### 5-Carbamoyl-methyluridine (ncm$^5$U)

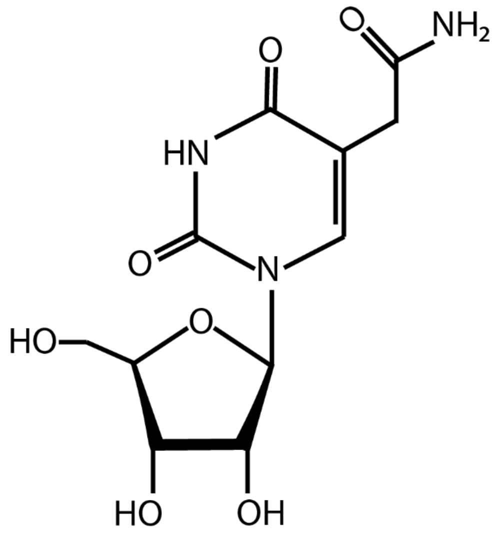


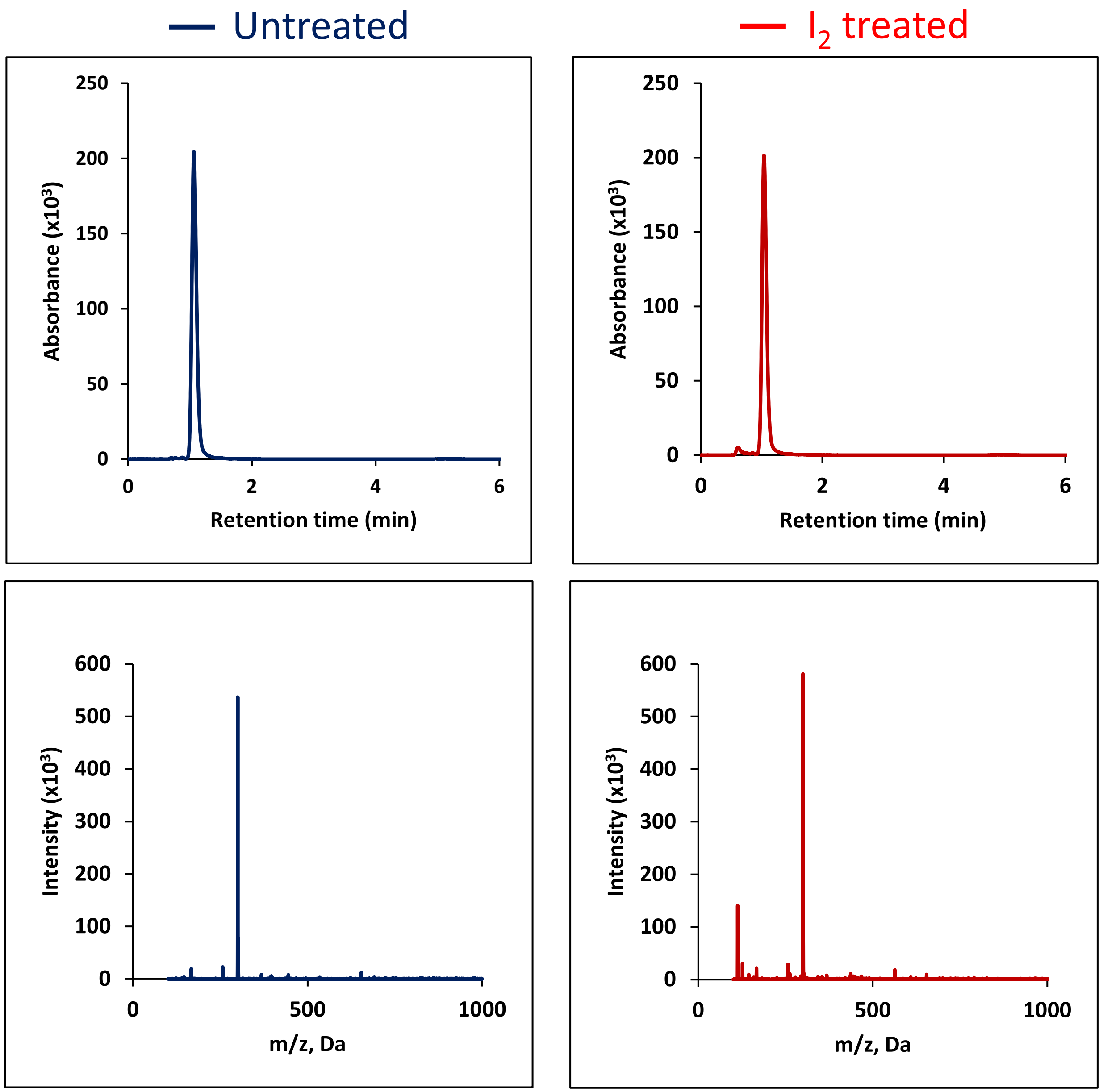

### 5-Methoxycarbonyl-methyluridine (mcm$^5$U)

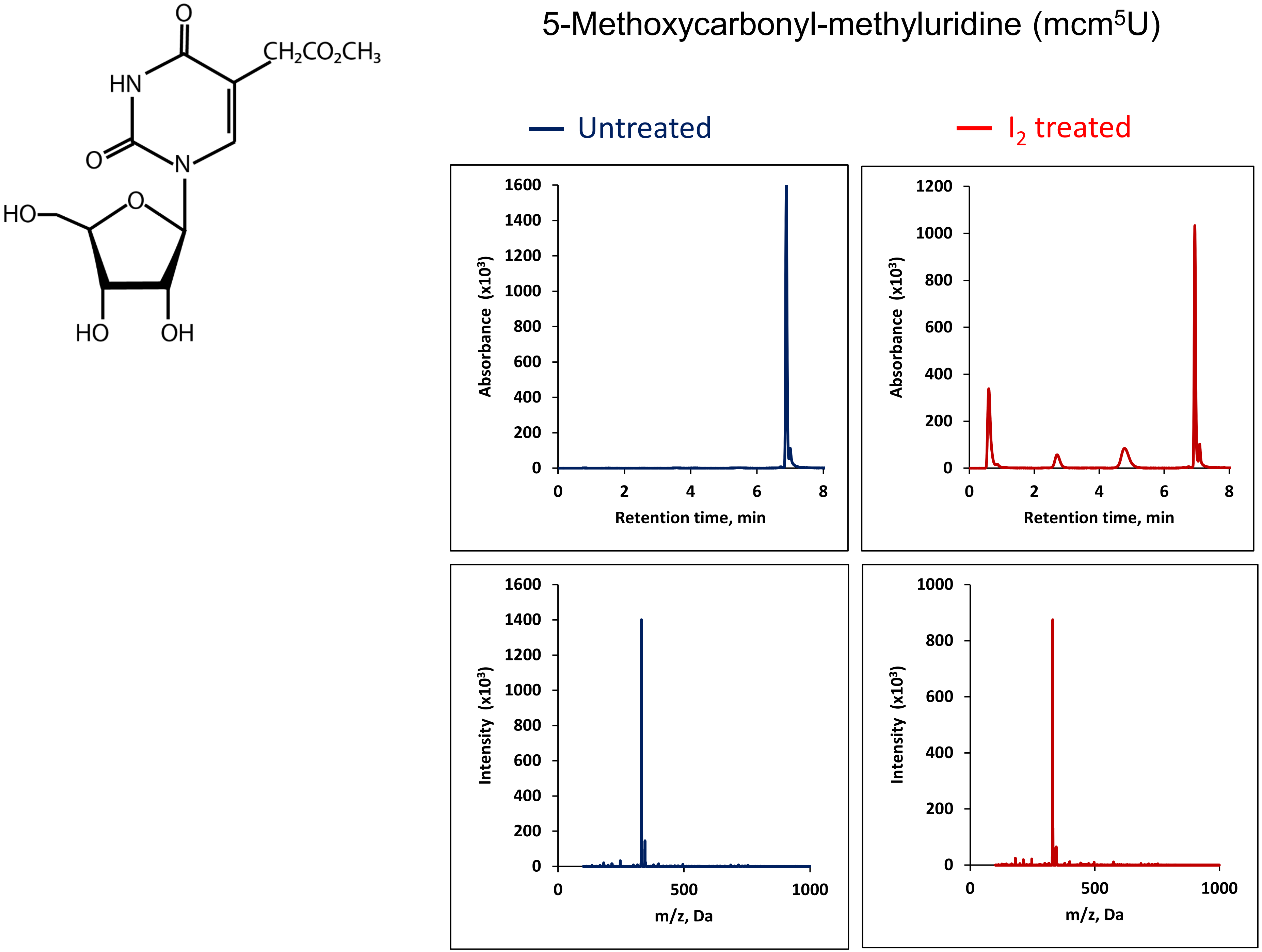

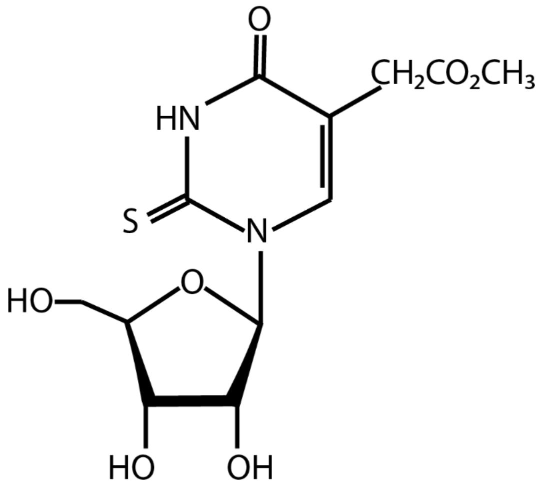


### 5-Methoxycarbonylmethyl-2-thiouridine ($mcm^5s_2U$)

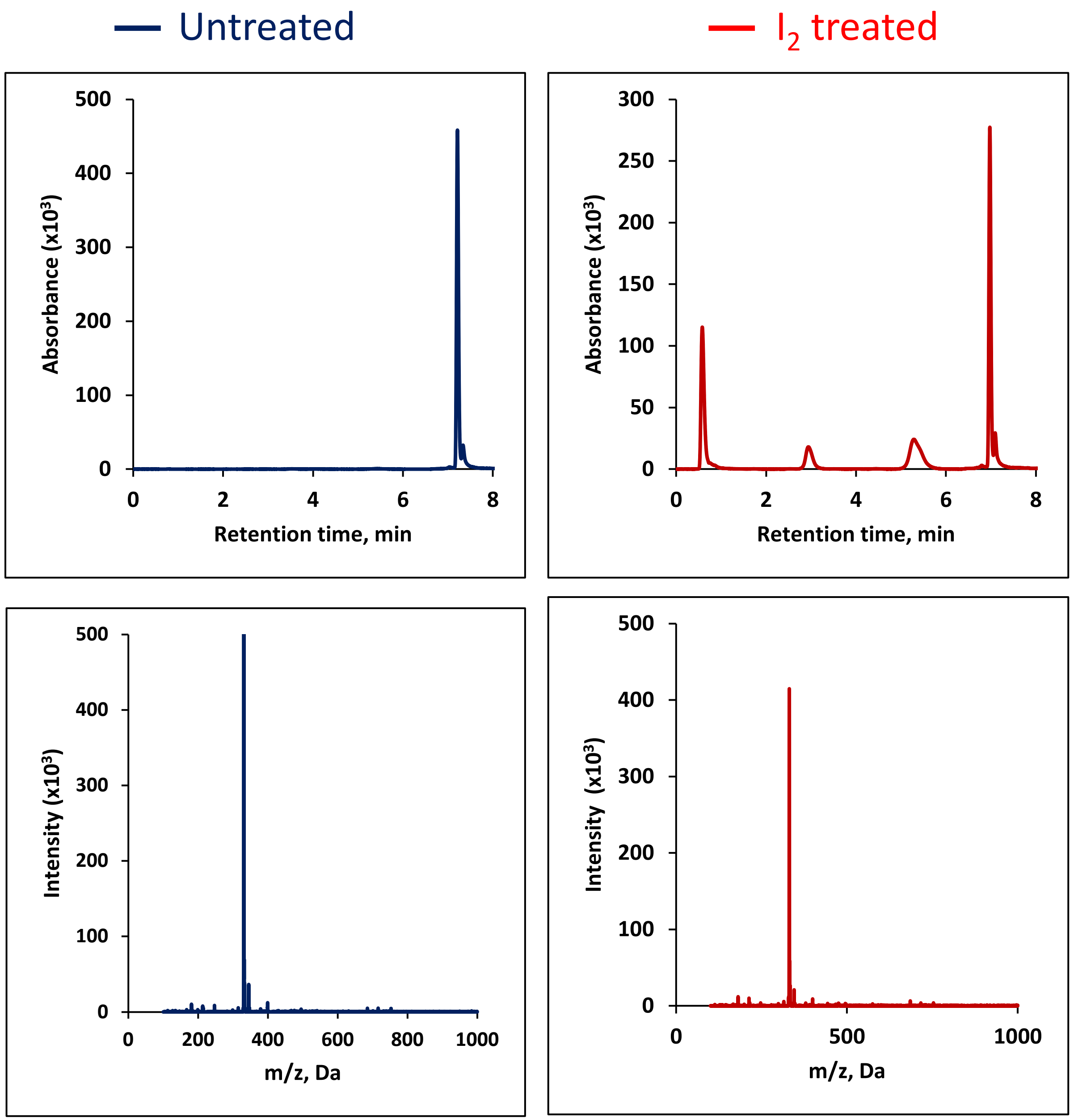

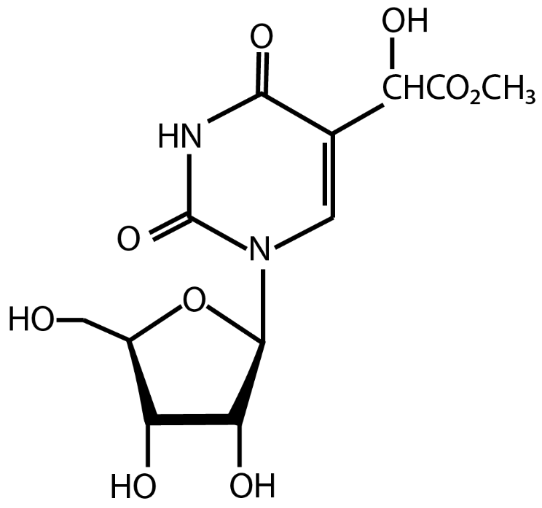


### 5-(carboxyhydroxymethyl)uridine methyl ester (mchm$^5$U)

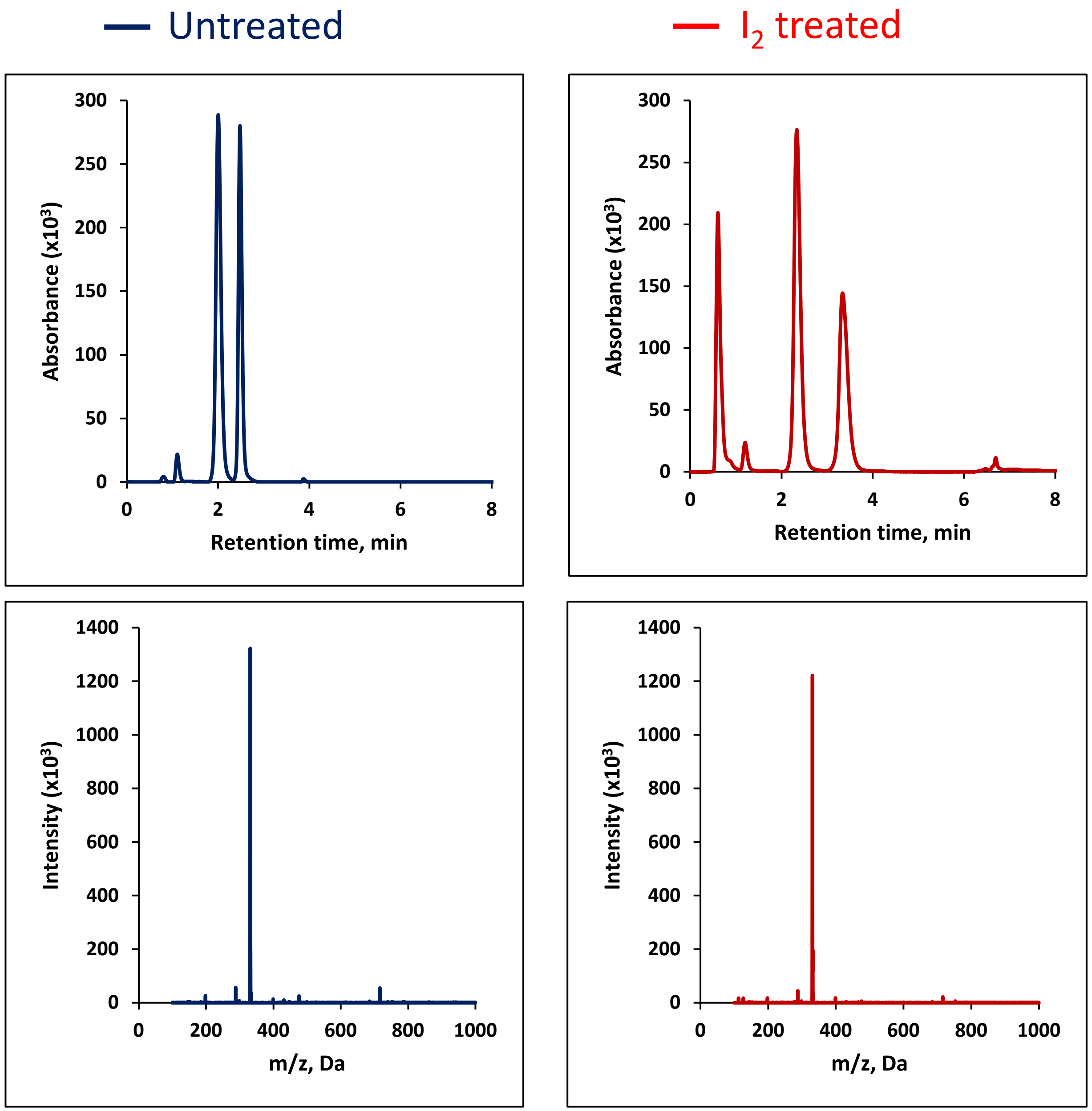

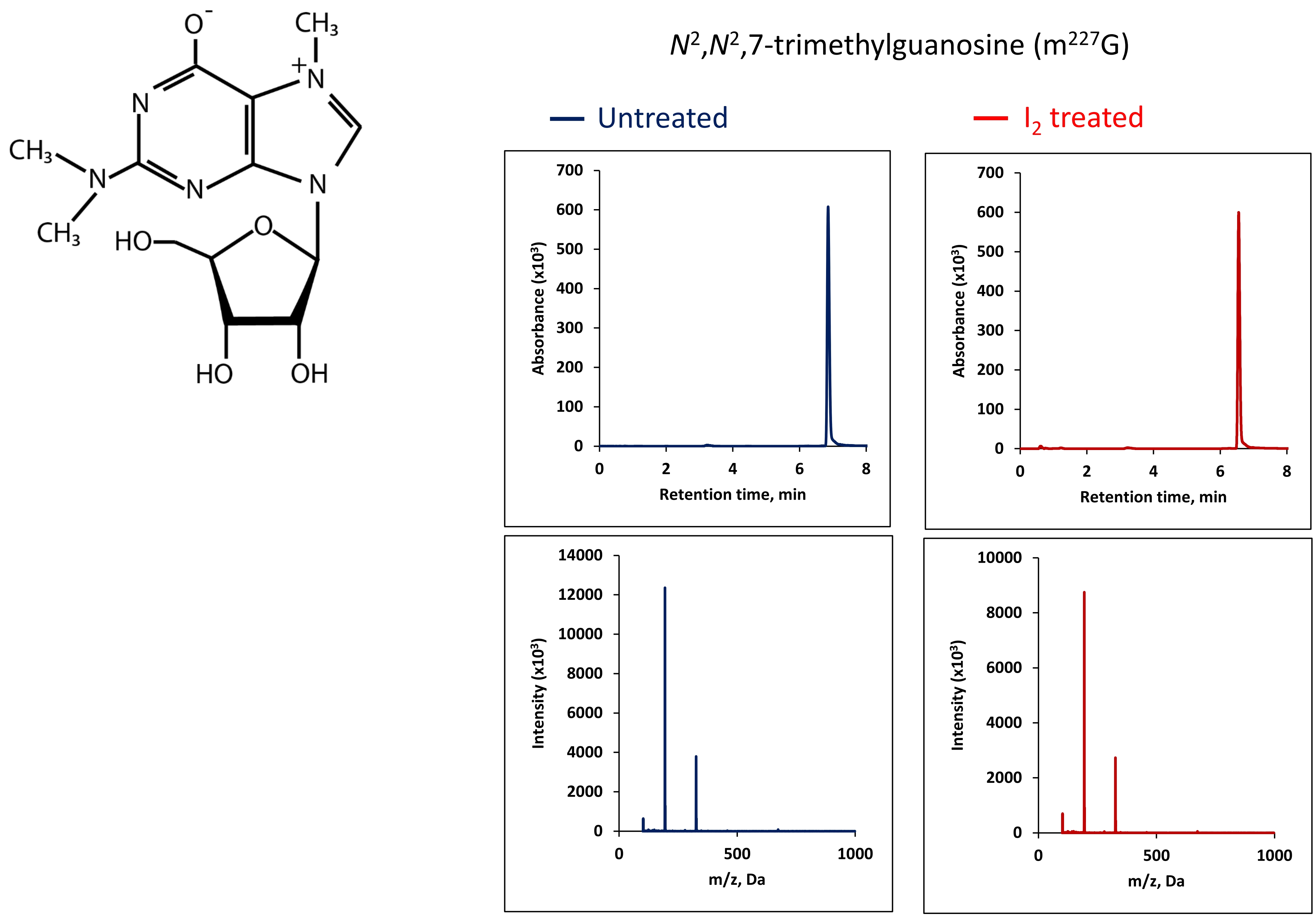

N2,N2,7-trimethylguanosine (m227G)
Untreated
I2 treated
Absorbance (x10³)
Retention time, min
Intensity (x10³)
m/z, Da
O-
CH3
N
CH3
HO
OH

### $N^2$,7-dimethylguanosine ($m^{27}G$)

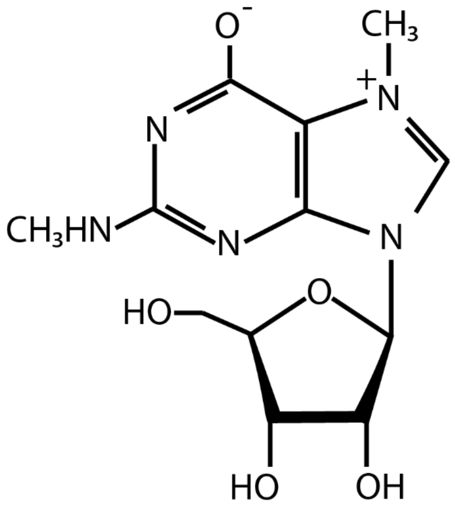


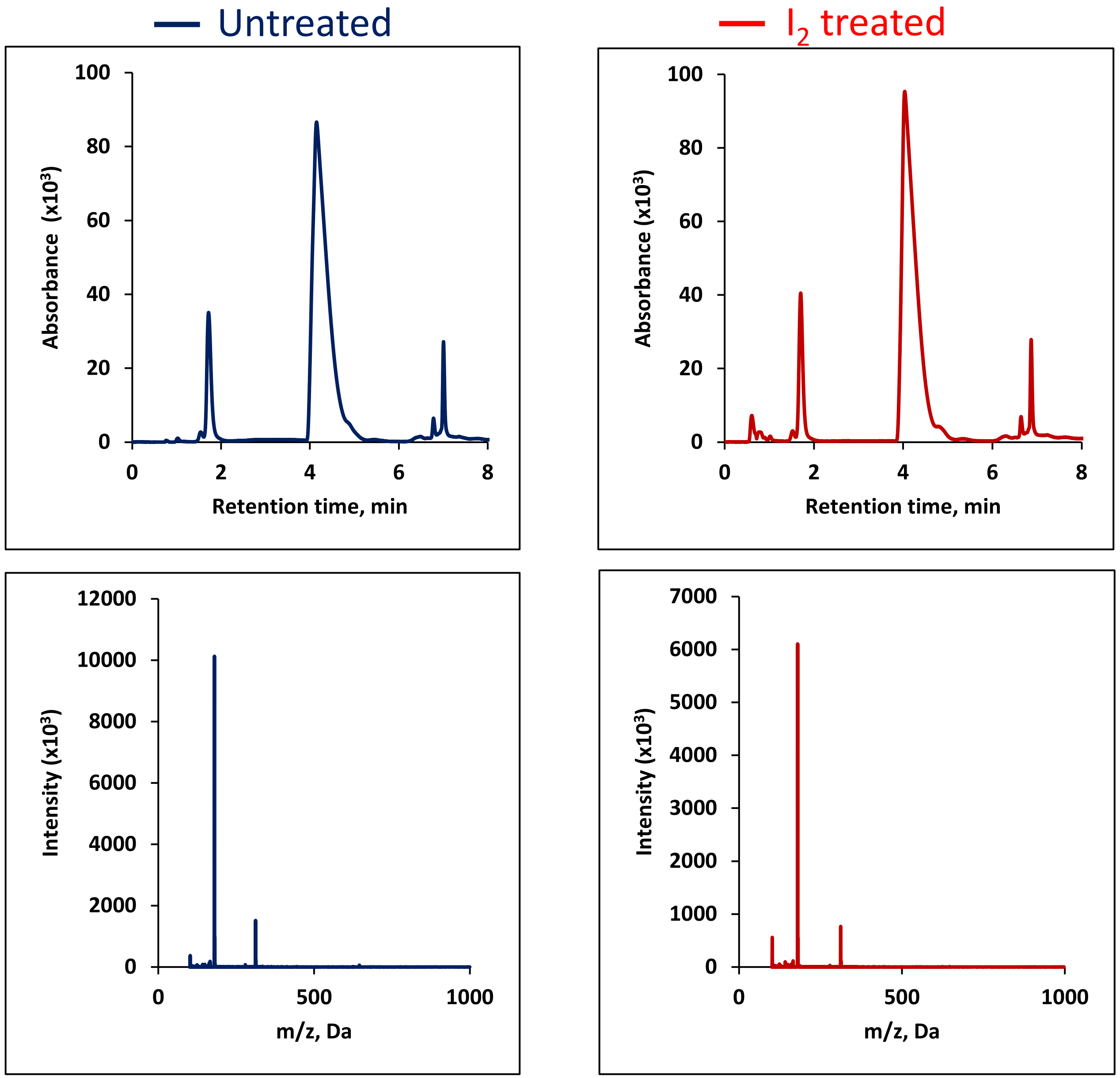

5-methyl-2-thiouridine ($m^5s_2U$)

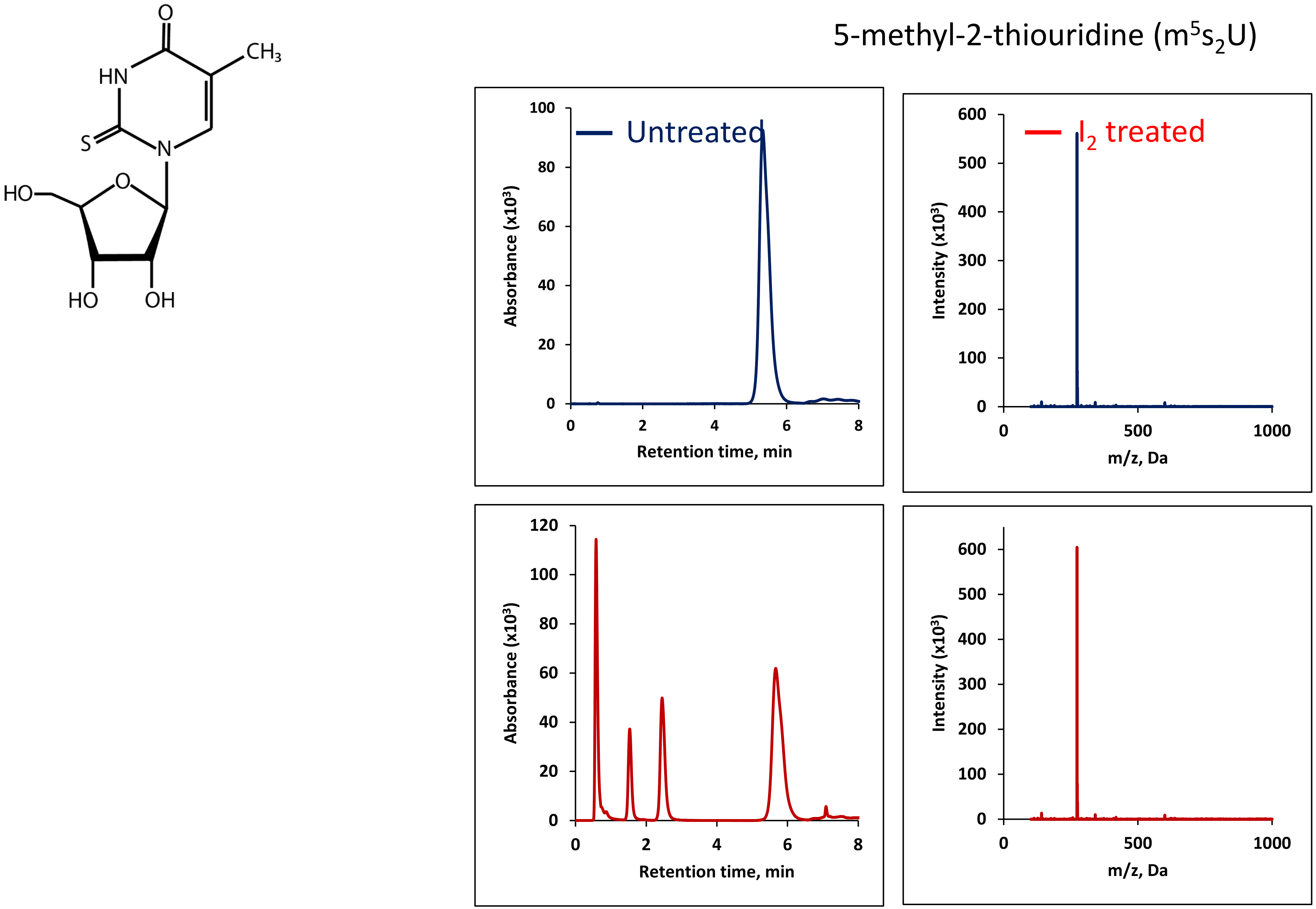

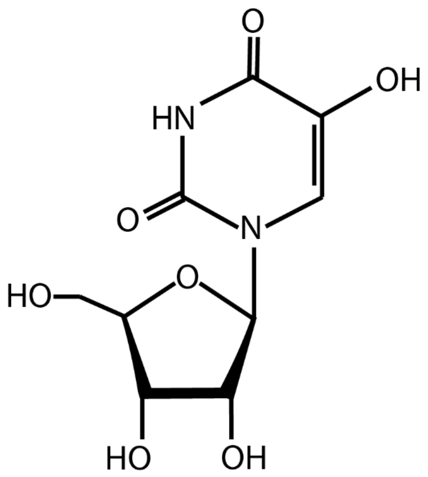


### 5-hydroxyuridine (ho5U)

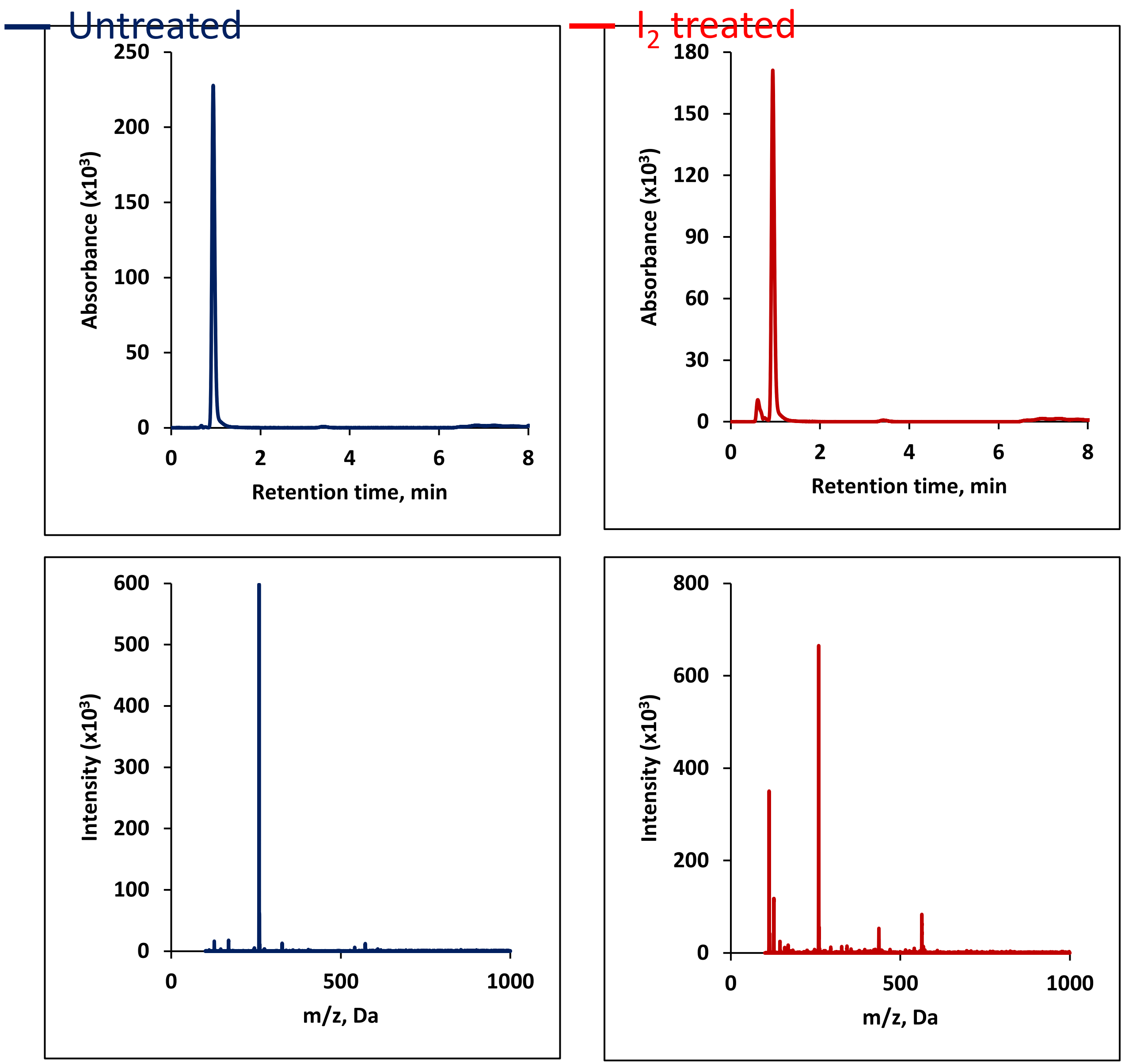

### Pseudouridine (ψ)

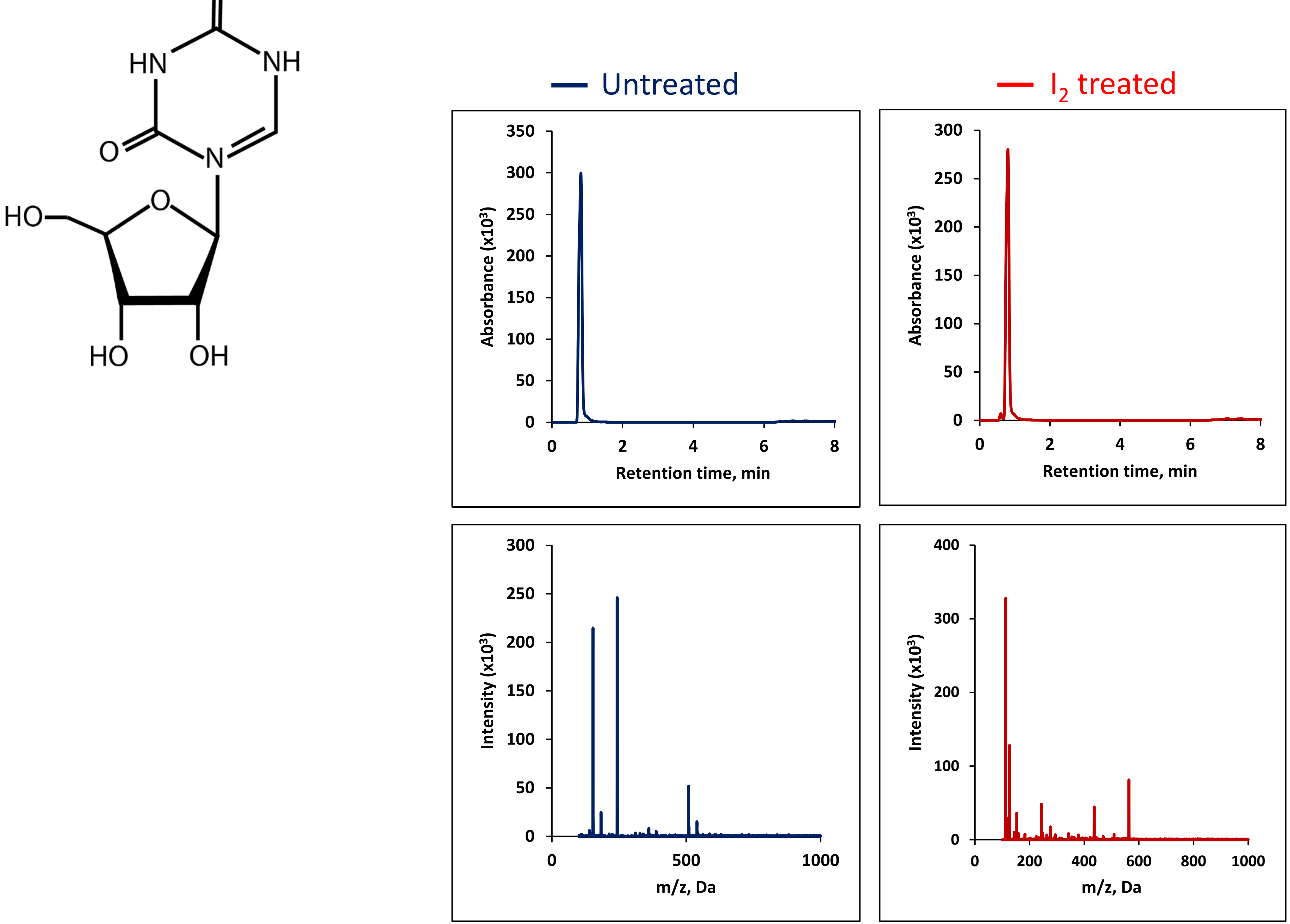

### N2,2-Dimethylguanosine (m22G)

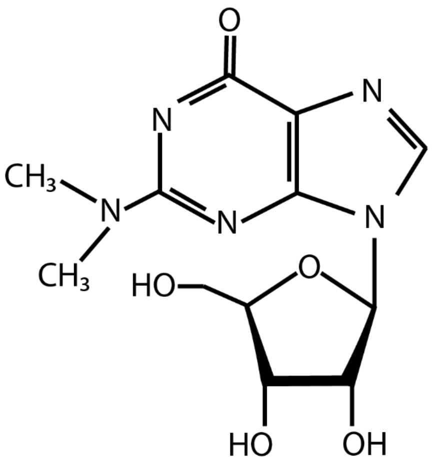


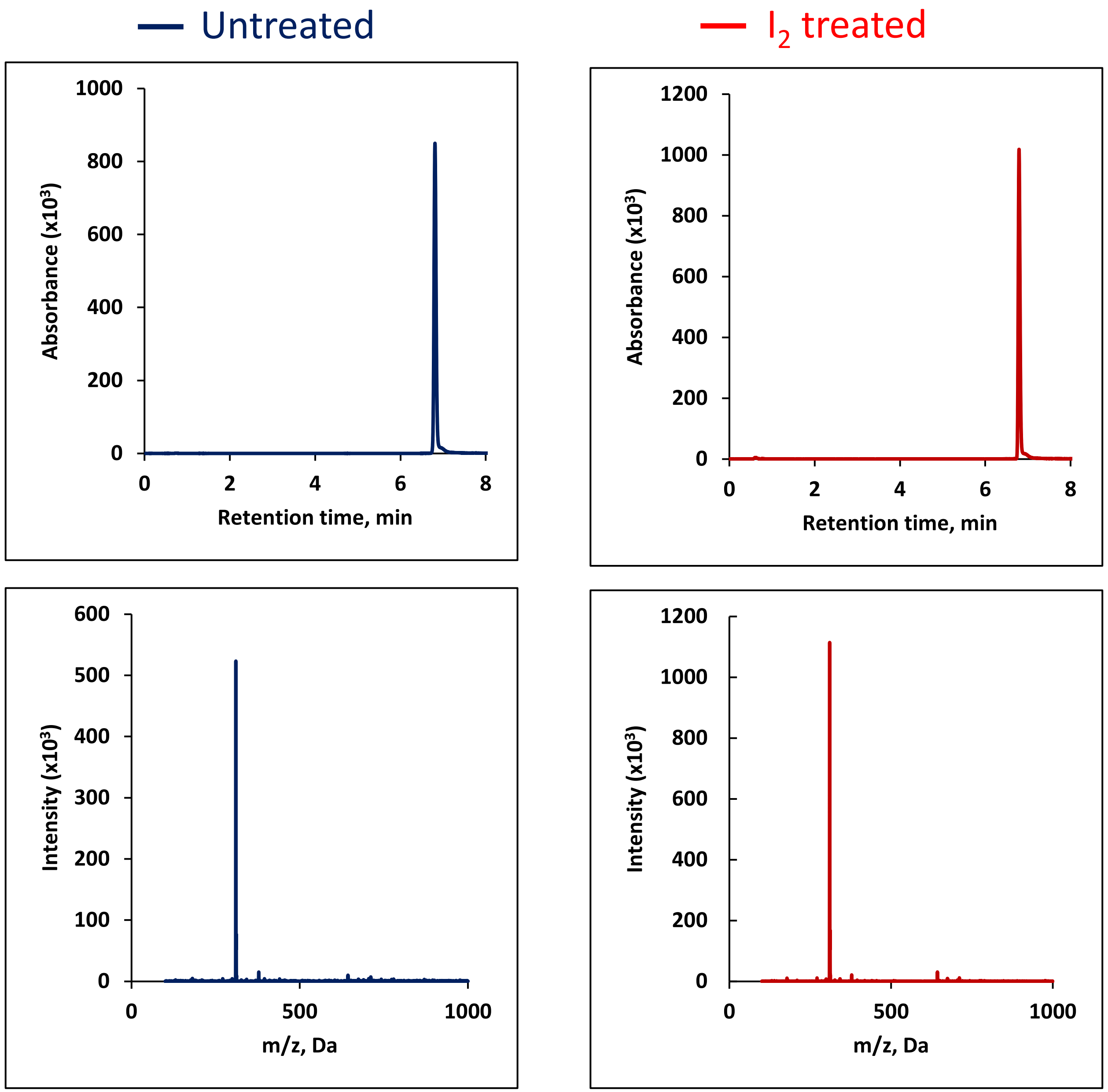

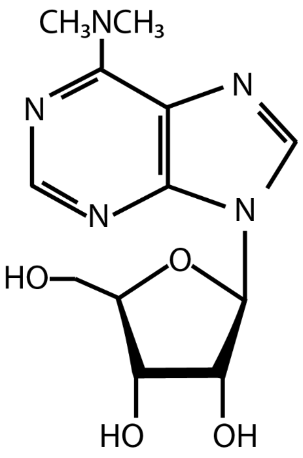


### N6,6-Dimethyladenosine ($m^6{}_2A$)

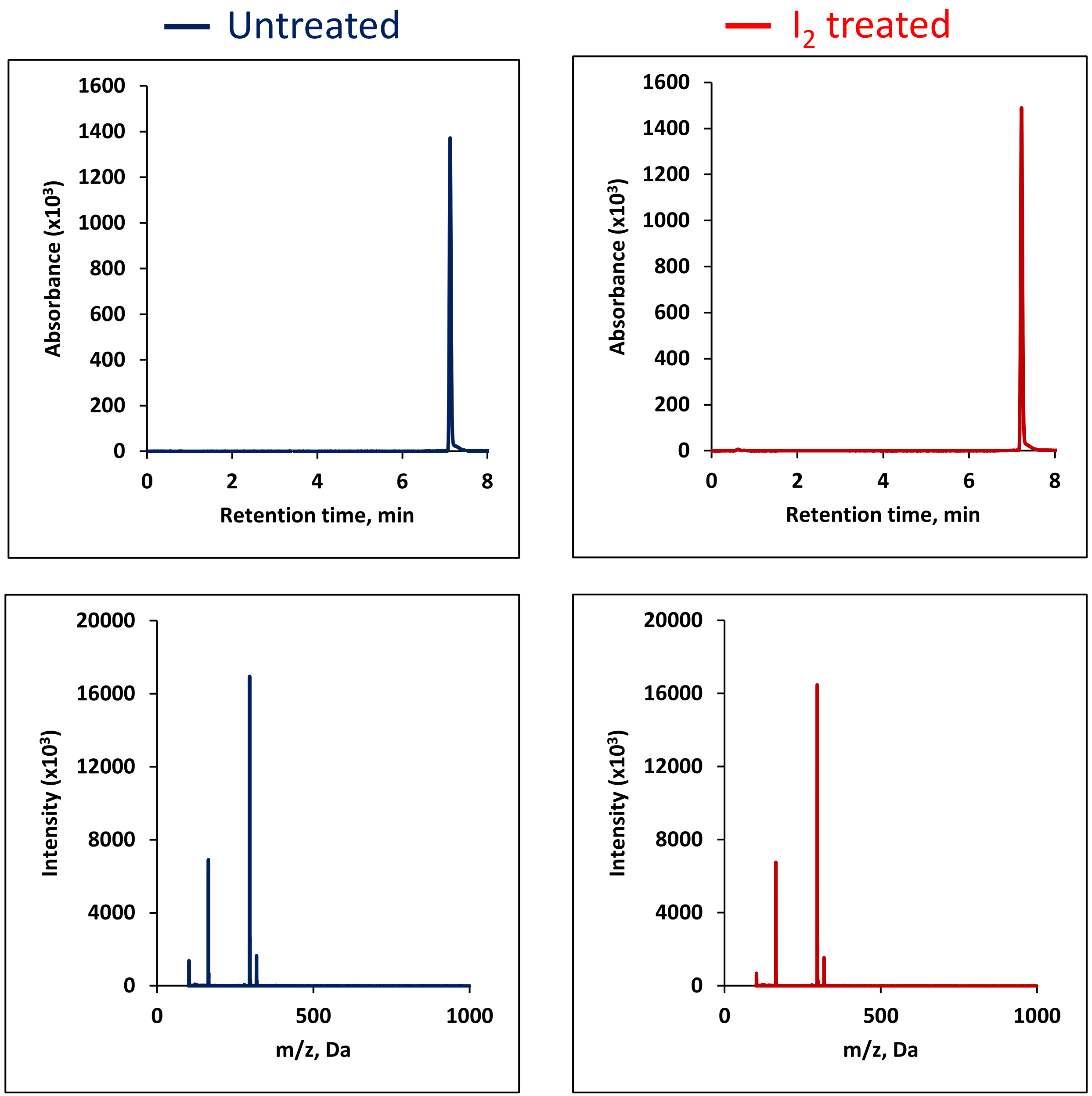

### 2-methylthio-$N^6$-isopentenyladenosine ($ms_2i^6A$)

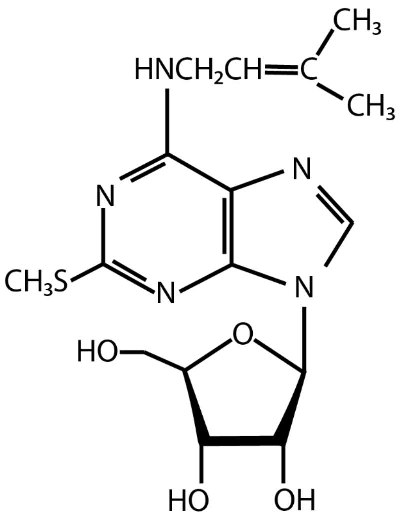

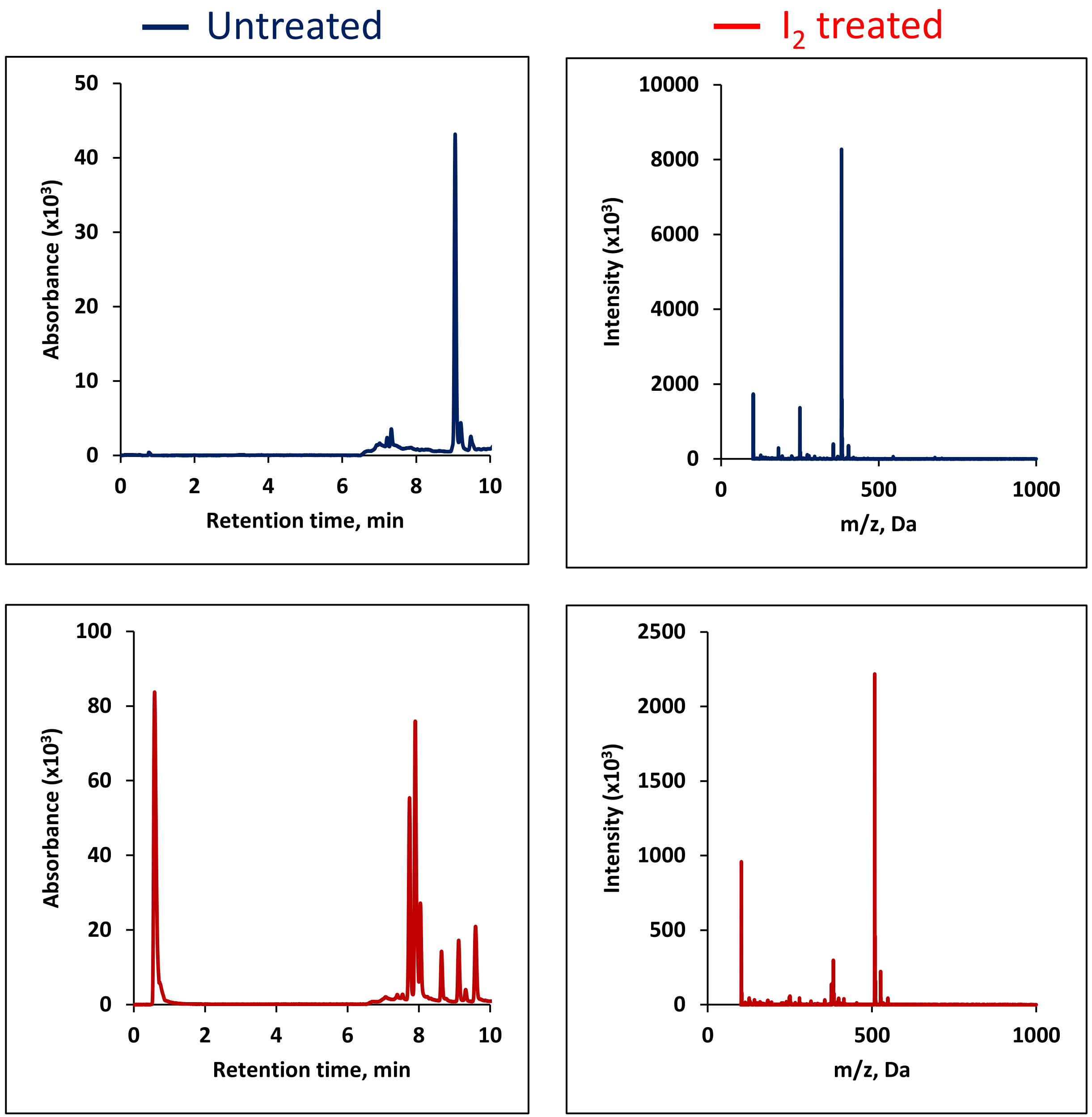

### 2′-*O*-Methylguanosine (Gm)

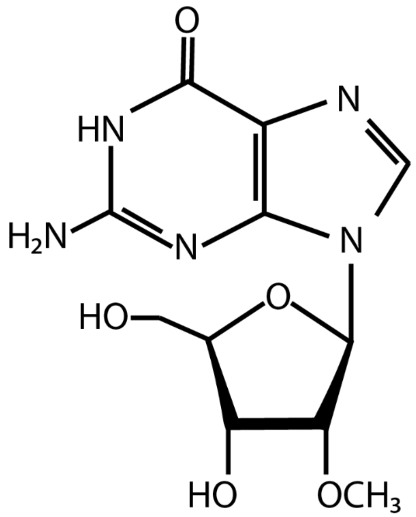


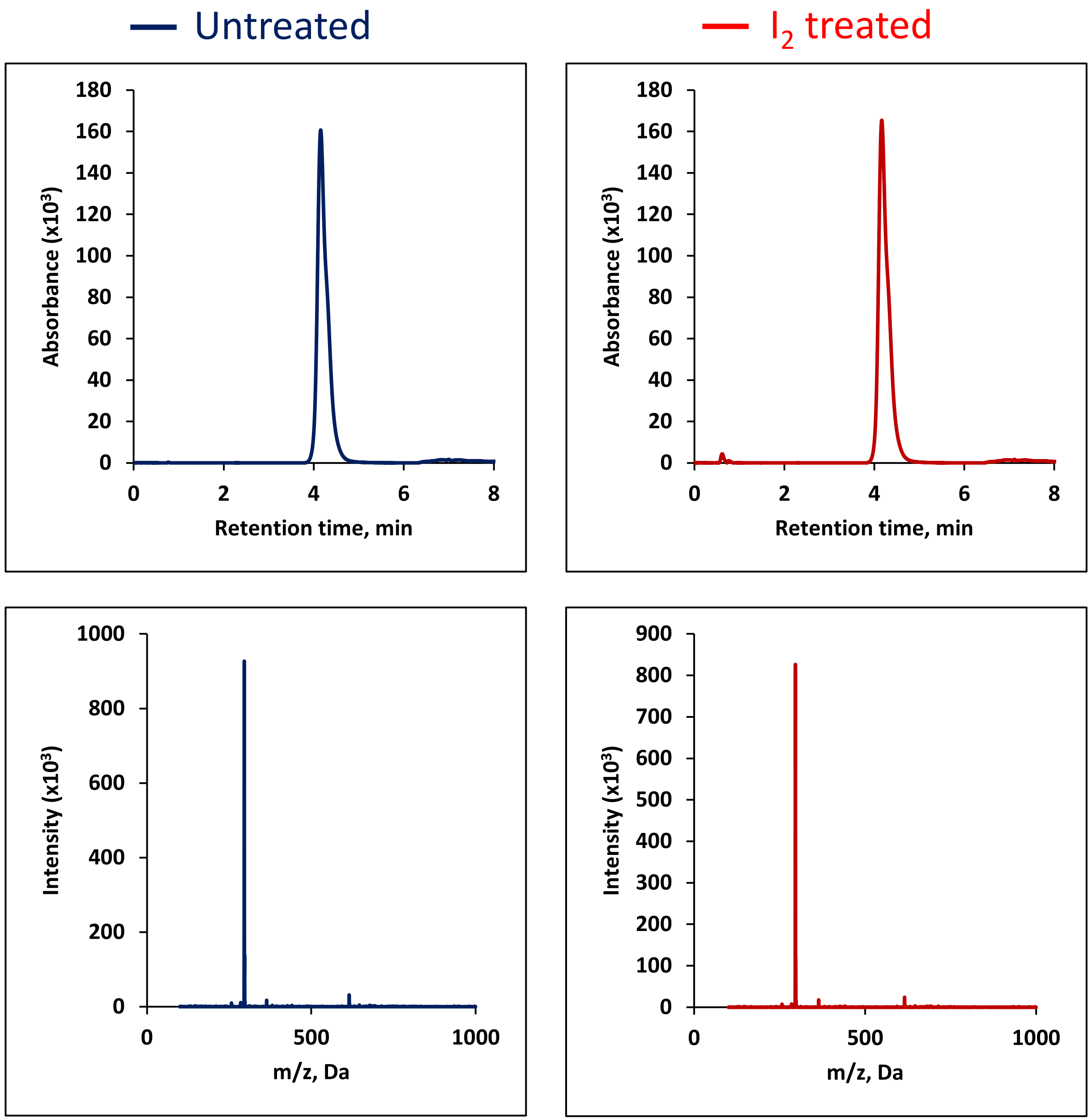

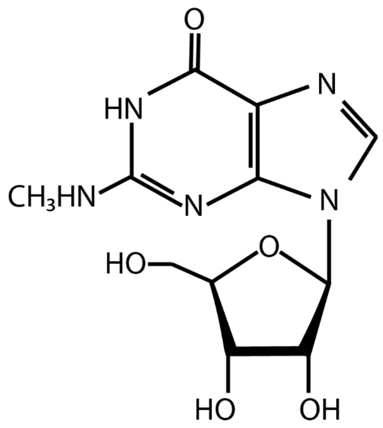


### N2-Methylguanosine ($m^2G$)

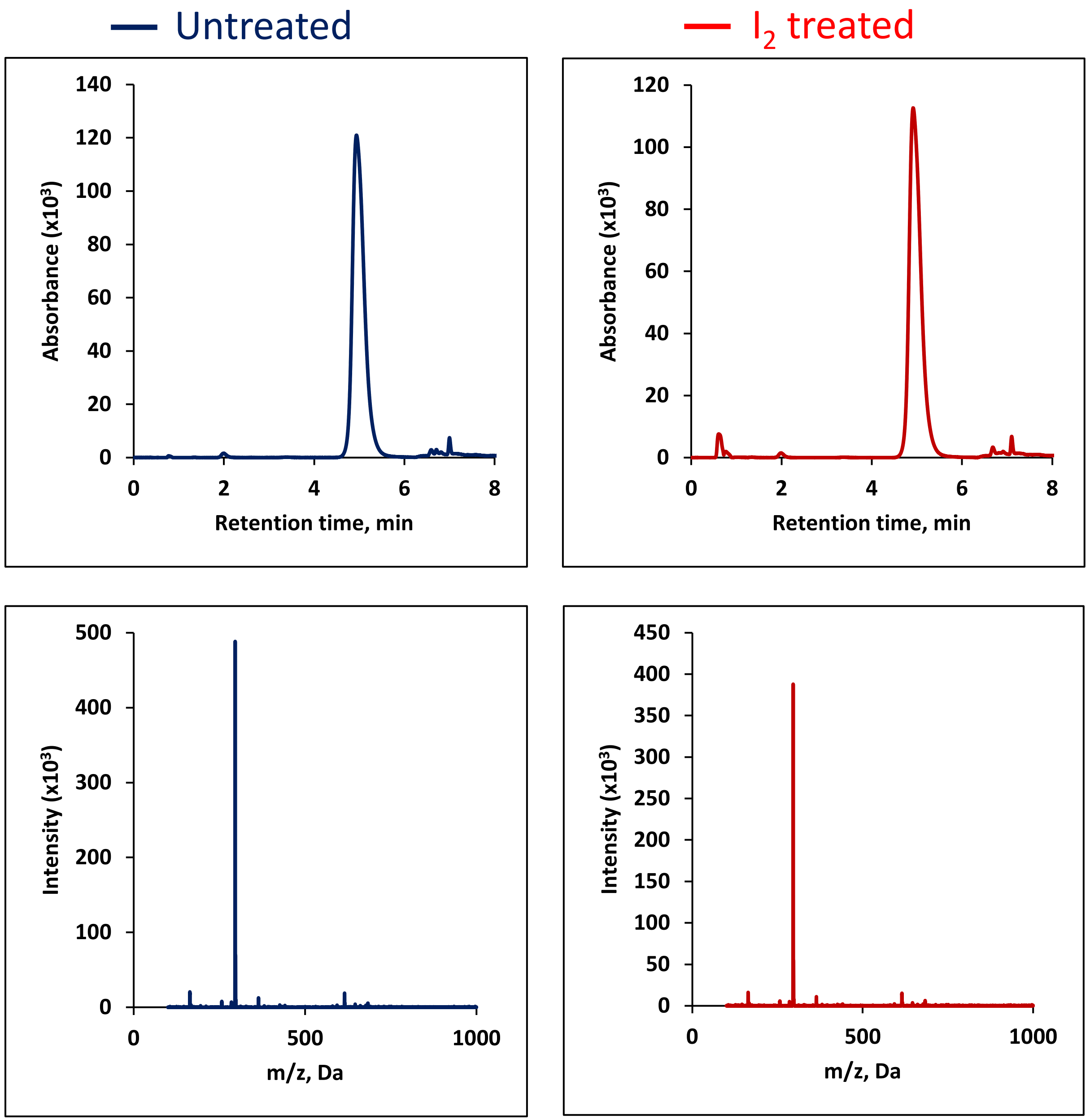

### 2′-*O*-Methylcytidine (Cm)

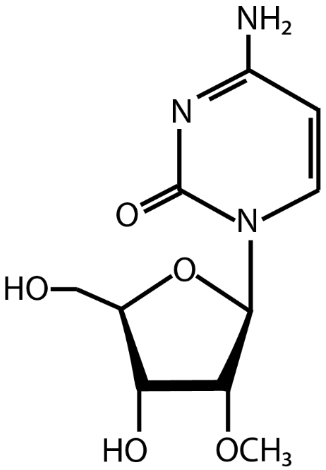


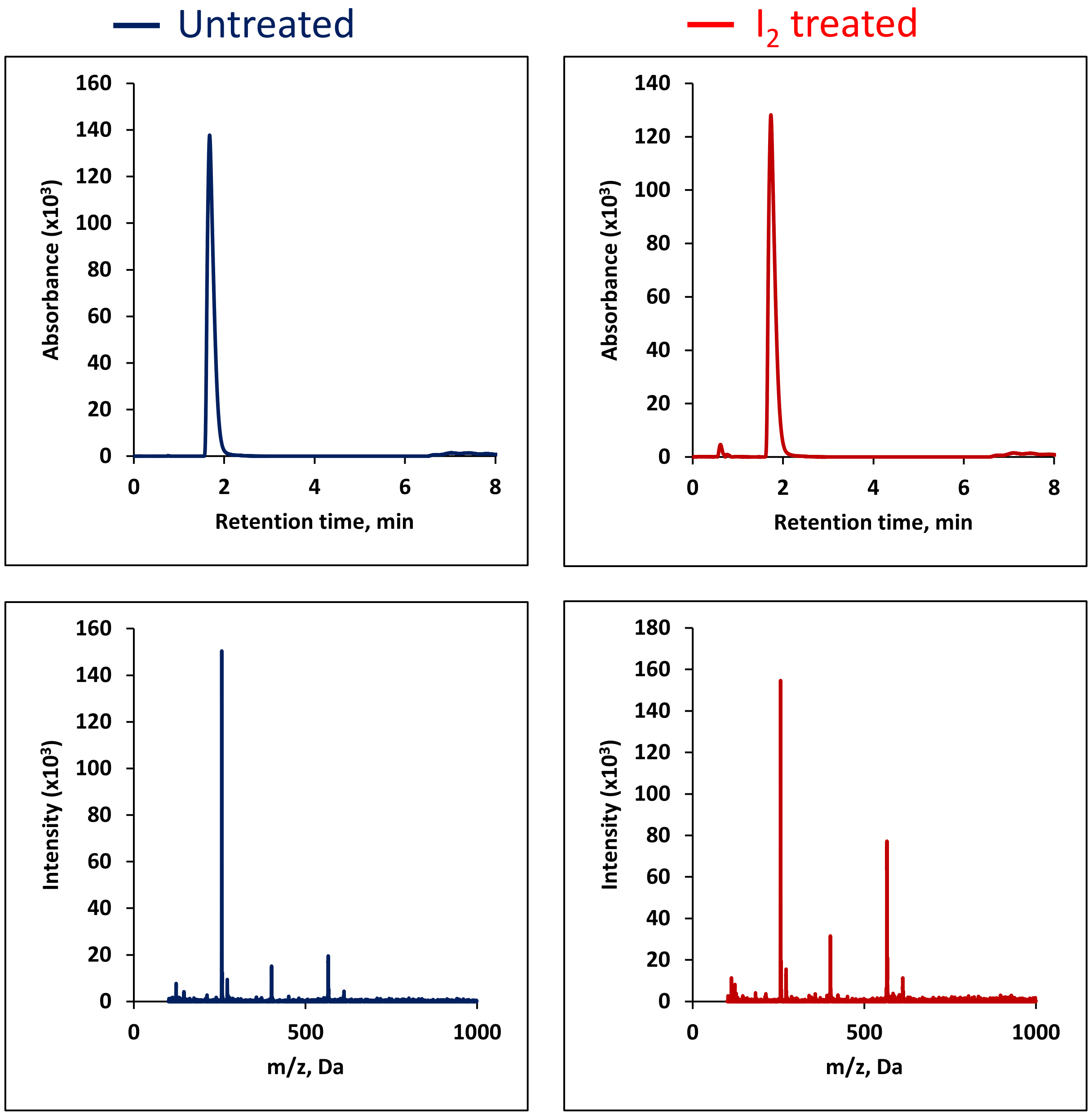

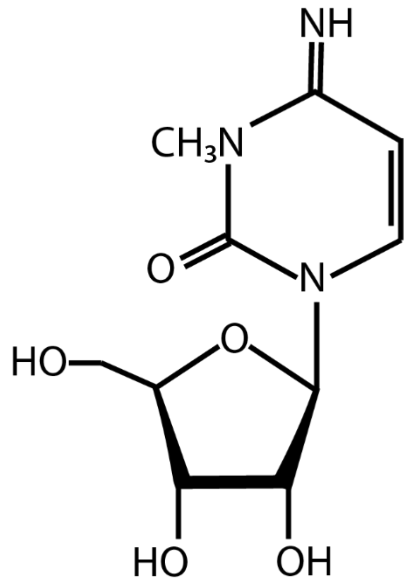


### 3-Methylcytidine ($m^3C$)

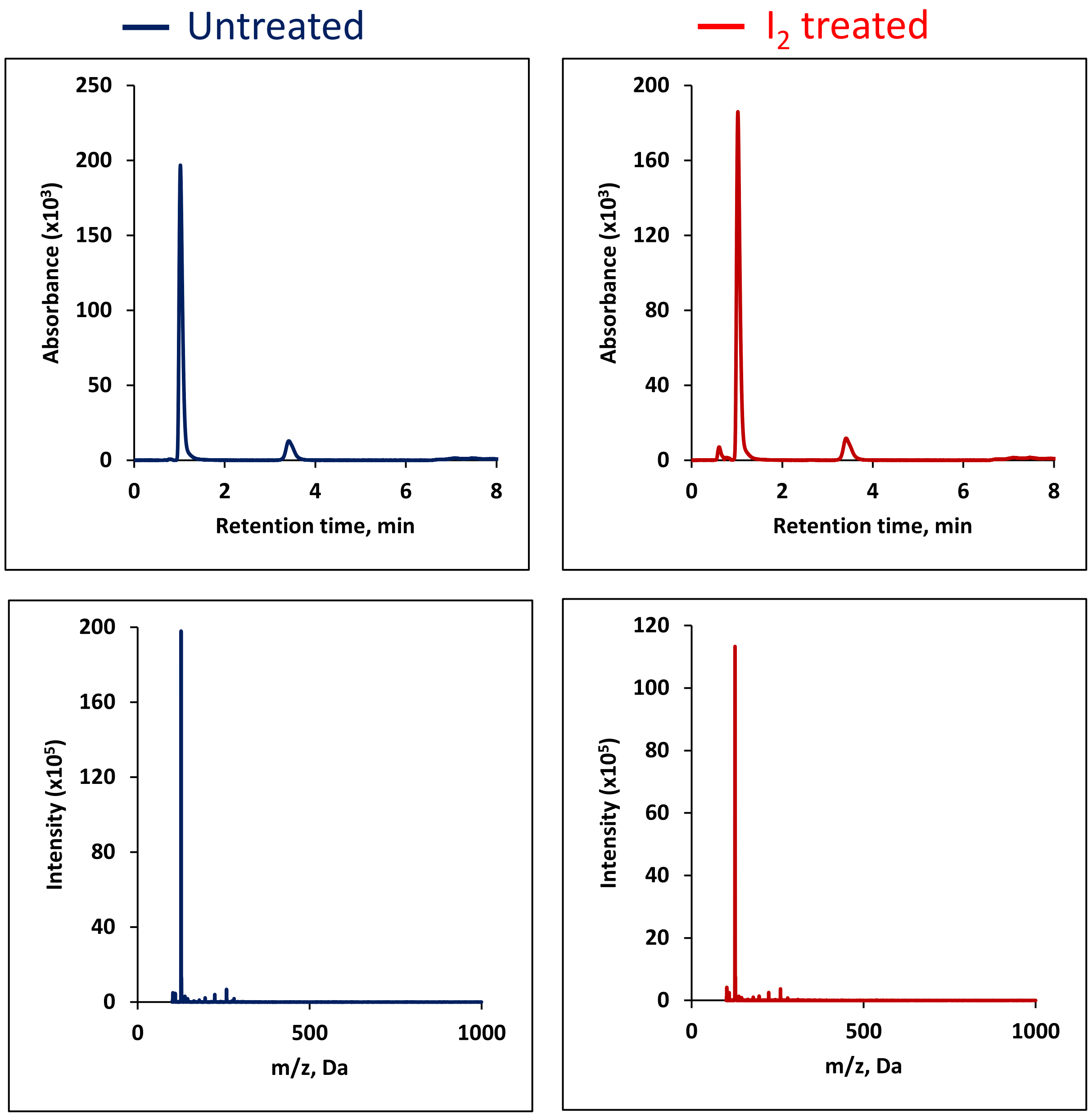

### 2′-O-Methyluridine (Um)

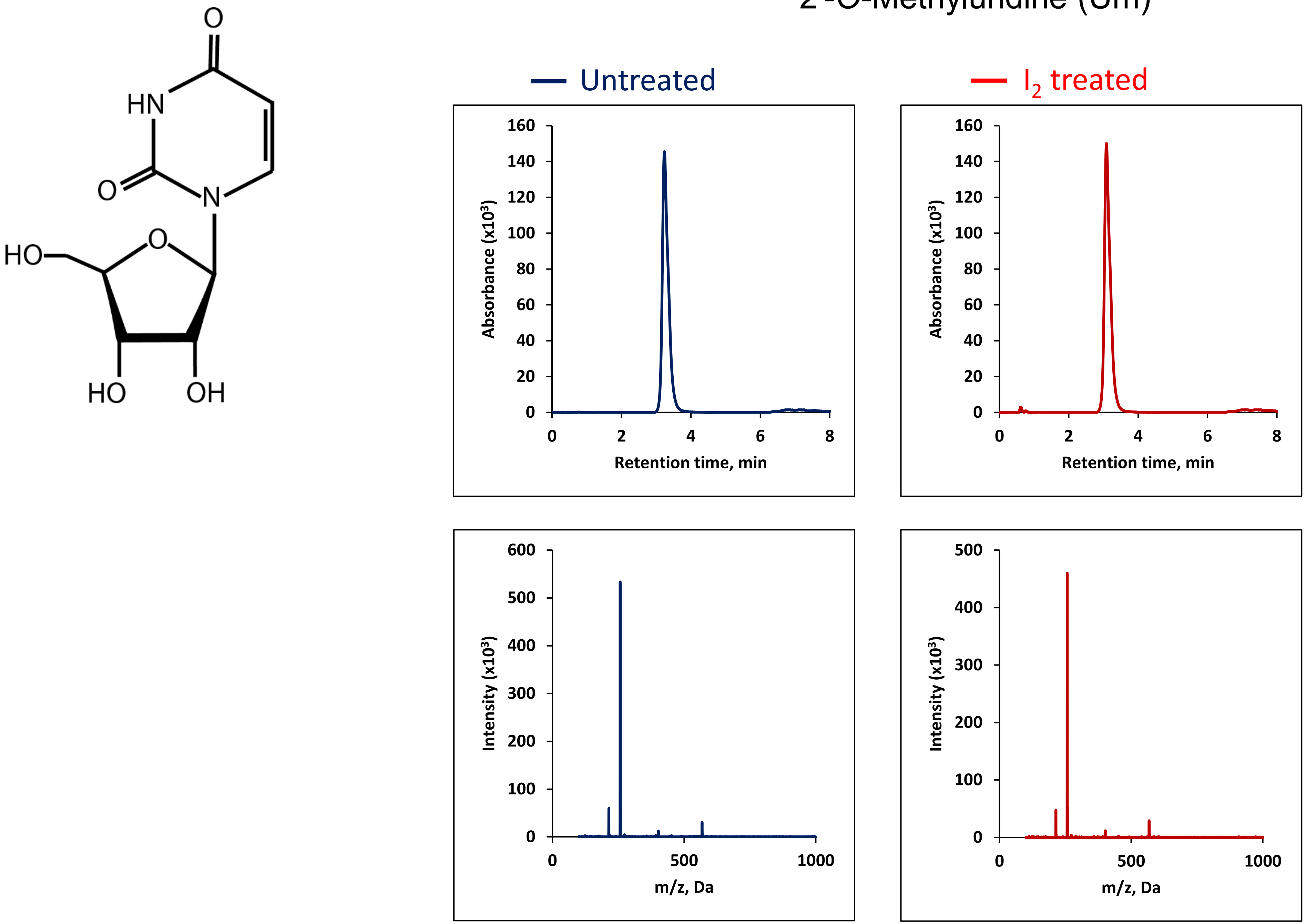

1,2'-*O*-dimethyladenosine ($m^1Am$)

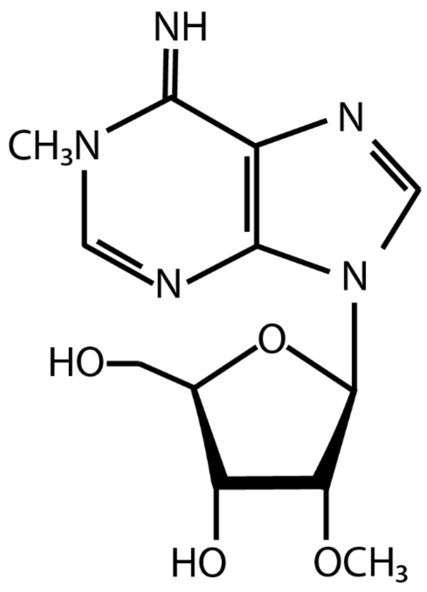


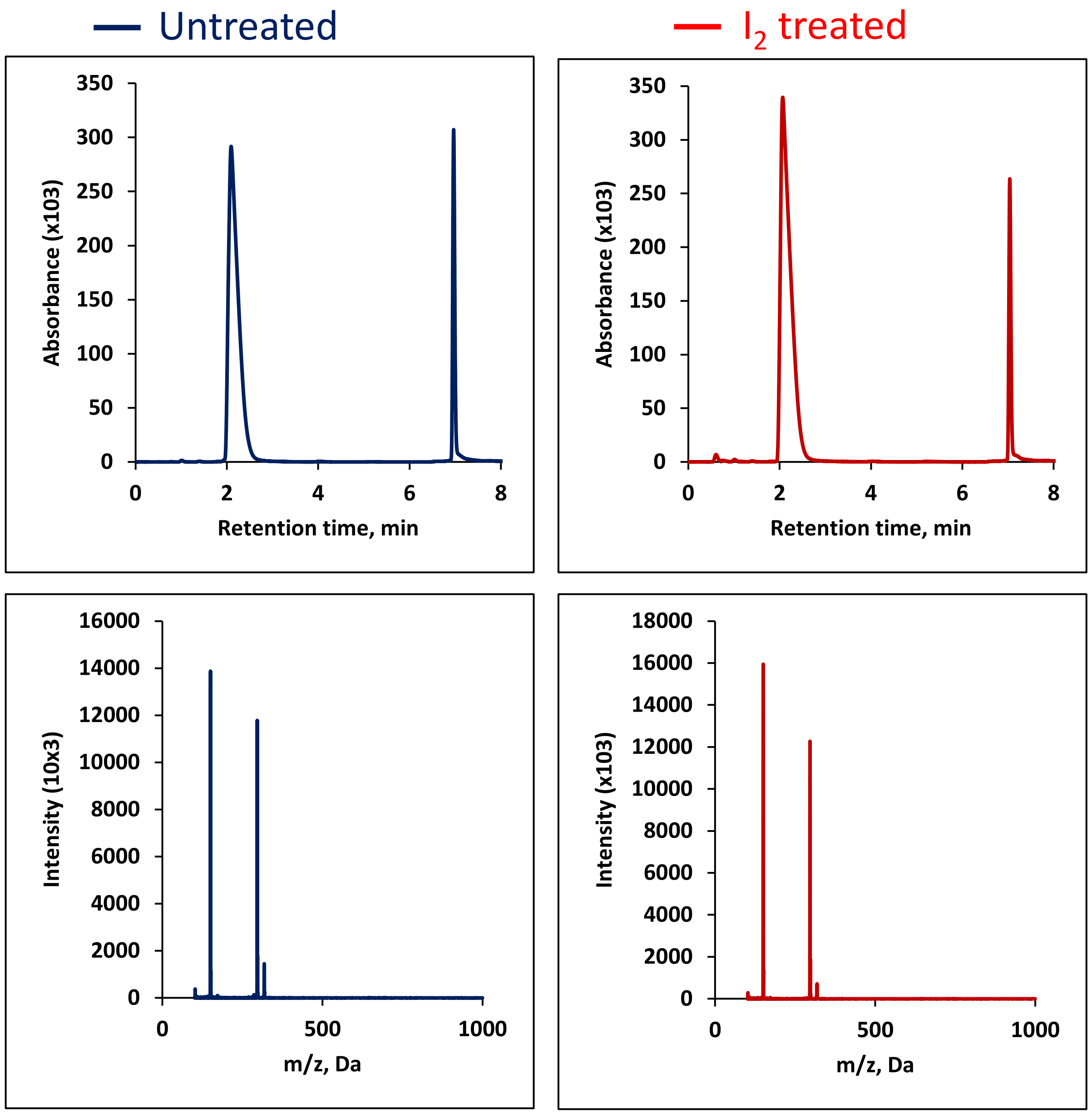

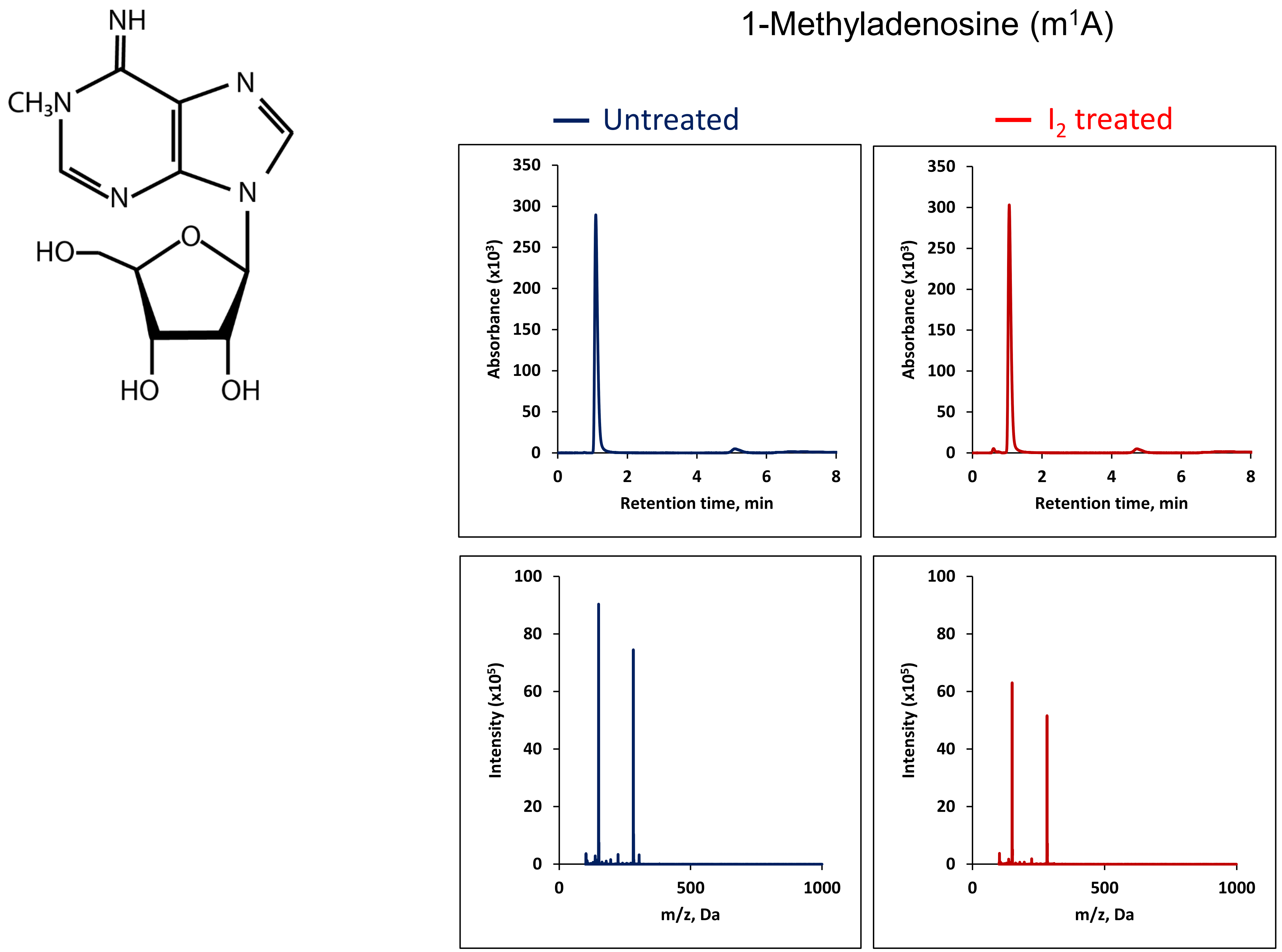
1-Methyladenosine (m1A)
NH
CH3N
N
N
N
HO
O
HO
OH
Untreated
I2 treated
Absorbance (x10³)
Retention time, min
350
300
250
200
150
100
50
0
0
2
4
6
8
Intensity (x10⁵)
m/z, Da
100
80
60
40
20
0
0
500
1000

### $N^6$-methyladenosine ($m^6A$)

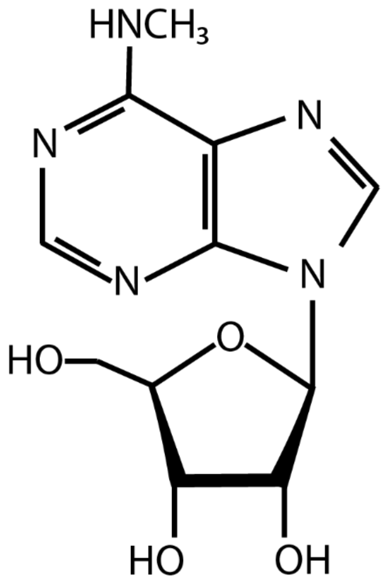


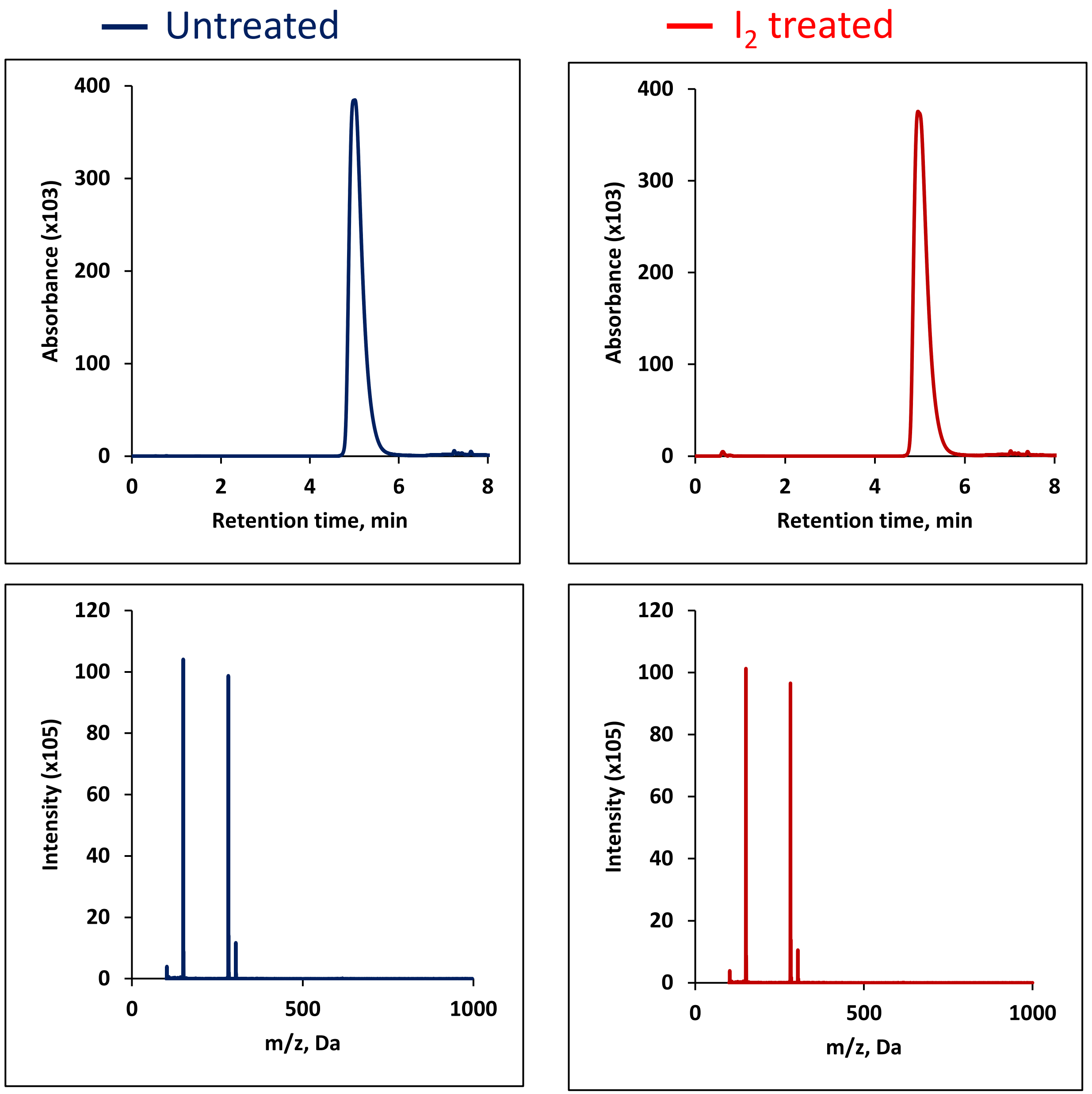

### 2′-*O*-Methyladenosine (Am)

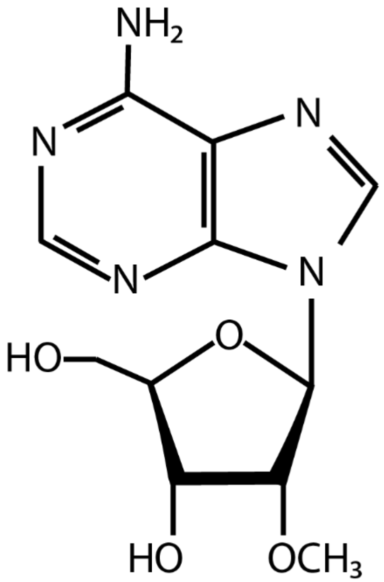


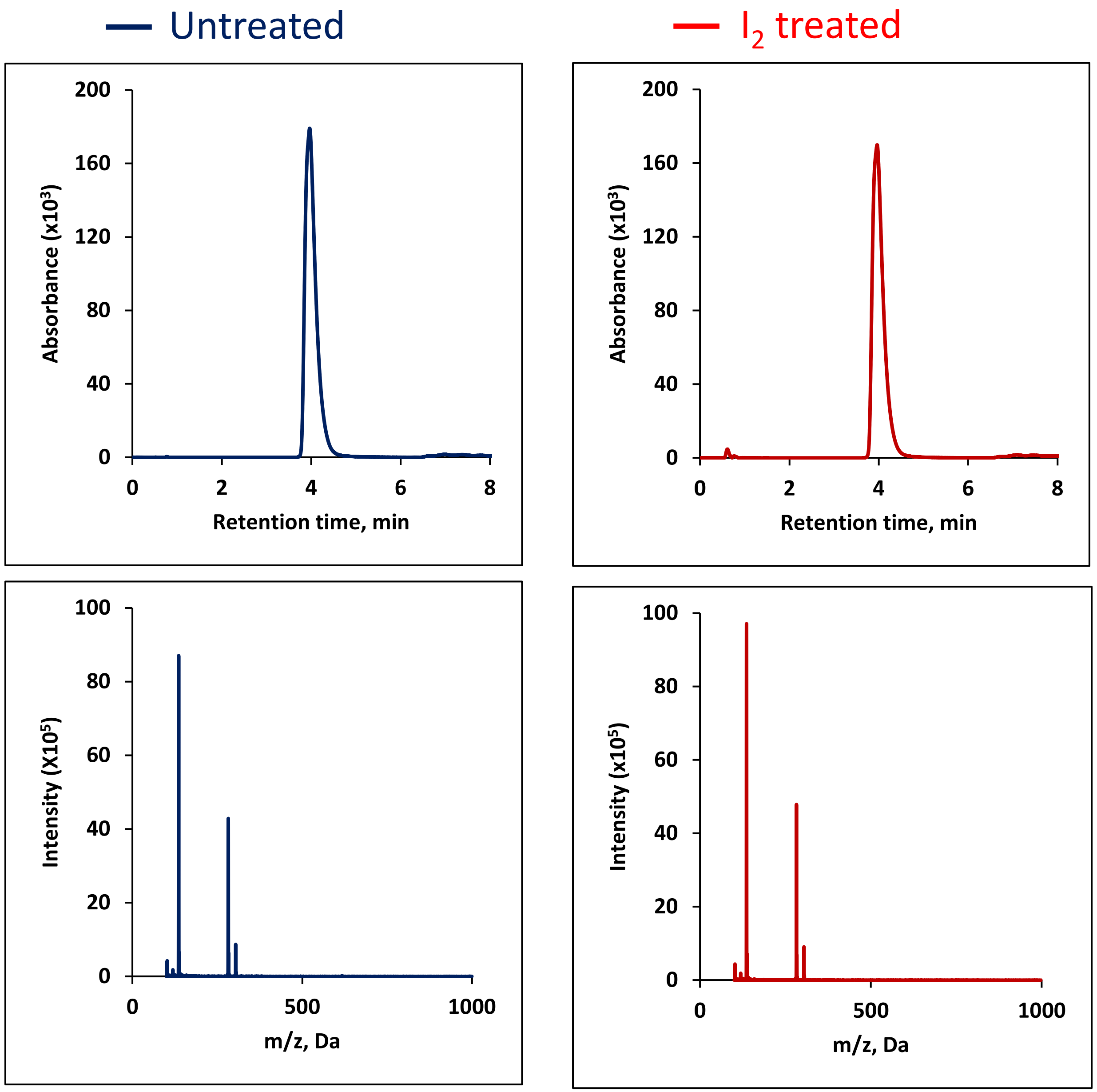

### 5-Methyluridine ($m^5U$)

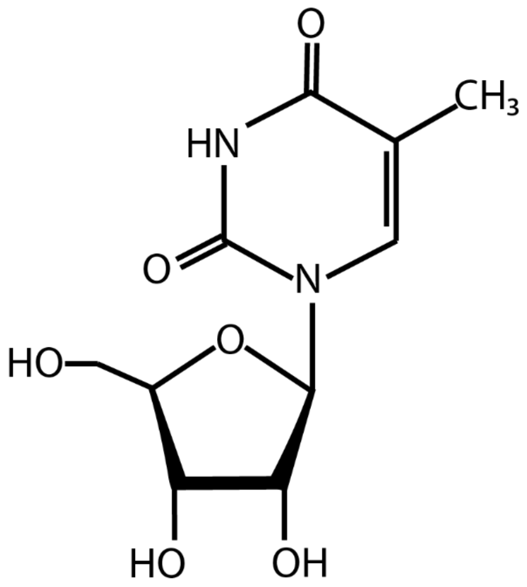


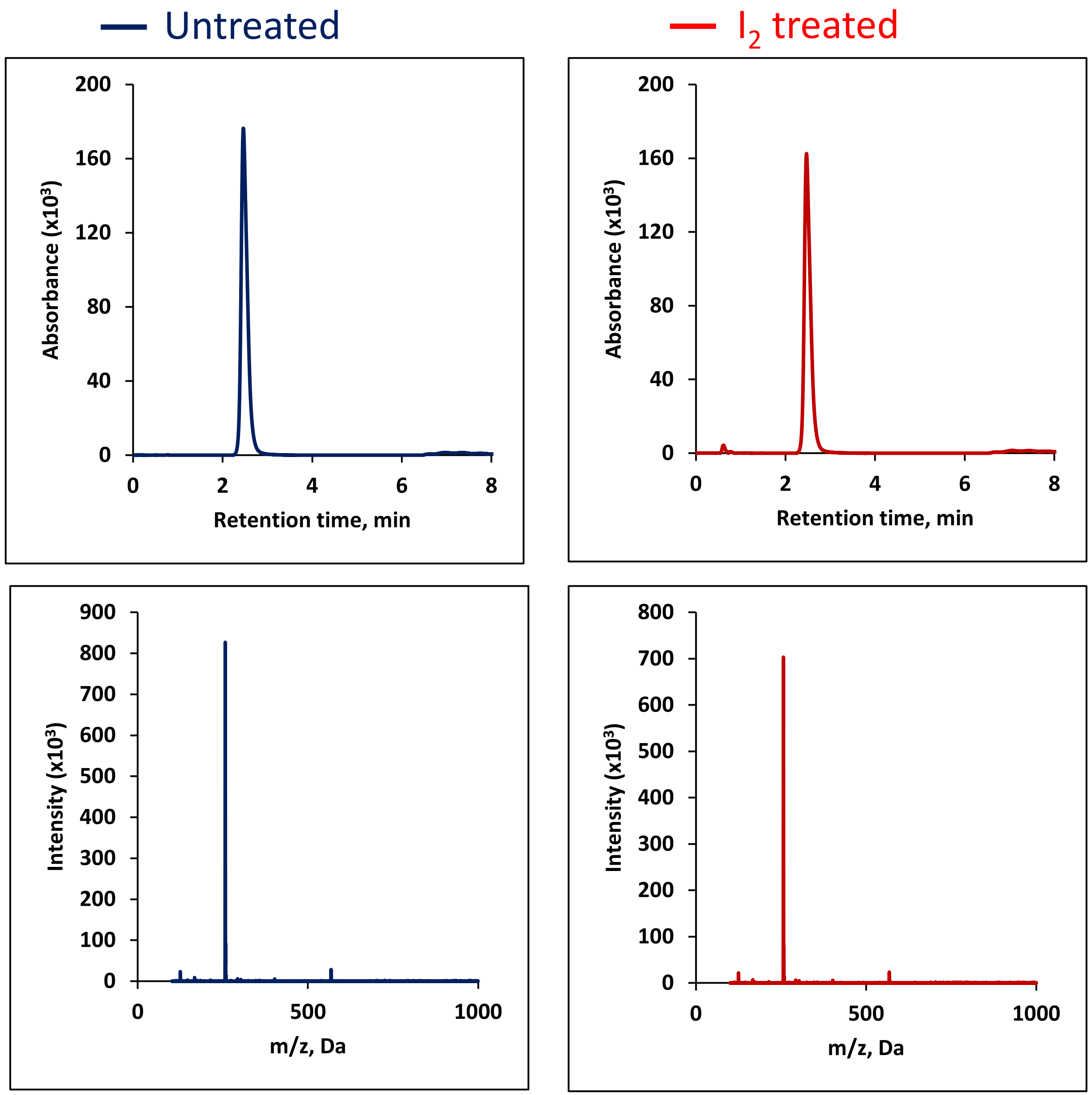

### 7-Methylguanosine ($m^7G$)

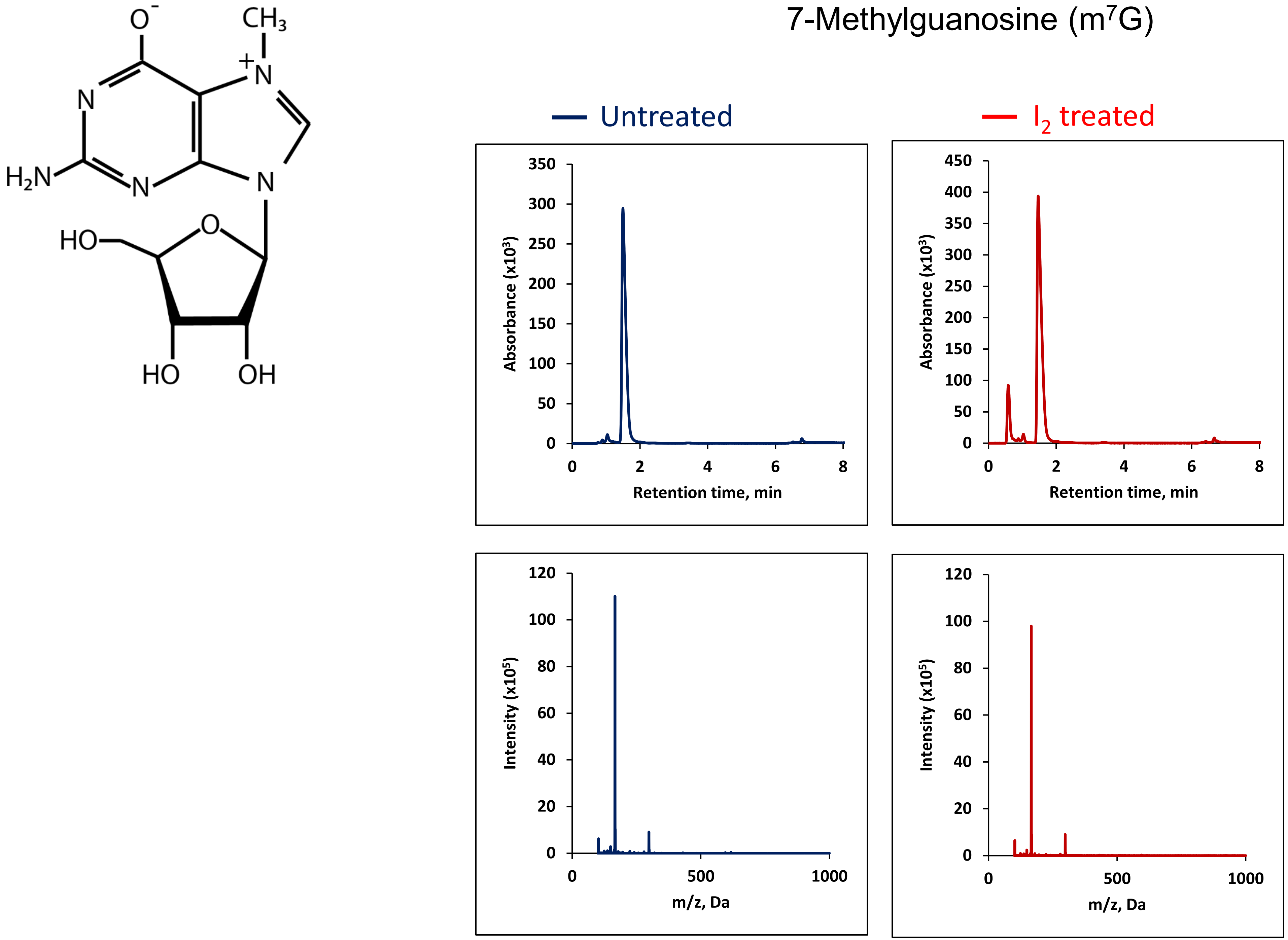

### 1-Methylguanosine ($m^1G$)

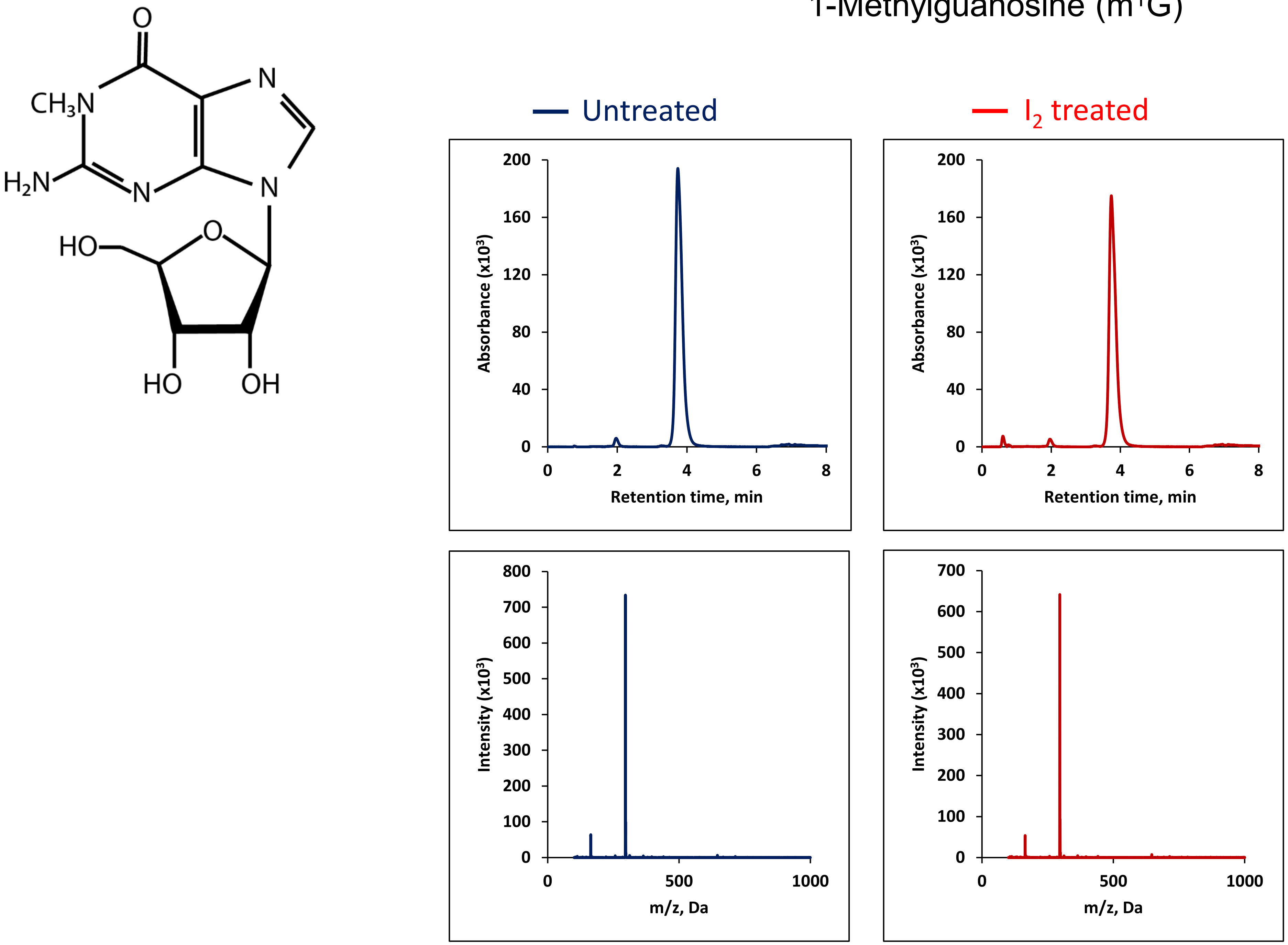

### 2-thiouridine ($s^2U$)

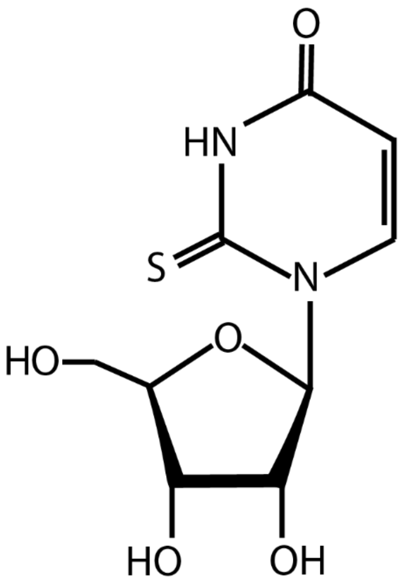


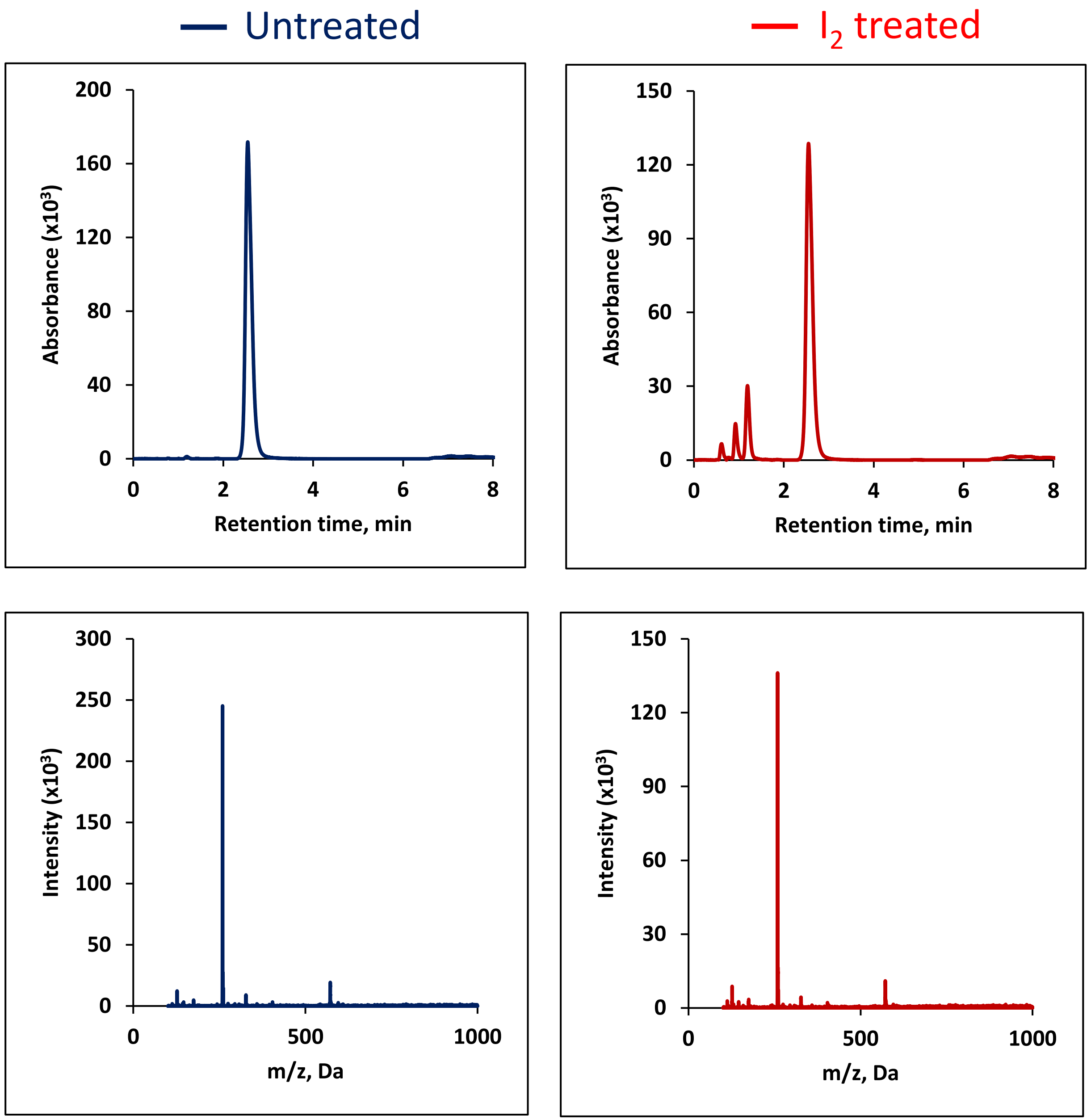

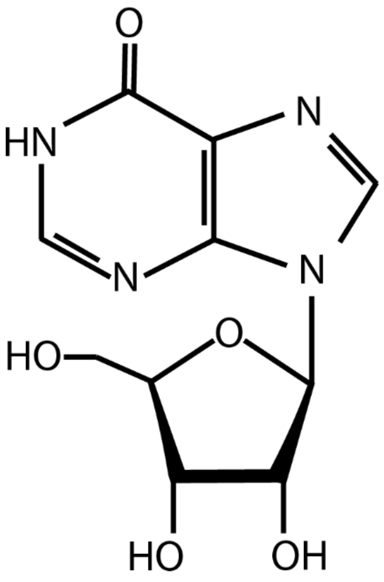
O
HN
N
N
N
HO
O
HO
OH

### Inosine (I)

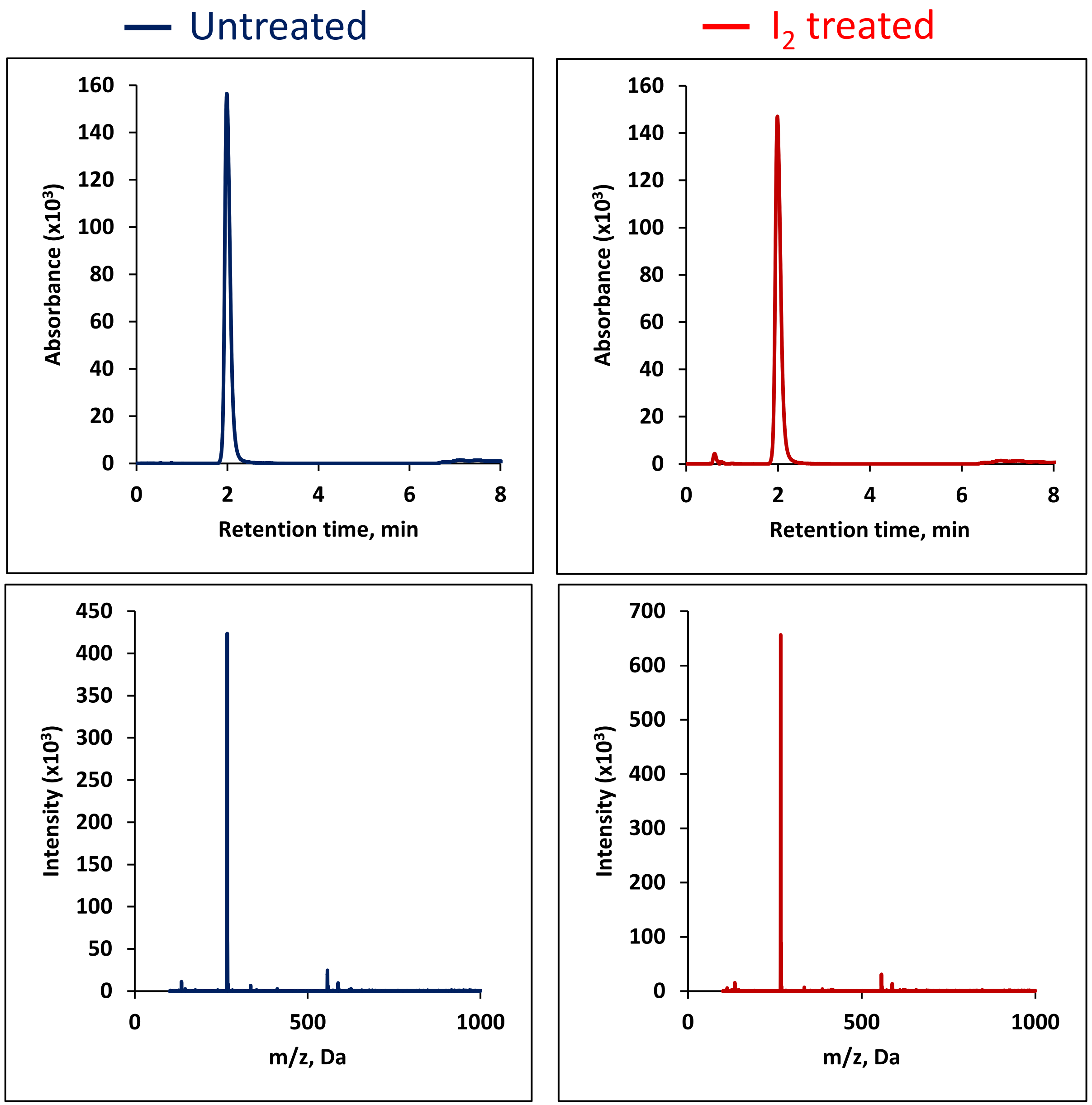
Untreated
I2 treated
Absorbance (x10³)
Retention time, min
Intensity (x10³)
m/z, Da
160
140
120
100
80
60
40
20
0
2
4
6
8
450
400
350
300
250
200
150
100
50
500
1000
700
600

### N6-Isopentenyladenosine ($i^6A$)

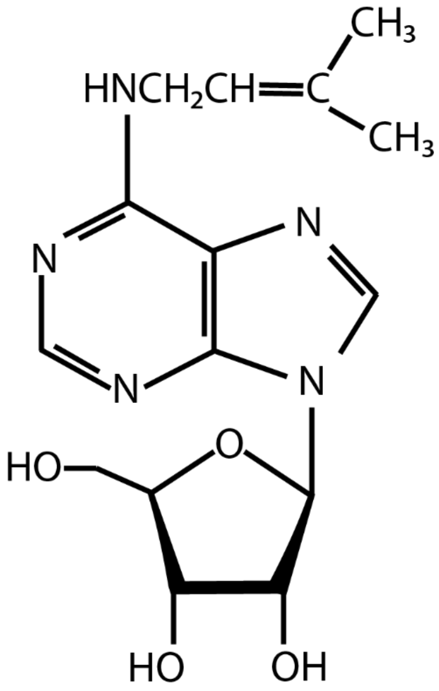


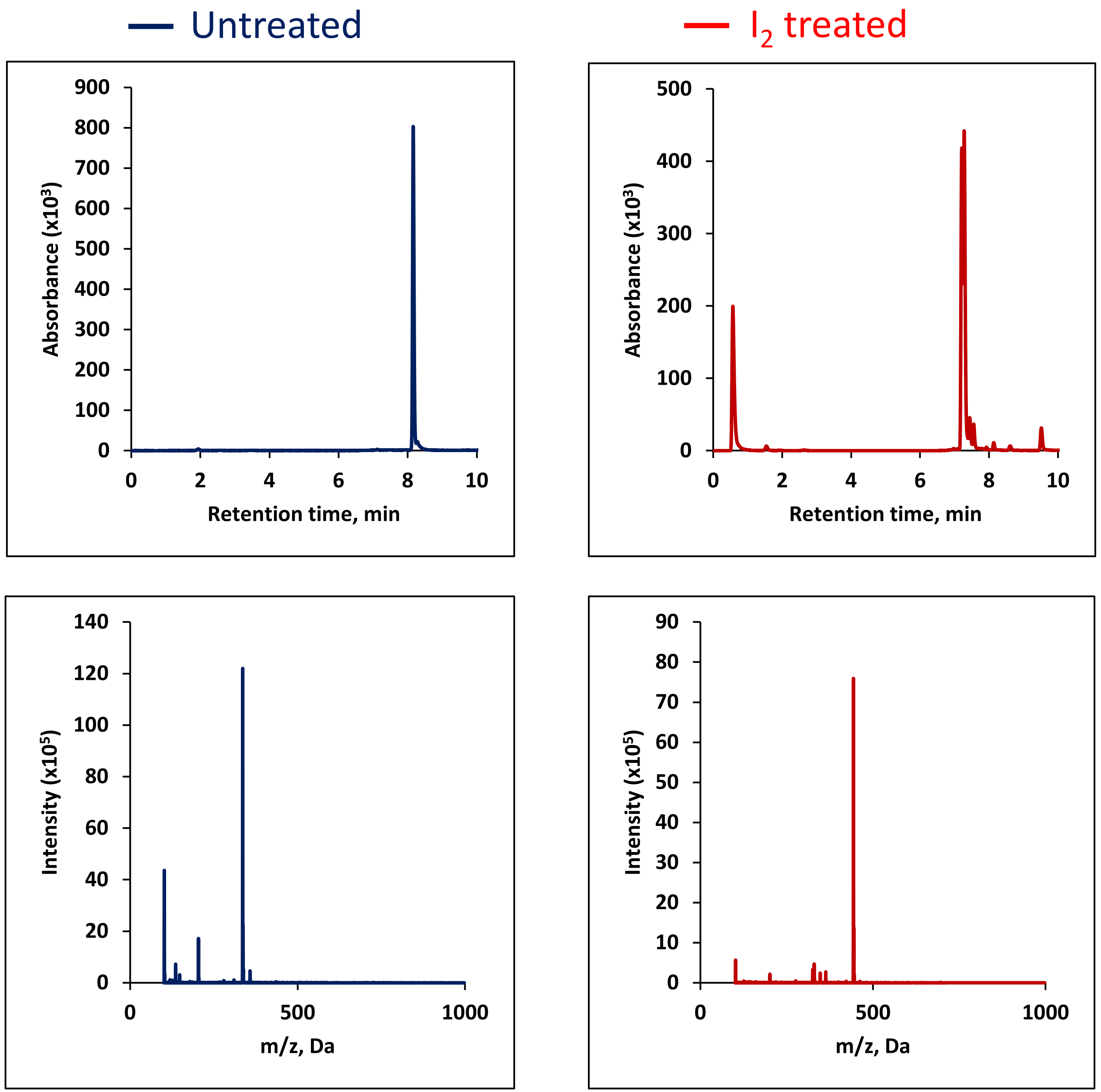

### 1-Methylinosine ($m^1I$)

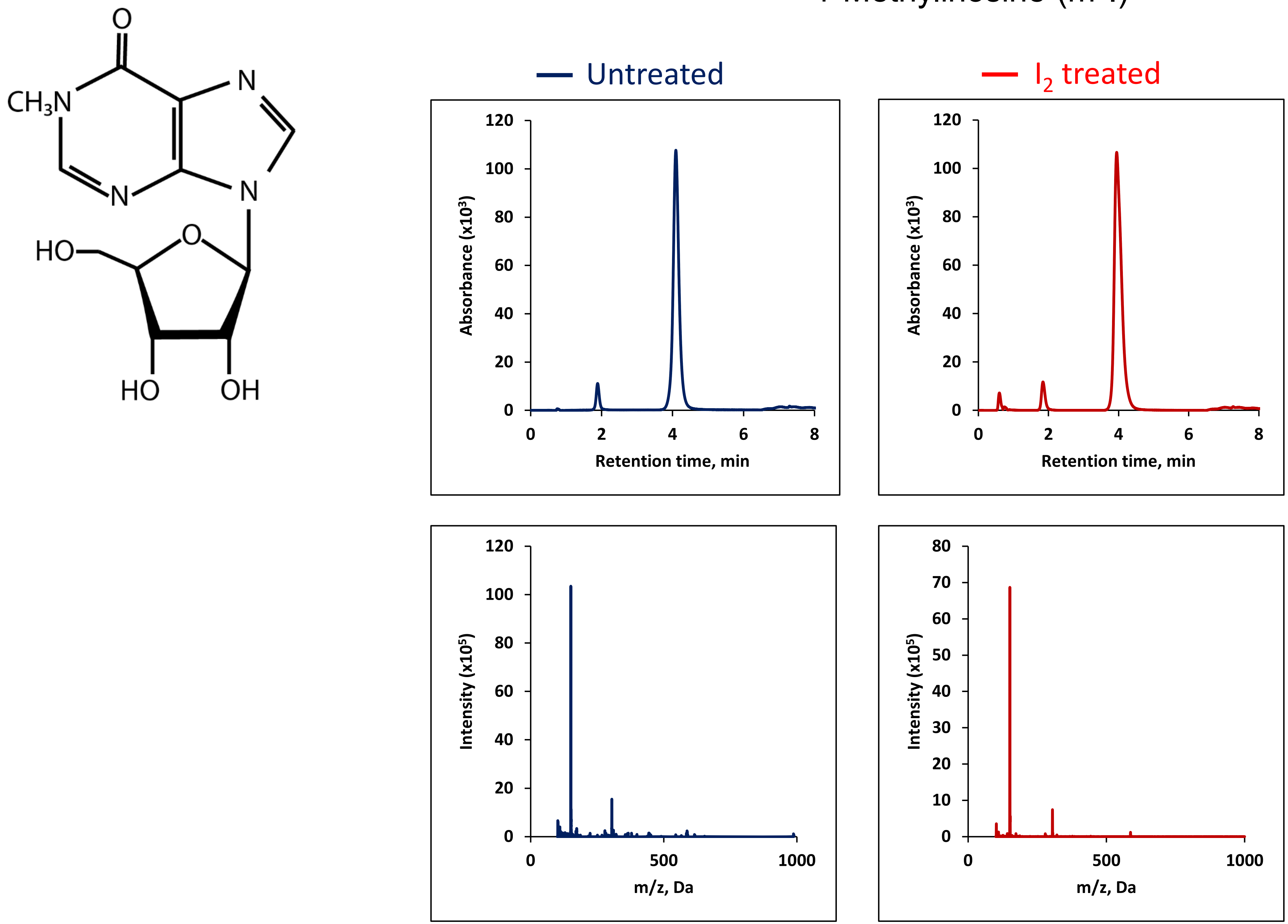

### 3-methyluridine (m$^3$U)

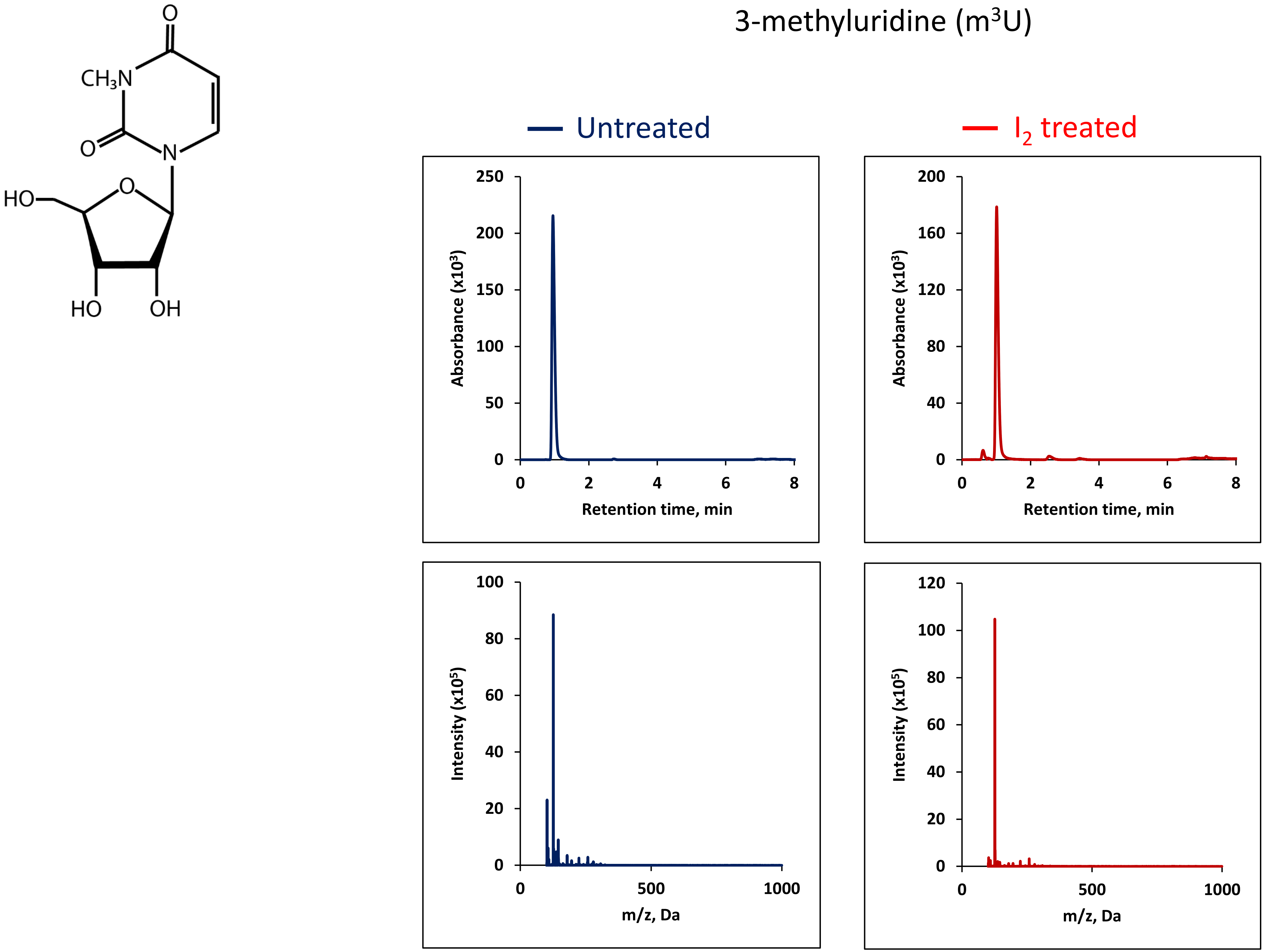

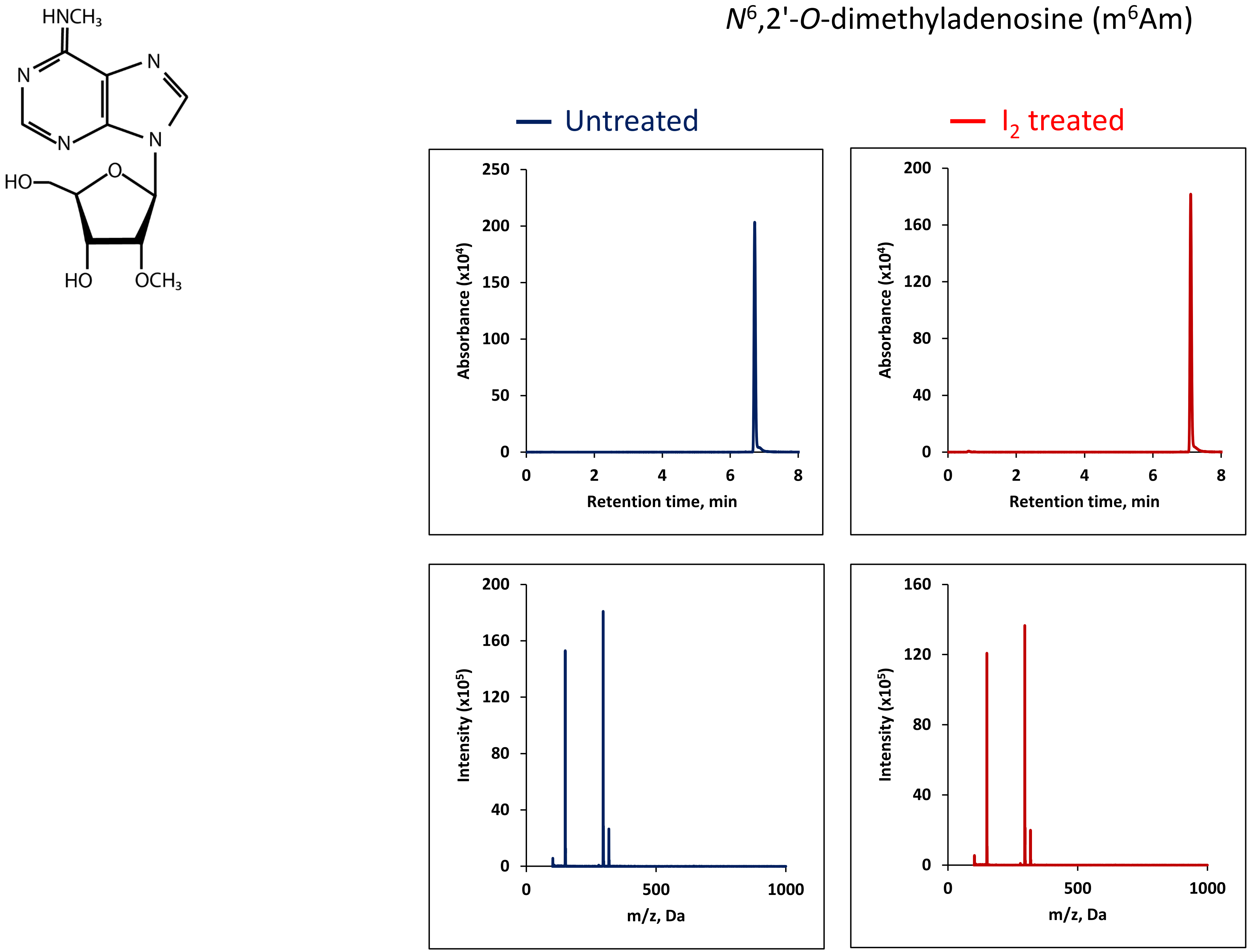
HNCH3
N
N
N
N
HO
O
HO
OCH3
N6,2'-O-dimethyladenosine (m6Am)
Untreated
I2 treated
250
200
150
100
50
0
Absorbance (x10⁴)
0
2
4
6
8
Retention time, min
200
160
120
80
40
0
Absorbance (x10⁴)
Retention time, min
Intensity (x10⁵)
500
1000
m/z, Da
Intensity (x10⁵)
m/z, Da

### 2-methylthio-$N^6$-threonyl carbamoyladenosine ($ms_2t^6A$)

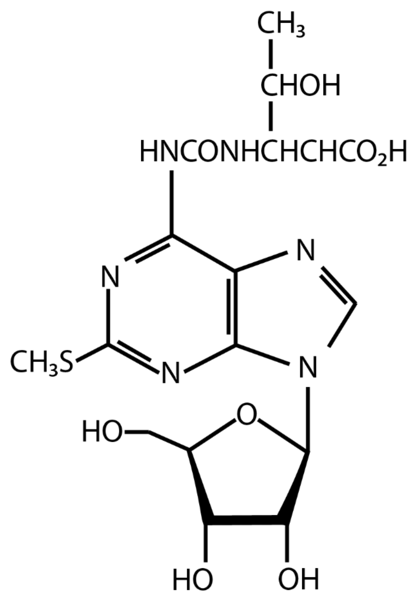


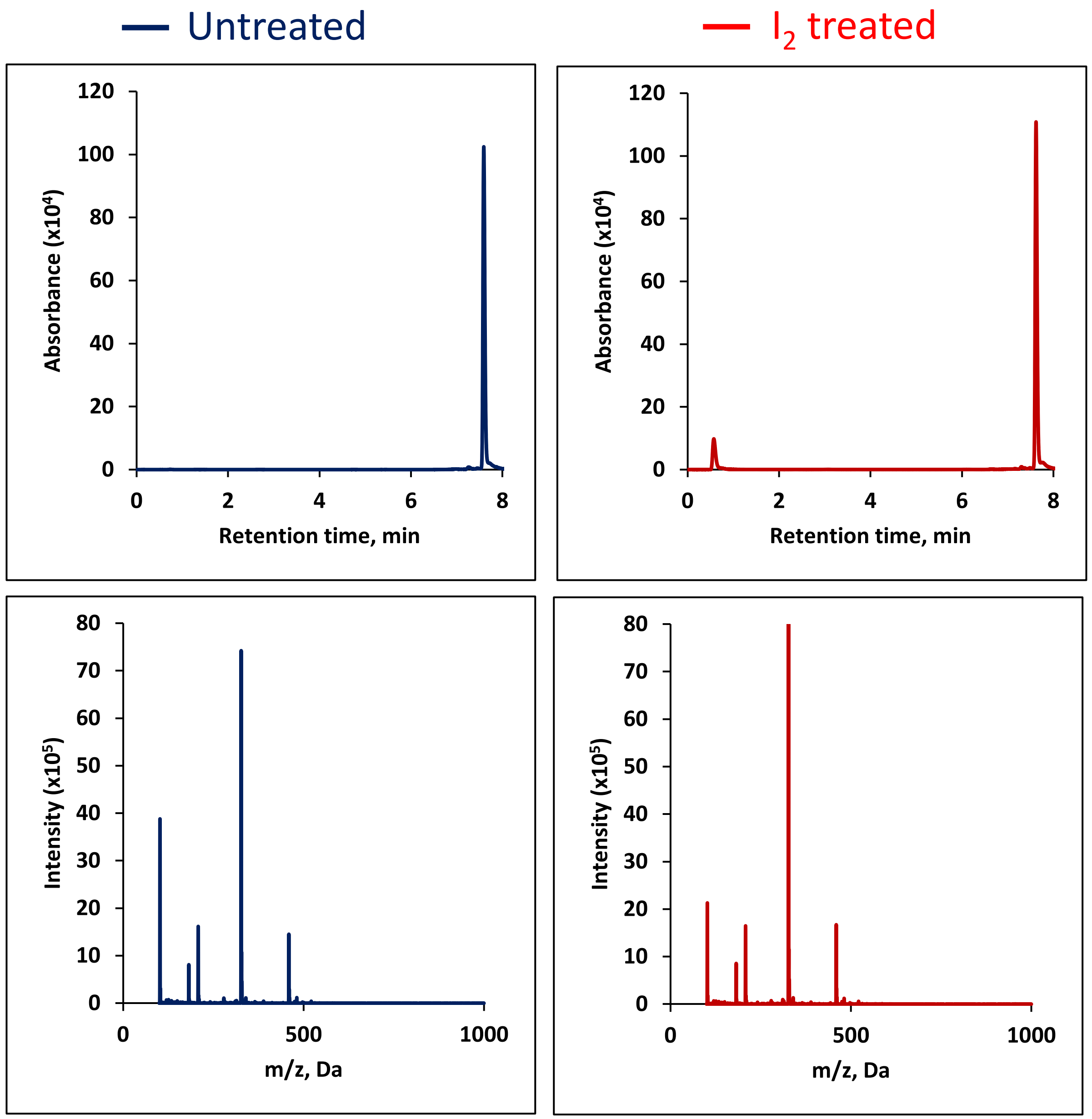

### N6-Threonylcarbamoyl-adenosine (t⁶A)

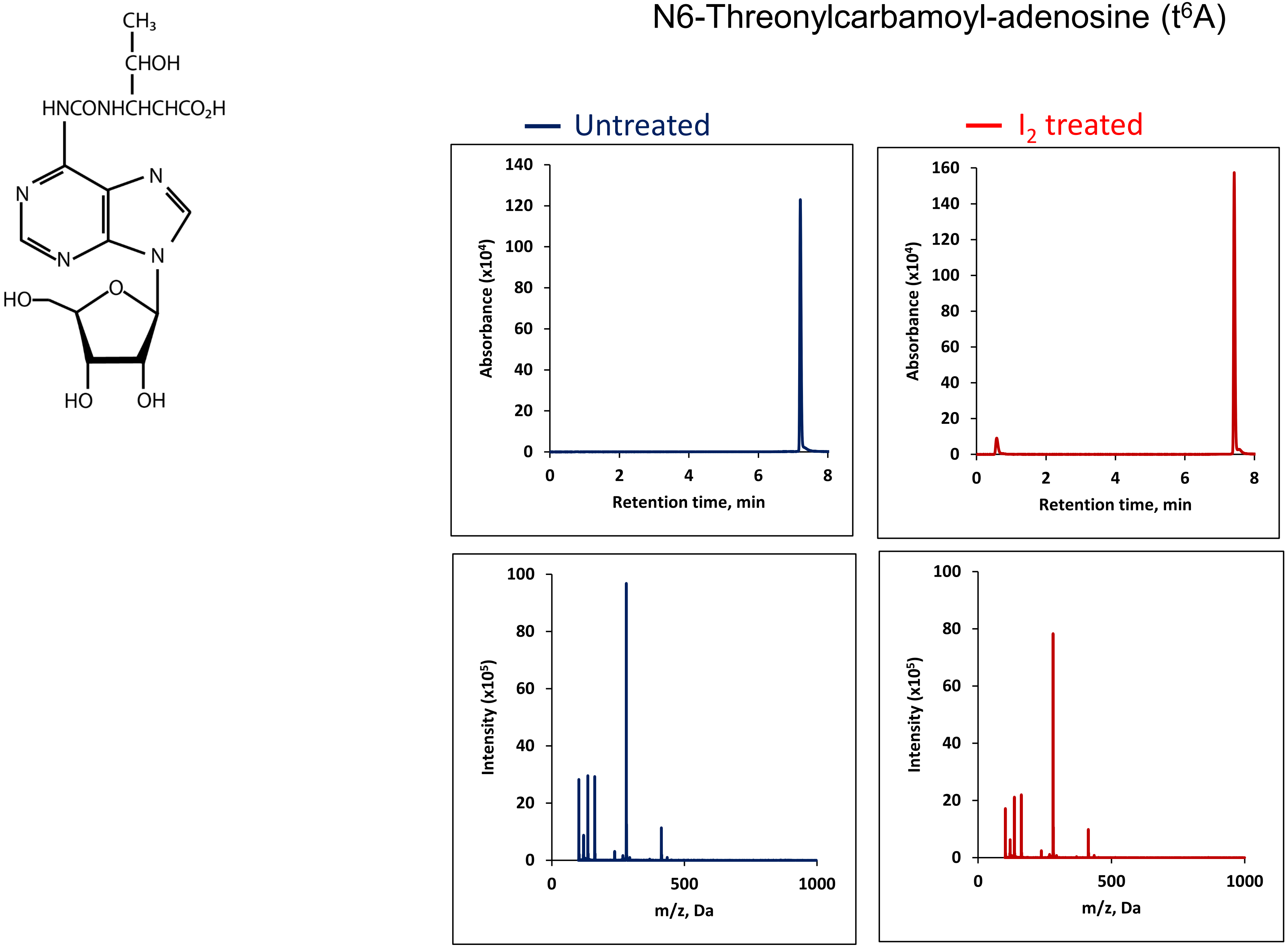

#### $N^4$-methylcytidine ($m^4C$)

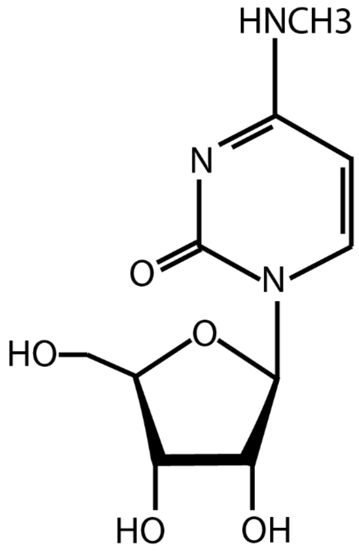


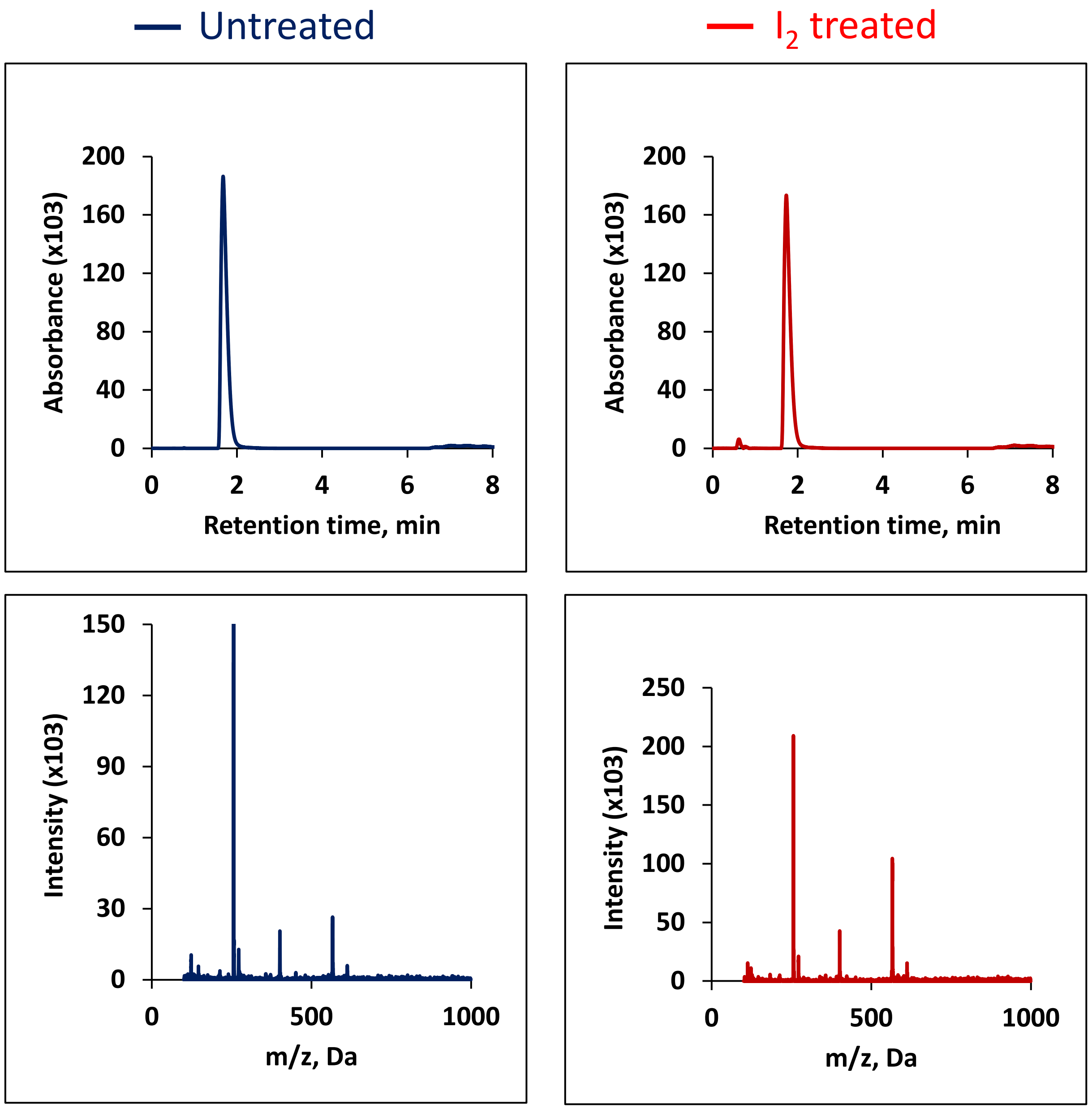

### 5-Methylcytidine ($m^5C$)

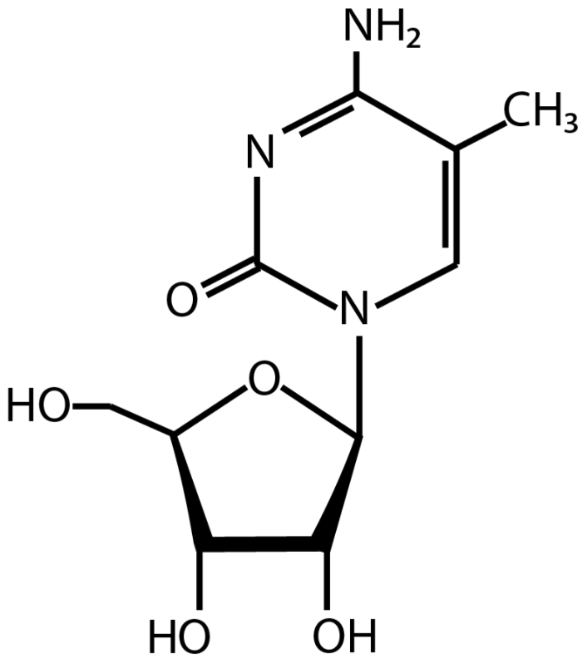


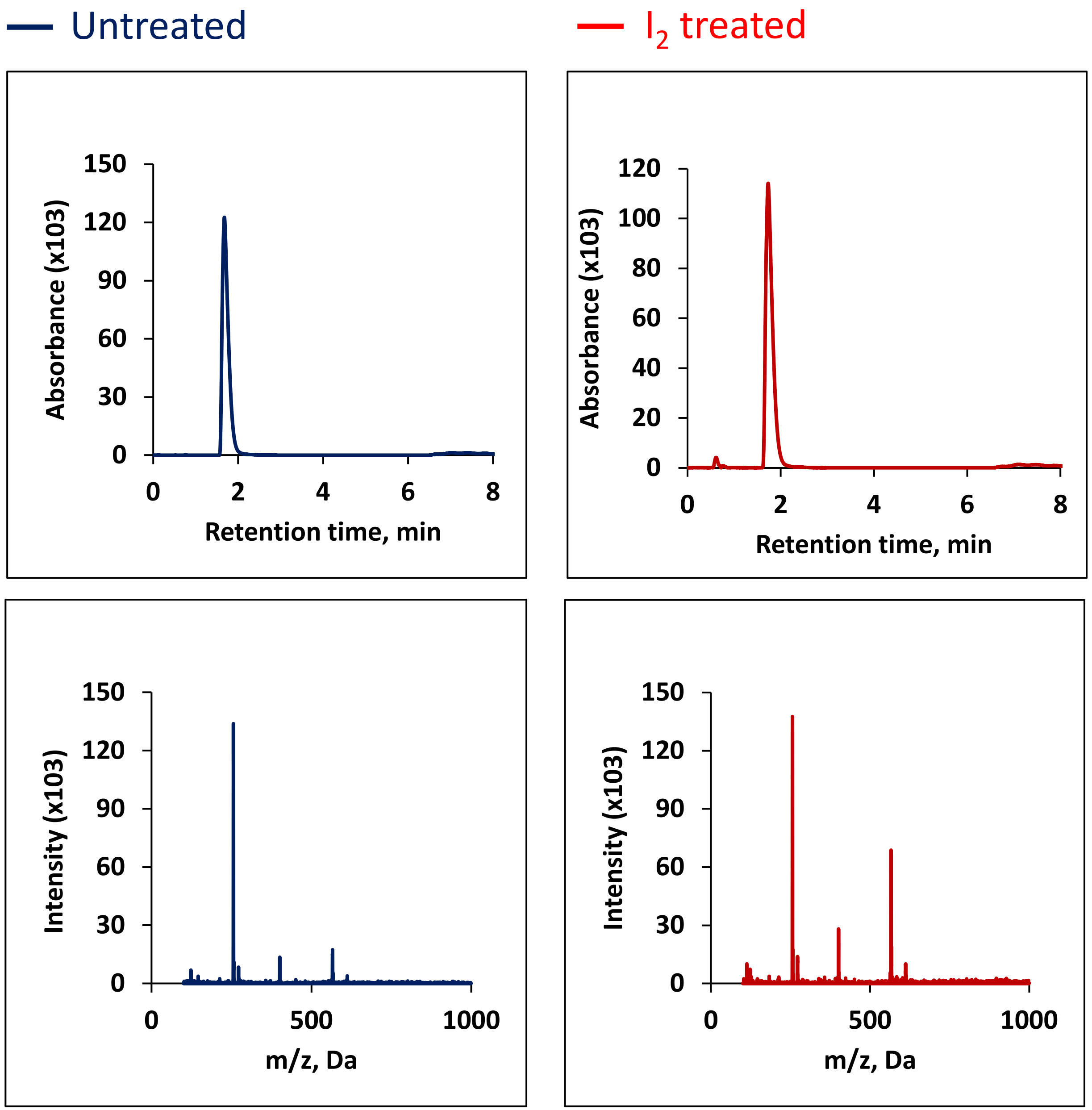

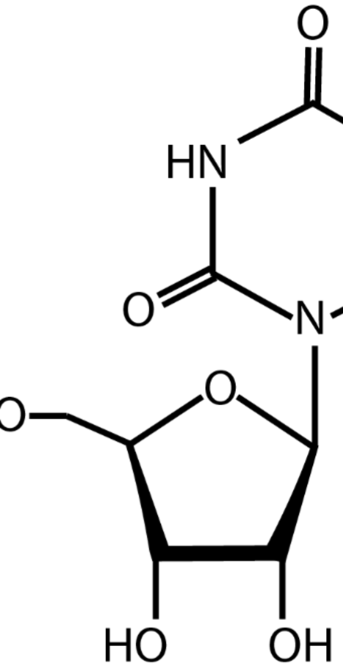


Dihydrouridine (D)

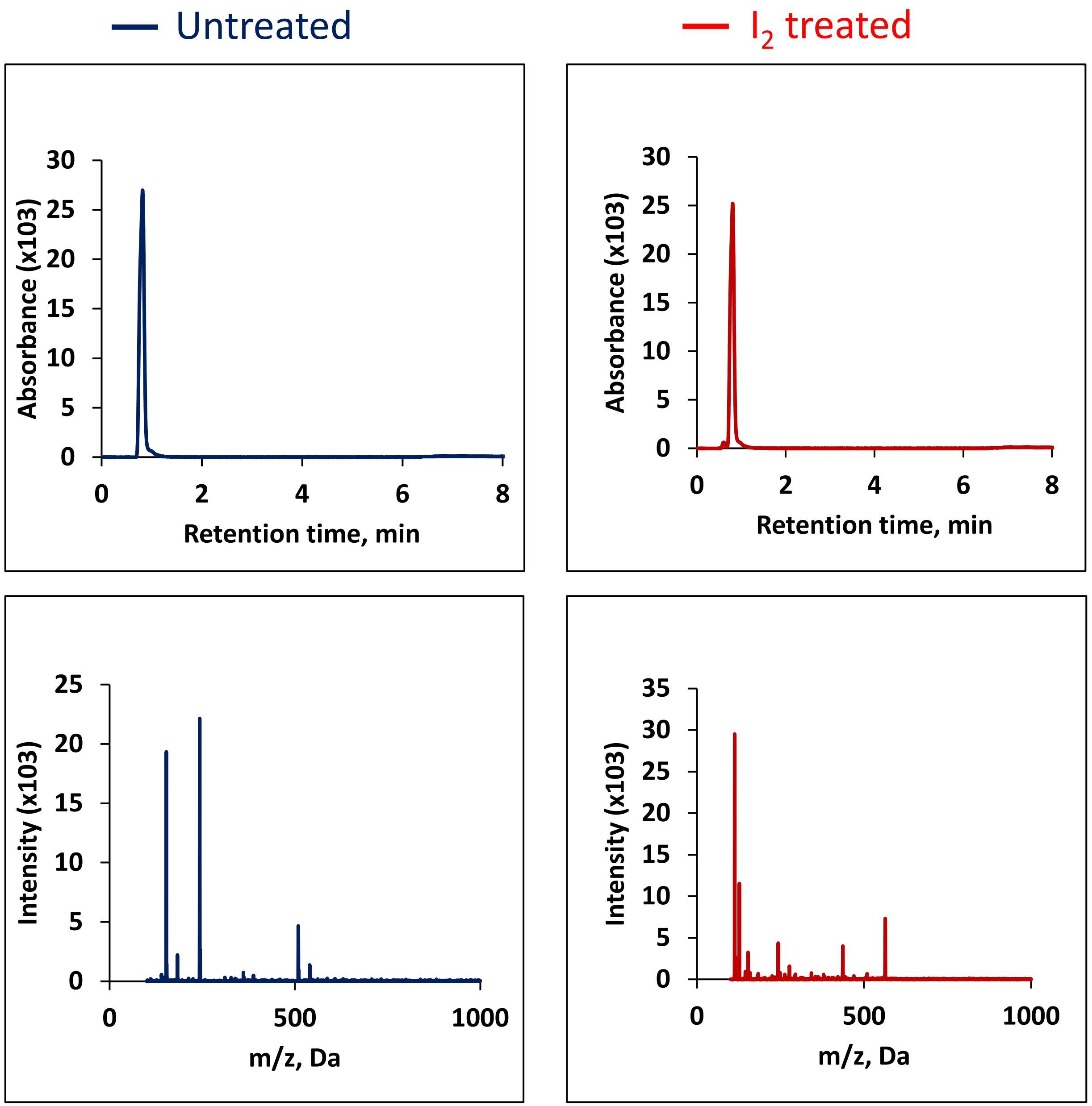

Supplementary Data 2

**Psmd1 mRNA nucleotide mutation rate**

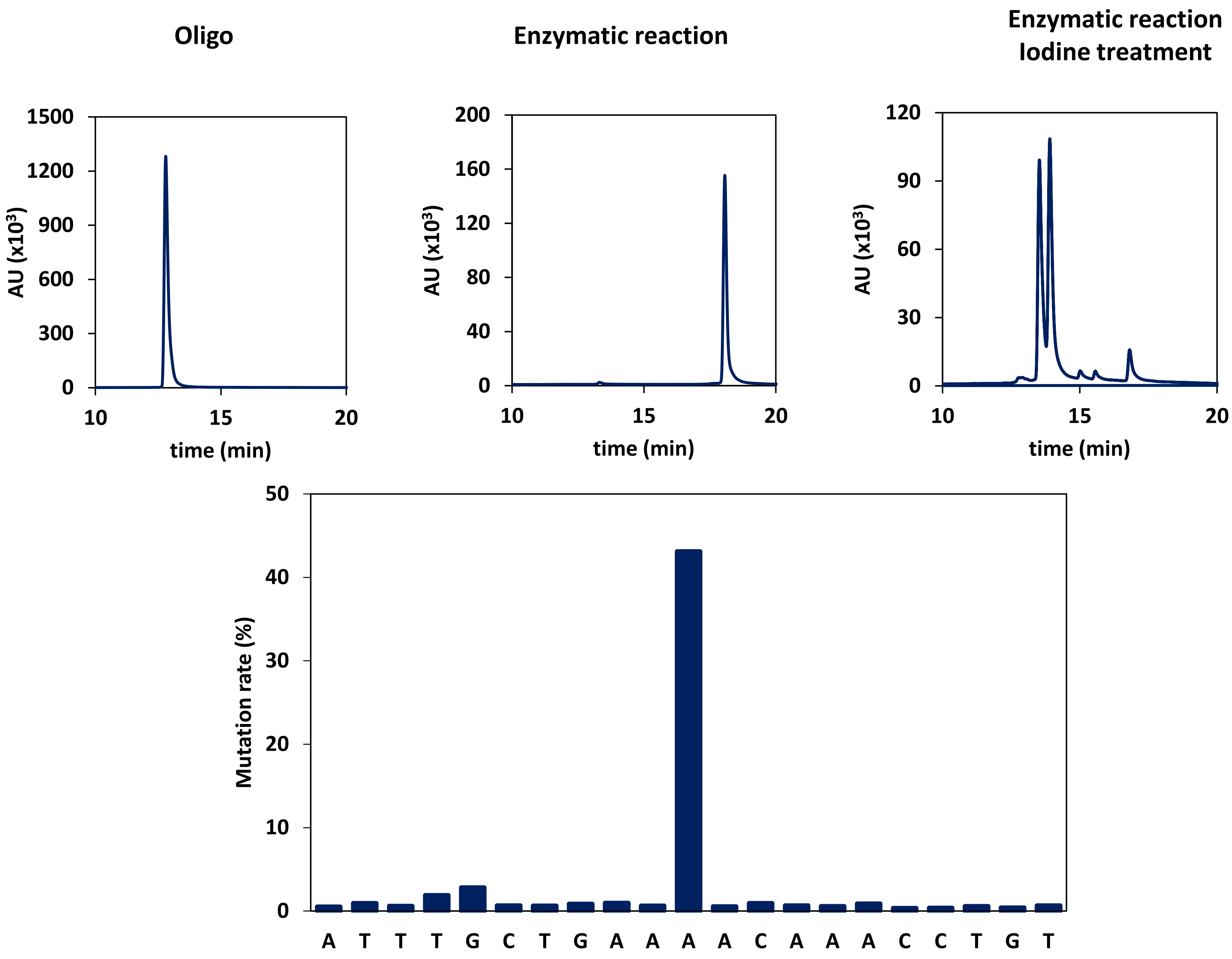

#### C4b mRNA nucleotide mutation rate

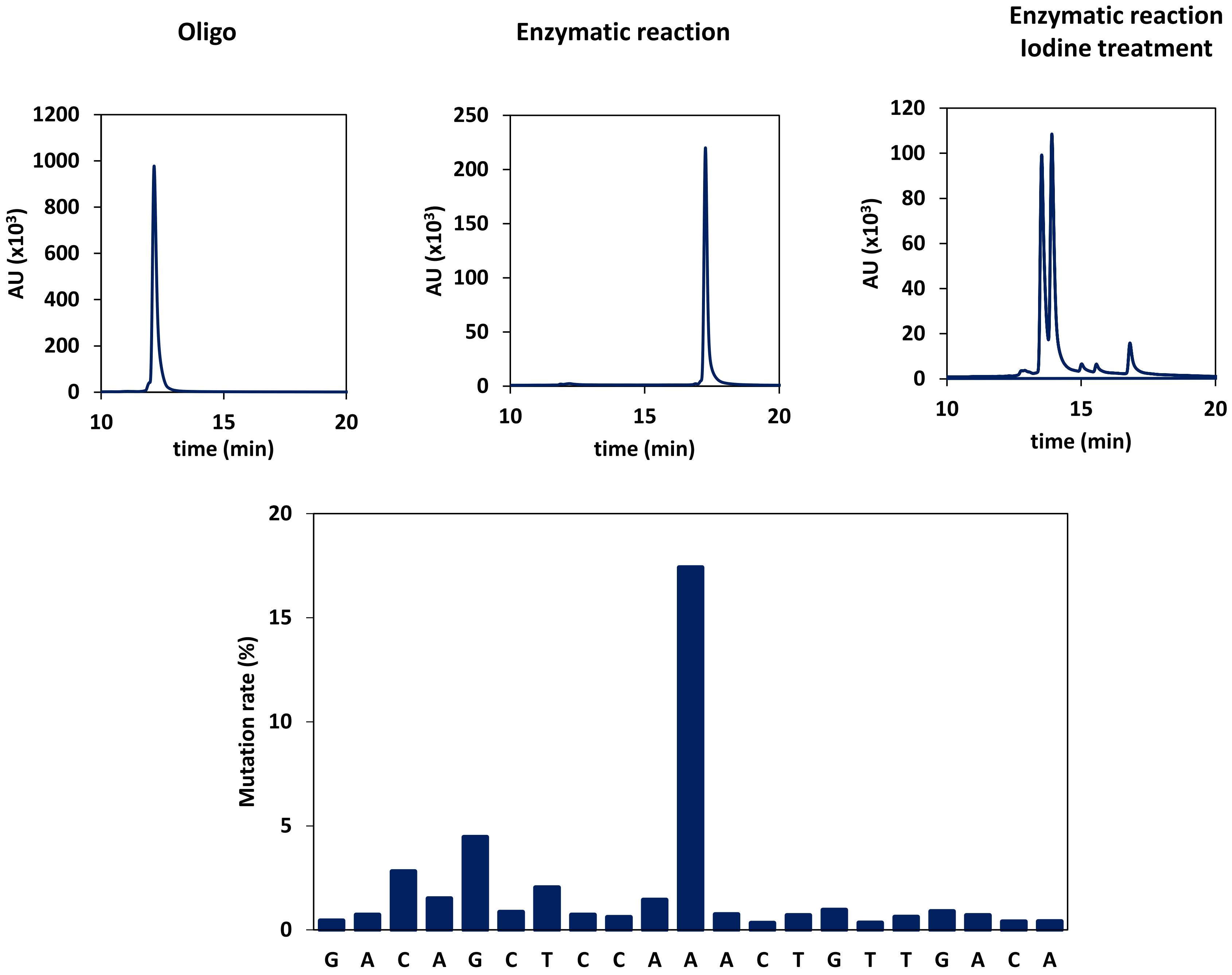

#### DHX9 mRNA nucleotide mutation rate

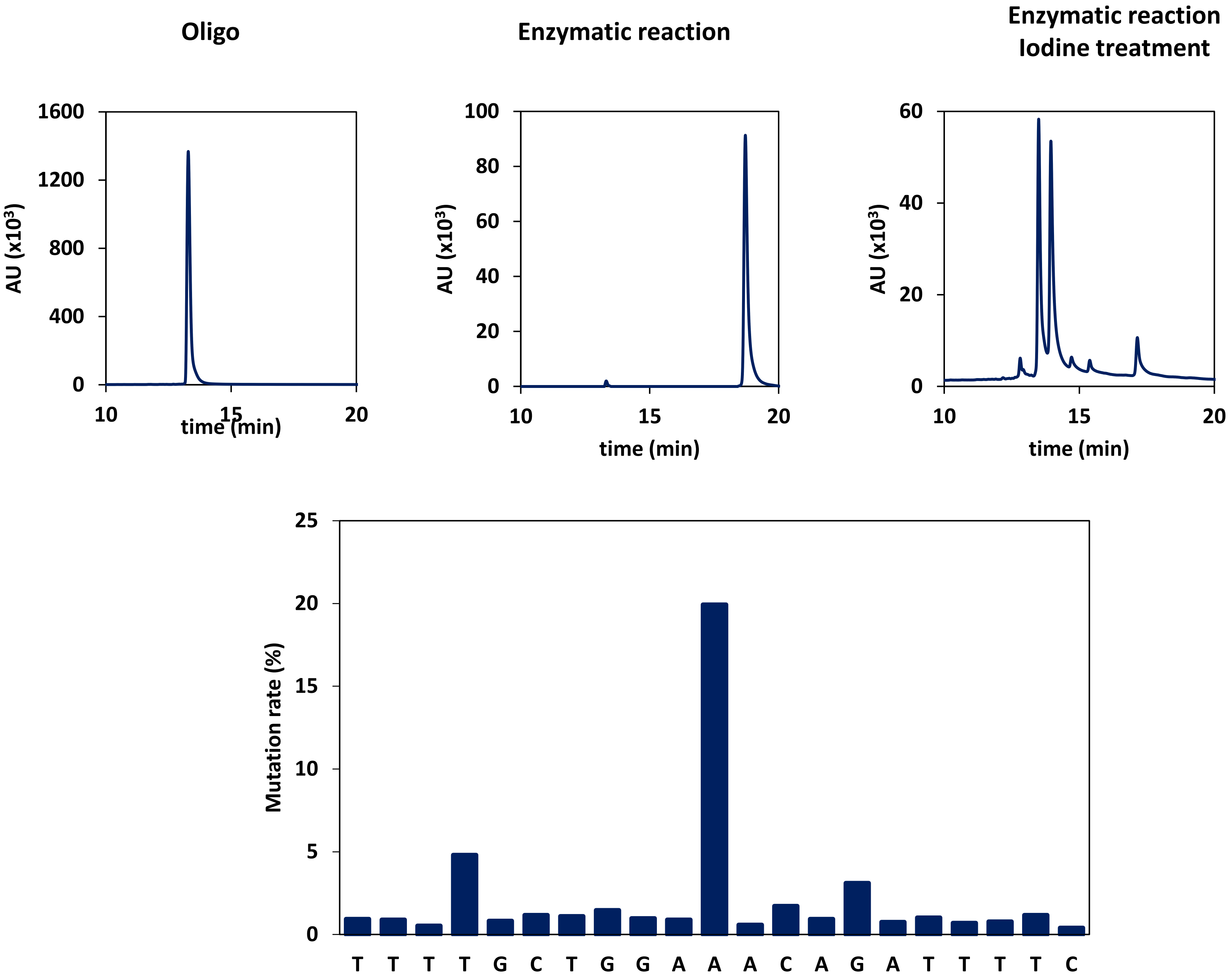

#### CCDC47 mRNA nucleotide mutation rate

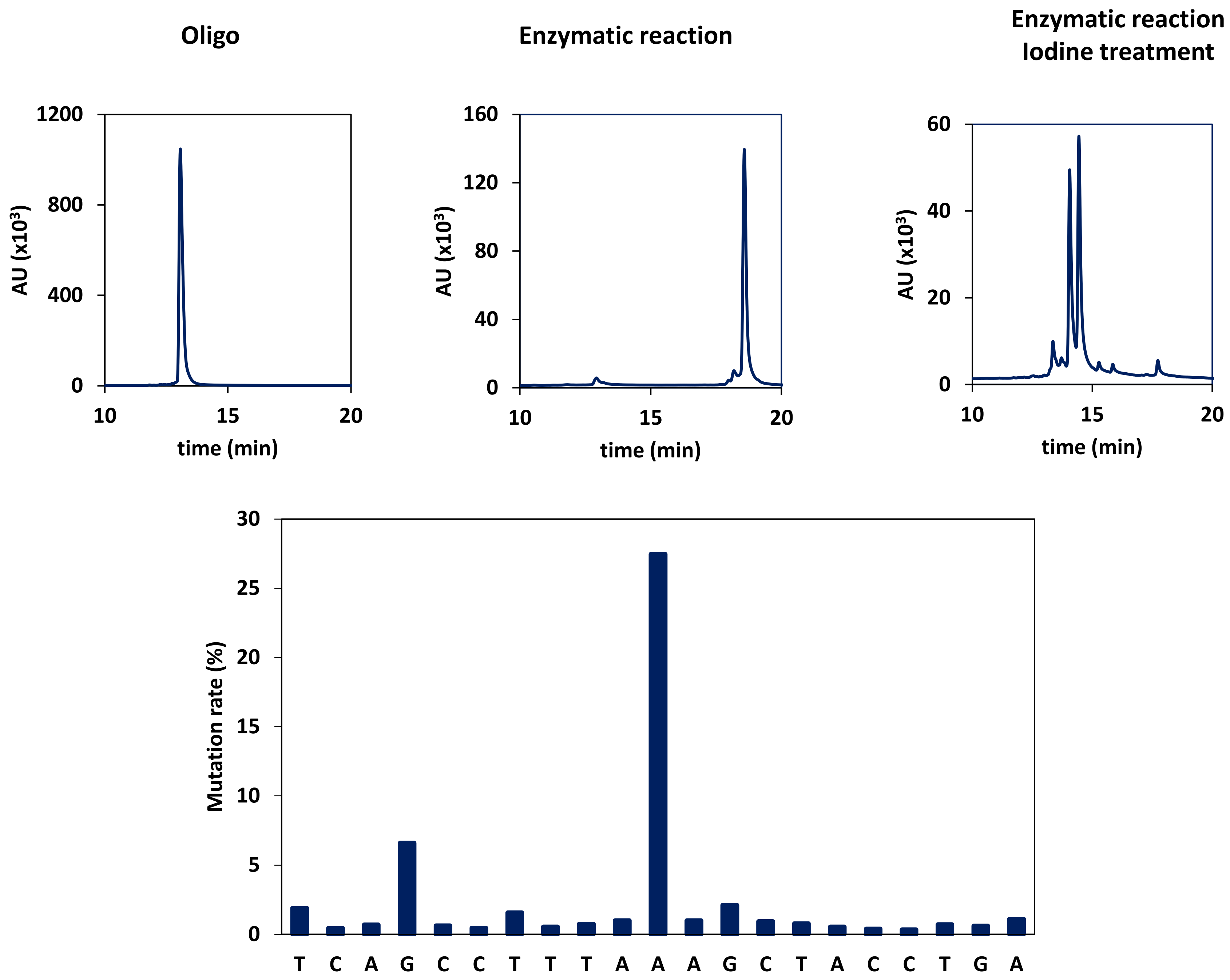

#### ANKRD17 mRNA nucleotide mutation rate

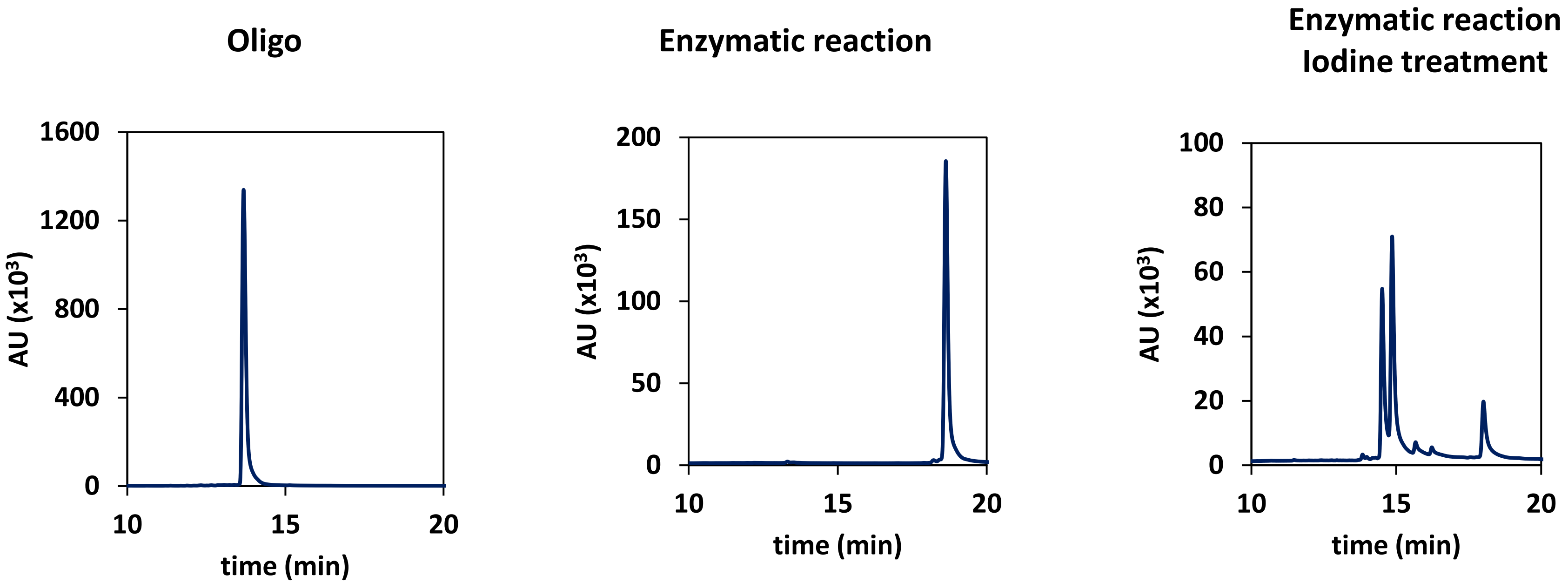


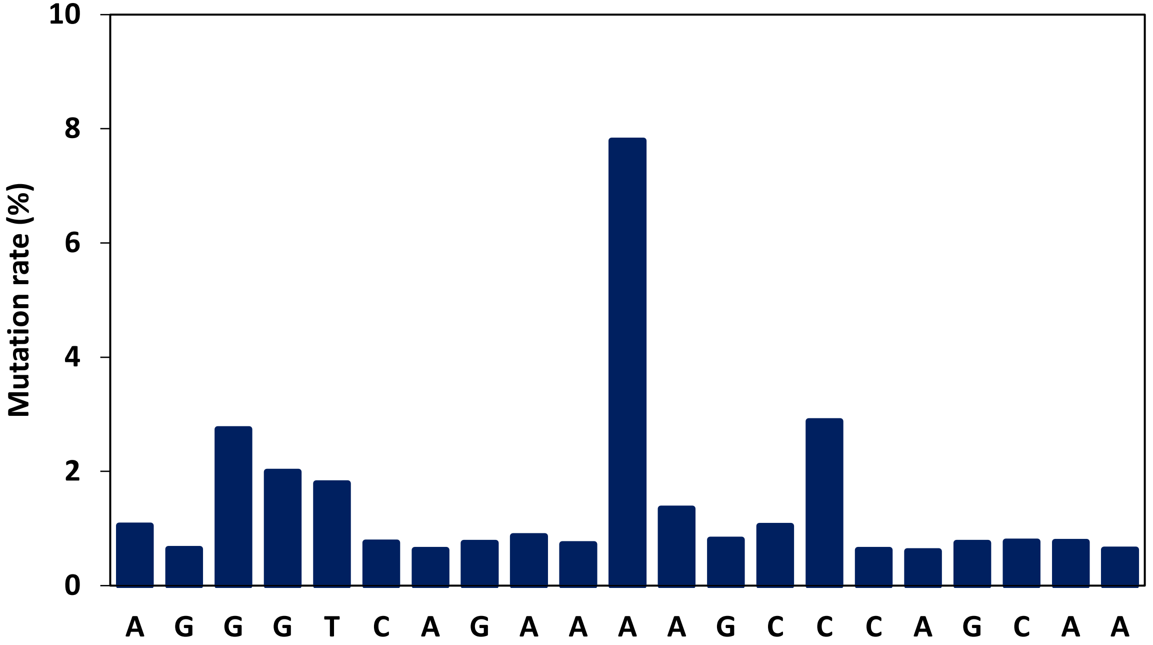

#### SLC16A1 mRNA nucleotide mutation rate

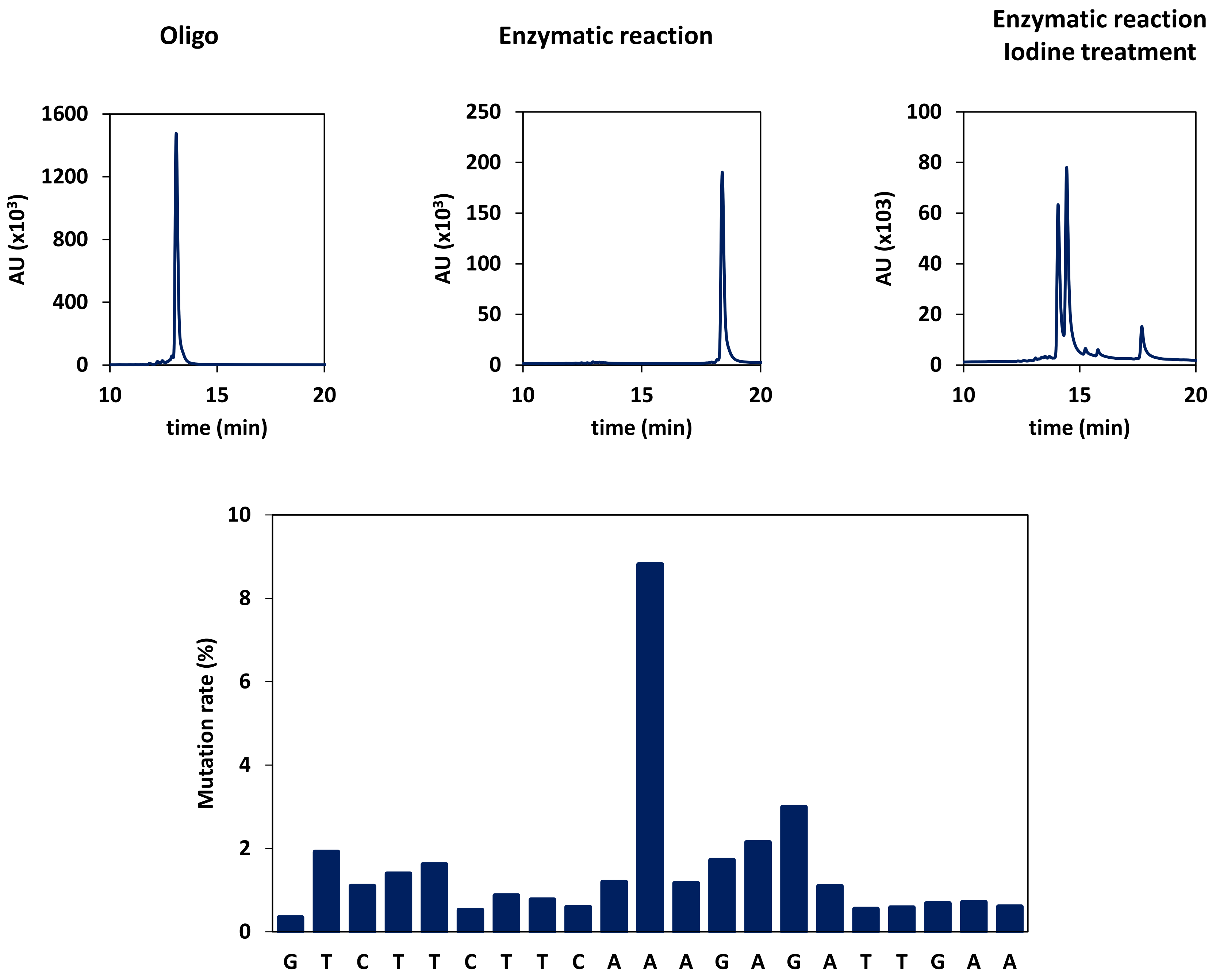
